# Contextualizing the polygenic basis of human brain shape

**DOI:** 10.64898/2026.10.01.754889

**Authors:** Judit Casas-Riu, Cedric Boeckx

**Affiliations:** University of Barcelona; Barcelona, Spain; Radboud University; Nijmegen, The Netherlands; University of Barcelona Institute of Complex Systems; Barcelona, Spain; University of Barcelona Institute of Neurosciences; Barcelona, Spain; Catalan Institute for Research and Advanced Studies (ICREA), Barcelona, Spain

**Keywords:** Globularity, brain evolution, pleiotropy, white matter, ventricle, cerebellum

## Abstract

As the most recently evolved neuroanatomical trait of our species, our globular head shape (‘globularity’) constitutes a promising point of entry to probe possible associations between brain morphology and cognitive traits that may be distinctive of *Homo sapiens*. To shed additional light on the underlying biology of globularity and its cognitive consequences, we capitalize on data pairing genetic variation to brain shape differences from magnetic resonance imaging (MRI) scans of 34,595 adults of European ancestry, and embed genome-wide associations of globularity within genetic variation underpinning up to 75 brain-related traits. Through extensive hypergeometric tests, we observe significant associations between globularity and white matter traits at the gene level, but stronger associations with cerebellar volume, ventricular size, and the fornix at the genomic variant level. Combining conjunctional false discovery rate (FDR) with genetic correlations, we notice homogeneously concordant relations between globularity and the volume of the brain ventricles, and homogeneously discordant relations with glymphatic function. We also unravel more selective, spatially organized relations between globularity and cerebellar volume, white matter connectivity and sulcal width. Spatial permutation tests reveal that sulcal width associations are systematically distributed along a sensorimotor-association gradient of cortical organization. Additionally, correlations of *brain*- and *lifespan*-wise gene expression patterns linked to globularity and the brain ventricles highlight recurrent associations with the pentose phosphate metabolic pathway. Finally, deploying an open repository of genetic variant age estimations and an extensive catalog of single nucleotide changes distinguishing hominins, we uncover a coincidence between the establishment of globularity and the time of emergence of modifications in the cerebrospinal fluid proteome. Overall, we provide a finer-grained genetic architecture for globularity, with novel biological associations to traits across evolutionary and developmental timescales, leading us to draw implications for the early development of cognitive networks but also for liabilities to neurodegeneration late in life.

## Introduction

The globular shape of the neurocranium has long been used as a diagnostic to identify *Homo sapiens* (Stringer, 2016). Indeed, unlike overall head size, ‘globularity’ markedly distinguishes our species from its most closely related extinct relatives. Such a difference emerged relatively late in our lineage (Neubauer, 2018), and is used to distinguish between “basal” and “derived” *sapiens* specimens (Stringer, 2022). Our head shape has been claimed to arise from a *sapiens-*derived pattern of early brain development (Gunz *et al*., 2010; Hublin *et al.,* 2015), the outcome of a perinatal ‘globularization phase’ during which brain growth plays a more determinant role than facial or cranio-basal traits. Virtual brain development reconstruction trajectories point to localized effects of globularization, concentrated at the posterior part of the brain: bulging parietal lobes and enlarged cerebelli (Pereira-Pedro *et al*., 2020; Bruner *et al*., 2023; Kochiyama *et al*., 2018). A reorganization of the parietal lobe is particularly salient from reconstructed endocasts (Bruner, 2004, 2010).

Given its *sapiens*-distinctive nature and the timing of its emergence at a key developmental period for neural connectivity, globularity has been hypothesized to have contributed to neural modifications underpinning behavioral and cognitive traits that are specific to us (Boeckx, 2013, 2017; Boeckx & Benítez-Burraco, 2014a-b; Lombard, 2025a-b; Zeloni *et al*., 2026). Importantly, in light of the ongoing reappraisal of archaic hominins’ cognitive sophistication (Sykes, 2020; Roebroeks & Soressi, 2016; Hoffmann *et al*., 2018), differences linked to globularity are thought to have been subtle, refining key brain structures, circuitry and cognitive capacities from a background of overall similarity between Neanderthals and *Homo sapiens*.

Figuring out what the cognitive impact of globularity may have been requires probing the nature of human brain development at a degree of resolution that goes well beyond what endocast research can provide. As valuable and reliable such signals are (Balzeau *et al.,* 2025), brain microstructure, circuitry and deeply embedded regions cannot be directly gleaned from them. For this reason, researchers have turned to (ancient) genomes and sought to associate *Homo sapiens*-specific mutations to distinct brain growth trajectories. Two studies have taken advantage of the existence of introgressed Neanderthal alleles to probe their impact on the shape of the neurocranium (Gunz *et al*., 2019; Zeloni *et al*., 2026). Recently, a study has instead leveraged *sapiens*-internal variation within contemporary populations (Cambridge Centre for Ageing Neuroscience, UK Biobank) to shed light on the neurobiological bases underlying human endocranial globularity through a genome-wide association study (GWAS). They discovered 77 and 155 significantly related genes and independent single nucleotide polymorphisms (SNPs), respectively (Molz *et al*., 2024). Through genetic correlations, their results point to substantial overlaps with the volume of the brain ventricular system, microstructural measures of white matter tracts, metrics of sulcal width, sagittal brain skew, and ultimately reading and language-related skills.

In this work, we place the results of this Globularity GWAS in a wider context, linking these findings to data from multiple brain-related GWAS from large cohorts (Suppl. Tables 1-3). With our choice of GWAS, we aim not only to extend and refine known relations of globularity with the phenotypes uncovered by Molz *et al*. (2024), but also to broaden our understanding of the structural and functional implications of the globular endocranium for the modern human brain. Thus, our selection of GWAS is further motivated by research highlighting the relevance of evolutionary impacts on the cerebellum (Moberget *et al*., 2026; Barton & Venditti, 2014), higher-order cognition or particular networks of brain activity (Mantini *et al*., 2013; Wei *et al*., 2019), and the timing and vulnerability of brain maturation and aging (Bufill *et al*., 2011; Vickery *et al*., 2024), as well as investigations suggestive of additional phenotypic relations of globularity-associated traits, for instance, with the glymphatic system (Wu *et al*., 2025) or macrostructural parameters of the white matter connectome (Sha *et al*., 2023).

Within our collection of GWAS, extensive hypergeometric tests uncover an asymmetry between gene level associations, where globularity is most strongly linked to white matter traits, and genomic variant level associations, where globularity most closely associates with structures like the cerebellum, the ventricles and the fornix. In light of the evidence that variant-specific information may offer greater trait-specificity (Spence *et al*., 2026), we resort to conjunctional FDR techniques (Andreassen *et al*., 2013) to enhance our identification of variants pleiotropically linked to globularity and other brain-related traits. By assessing the effects of these shared variants, we reveal homogeneously concordant and discordant relations between globularity and brain ventricular volumes and glymphatic clearance, respectively. Further, we unravel spatially organized heterogeneous relations with the cerebellum, white matter connectivity and sulcal width, while providing additional support for some of our findings through genetic correlations. Transcriptomic data helps us uncover relevant spatiotemporal contexts in which variants may exert their pleiotropic effects. Finally, we place our findings in the context of a comprehensive catalog of single nucleotide changes distinguishing modern humans from archaic hominins (Kuhlwilm & Boeckx, 2019) and detect a coincidence in time between globularity and modifications on the cerebrospinal fluid proteome. Cumulatively, using modern within-species variation as a proxy for the evolutionarily relevant phenotype of the globular brain shape, our work recovers and sharpens previously noted neurobiological associations of globularity (e.g., parietal structures, cerebellum, ventricles), while also expanding the significance of this *sapiens*-distinctive trait for the evolution and development of further brain-related phenotypes (e.g., cognitive networks, psychiatric disorders, glymphatic function).

## Results

To understand the genetic architecture of globularity in relation to that of other brain-related traits, we first carried out a literature-based search and selection of GWAS using the NHGRI-EBI GWAS Catalog (accessible at https://www.ebi.ac.uk/gwas/), and Pubmed results (https://pubmed.ncbi.nlm.nih.gov/). Findings obtained by Molz *et al*. (2024) pointing to associations between globularity, the brain ventricles and developmental pressures from the cerebrospinal fluid system led us to include genetic studies for metrics of ventricular volume and structure (e.g., Ge *et al*., 2024; Zhao *et al*., 2022), the cerebrospinal fluid proteome (e.g., Western *et al*., 2024) and metabolome (e.g., Caro *et al*., 2025; Wang *et al*., 2024), idiopathic normal pressure hydrocephalus (Räsänen *et al*., 2024), and the DTI-ALPS (diffusion-tensor imaging analysis along the perivascular space) index of glymphatic function (e.g., Ran *et al*., 2025; Huang *et al*., 2025), which measures the efficiency of exchange dynamics between cerebrospinal and interstitial fluids and has been shown to relate to ventricular size (Wu *et al*., 2025; Li *et al*., 2023). Further postulated phenotypic links of globularity led us to extract data for traits like cerebellar size (Moberget *et al*., 2026), the geometry of subcortical and cerebellar structures (Primus *et al*., 2025), white matter *micro-* (Zhao *et al*., 2021) and *macro-*structure (Sha *et al*., 2023), or sulcal morphology (Sun *et al*., 2022; Lemaitre *et al*., 2023). In an attempt to shed light on the potential implications of globularity for cognitive health and disease, we also included datasets related to language-related white matter connectome microstructure (Mekki *et al*., 2022), psychiatric disorders (e.g., Sha *et al*., 2025), network measures of brain activity (Bell *et al*., 2022), and brain aging (e.g., Kim *et al*., 2026; Wightman *et al*., 2021). (For a full list of all the datasets retrieved, see Suppl. Tables 1-3).

### Asymmetric associations of globularity at the gene and variant levels

We performed pairwise intersections between our datasets to quantify the number of independent significant variants and genes they share (Suppl. Tables 4-5). The significance of the overlaps was assessed via hypergeometric tests (with Benjamini-Hochberg correction), setting the *gene*- and *variant*-level universes to 20,781 and 8,000,000, respectively (Suppl. Tables 6-7). After deriving matrices of negative log10-transformed *p* values, we resorted to the *Louvain* method for community detection. This procedure indicates a low degree of modularity both for *variant*- (0.22) and, especially, *gene*-level (0.13) matrices, suggesting relative continuity and inter-connectedness of our datasets, rather than sharply distinguishable clusters among them.

Our first key finding is that globularity is most substantially linked to *macro*- (Sha *et al*., 2023) (*p* = 3.53E-51) and *micro*-structural (Zhao *et al*., 2021) (*p* = 2.46E-25) measures of white matter connectivity, as well as cerebellar grey matter volume (Moberget *et al*., 2026) (*p* = 4.9E-23), from the perspective of genes (Suppl. Table 6, Fig. 1b-c). By contrast, its most notable overlaps in terms of variants are with cerebellar grey matter volume (Moberget *et al*., 2026) (*p* = 4.5E-36), ventricular morphology (Ge *et al*., 2024) (*p* = 3.66E-20), and fornix white matter microstructure (Ou *et al.,* 2023) (*p* = 3.38E-19) (Fig. 1b,d, Suppl. Table 7). In particular, we observe a set of four *C16orf95* (rs1108048, rs62056161, rs62056167) and *GNA12* (rs1183079) variants exclusively shared between these phenotypes (fornix, cerebellum, ventricles) and globularity. Lower-ranking datasets present asymmetries in their associations with globularity as well. *Significance of overlaps is not directly tied to set length* (Fig. 1b).

**Fig. 1.**
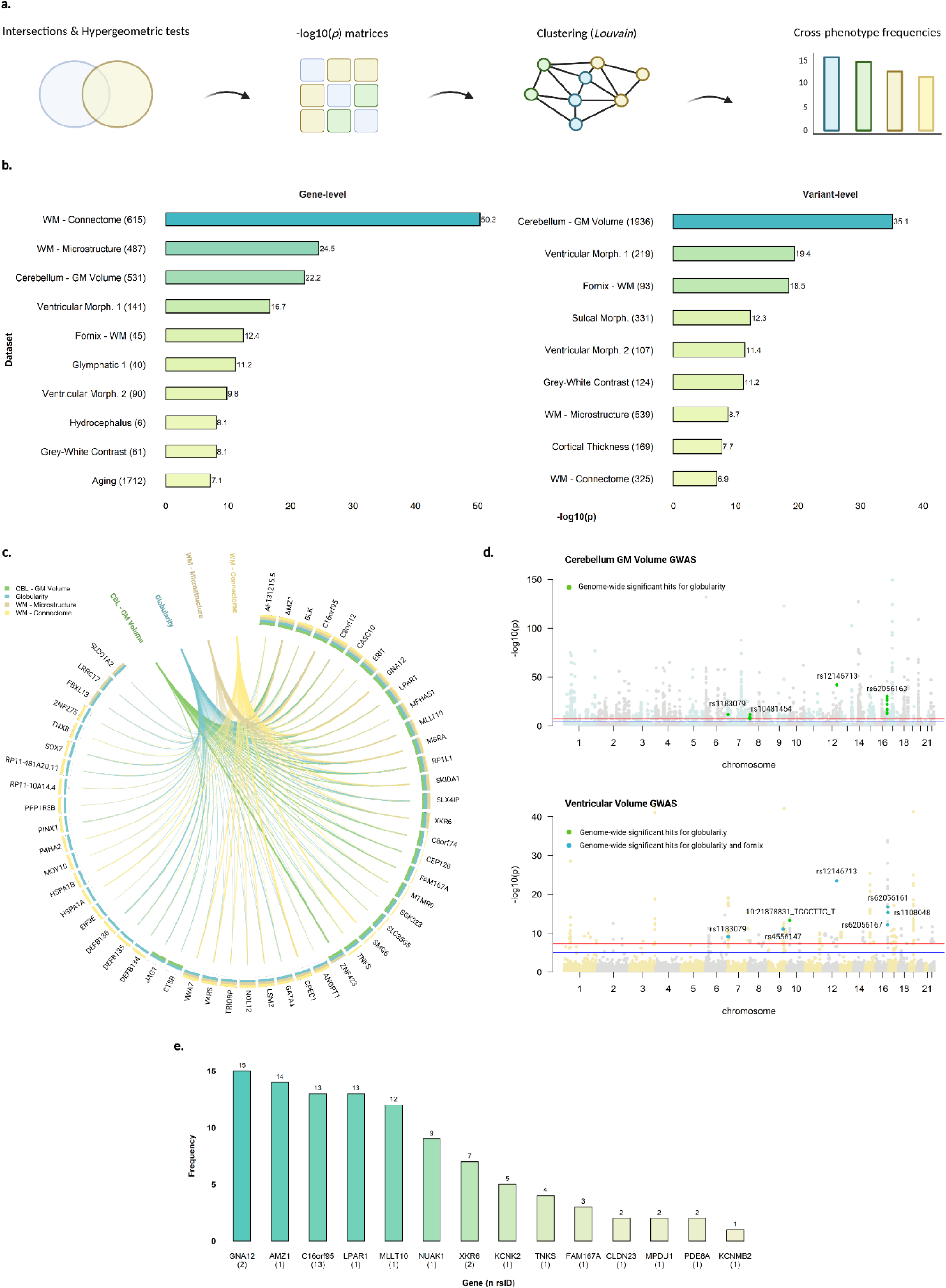
GWAS intersection analysis. **a.** Overview of gene- and variant-level assessments. **b.** Topmost significant phenotypes intersecting with globularity at the levels of genes and variants. Significance is evaluated in terms of negative log10-transformed *p* values (x-axis, Benjamini-Hochberg-corrected) resulting from computing the hypergeometric distribution function of each overlap between globularity and a brain-related trait. Each phenotype label in the y-axis reflects a single GWAS-derived set of genes or variants, with its associated length in parentheses (cf. Suppl. Tables 1-2 for correspondences). **c.** Genome-wide significant genes shared between globularity and three phenotypes from our database (*white matter macrostructural connectome, white matter microstructure, cerebellar grey matter volume*) most closely associated to it at the gene-level. **d.** Manhattan plots for globularity’s closest variant-level correlates in our pool of datasets. Negative log10-transformed *p values* reflect those of the original GWAS (top: *cerebellar grey matter volume*; bottom: *ventricular volume).* Hits that are also genome-wide significant for globularity are marked in green in both Manhattan plots; at the bottom plot, variants additionally significant for the fornix’s white matter microstructure are represented in blue. Annotations indicate the most significant variants intersecting with globularity’s at each chromosome. **e.** Frequency of occurrence, across GWAS in our pool, of genes mapped to variants commonly linked to globularity and at least one other trait in our database. In the x-axis, gene names are shown with the number of globularity-related variants mapped to them in parentheses.

Because GWAS have been claimed to prioritize variants that are trait-specific either by virtue of their context-specific effects on pleiotropic genes or by virtue of their impacts on trait-specific genes (Spence *et al*., 2026), we asked whether the pool of shared variants we obtained might be related to the former (i.e., *context*) or latter (i.e., *gene*) forms of trait-specificity. To address this question, we calculated cross-phenotype frequencies of occurrence of genes and variants in our data (Suppl. Tables 8-9) as a proxy for pleiotropy, and extracted frequencies for genes mapped to globularity-intersecting variants. We noticed that most globularity-intersecting variants are generally linked to genes with high cross-dataset frequencies (e.g., *GNA12, AMZ1, C16orf95, LPAR1;* Fig. 1e), pointing to pleiotropy, and thus context, as the potentially prevailing driving force of trait-specificity for the variant associations reported.

### Globally homogeneous effects

We then sought to improve our chances of discovering variants likely affecting globularity and brain-related traits. To do so, we leveraged overlaps between independent genome-wide significant hits, as well as SNPs resulting from the pairwise application of conjunctional FDR analyses (Andreassen *et al*., 2013) to the full summary statistics (unavailable for globularity) of 50 phenotypes (excluding trait combinations stemming from the same study; cf. Methods, Suppl. Table 3).

Evaluating the effect directions of the resulting shared variants, we first observed a global concordance between the globular endocranium and the volume of the ventricular system, consistent with Molz *et al*.’s (2024) genetic correlations. Namely, across all four ventricles, GWAS-significant variant effects were homogeneously congruent with measures of globularity (100% concordance), signaling a potential connection between this trait and increases in ventricular size. Conjunctional FDR SNPs from the summary statistics of the four ventricles provided further evidence for this conclusion (Suppl. Table 10, Fig. 2b), with variants mapped to *C16orf95* (rs11860417, rs4843549, rs11641115) and *SLCO1A2* (rs10841789).

**Fig. 2.**
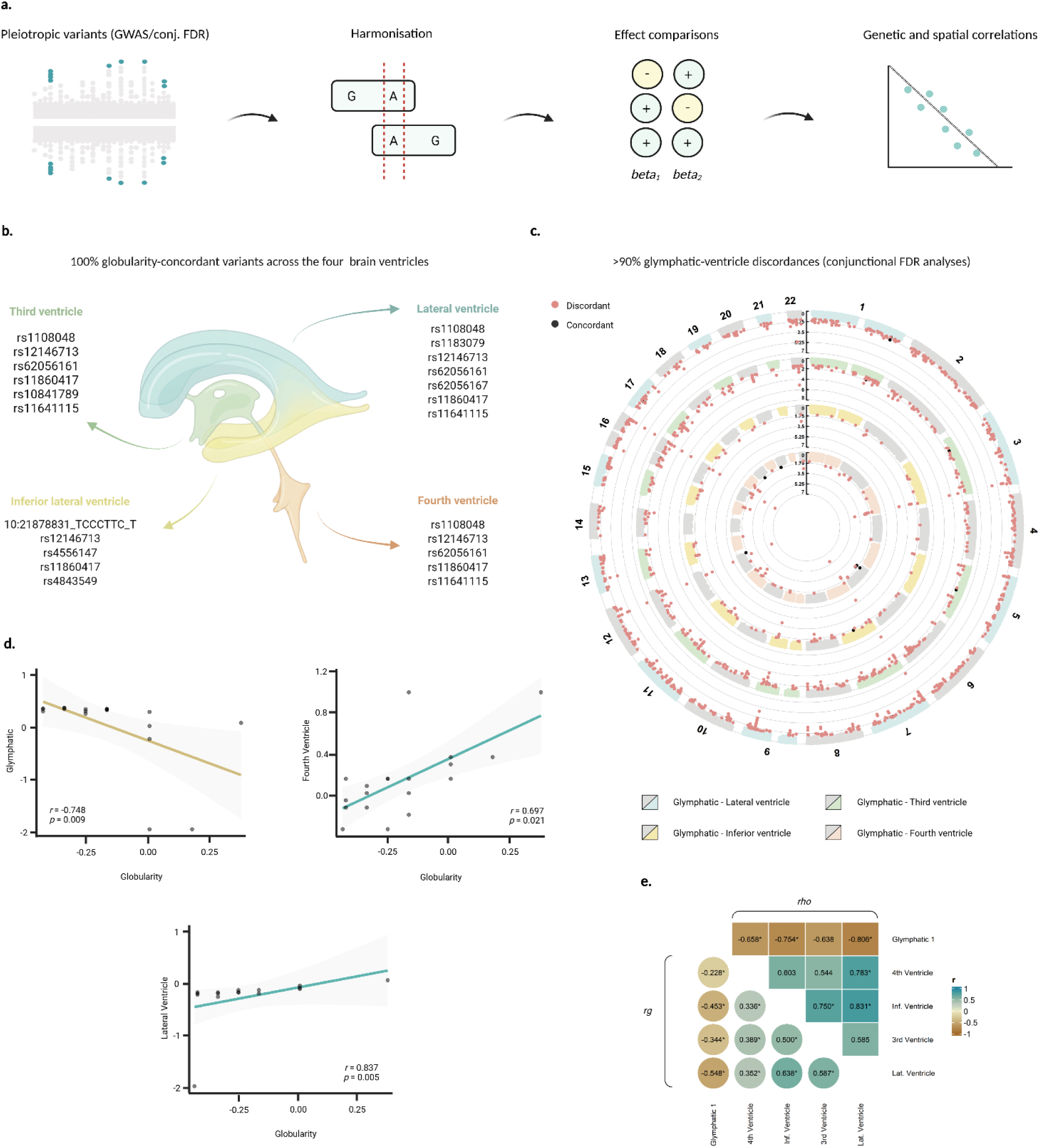
Globally homogeneous effects. **a.** Overview of analyses of *variant-level* effects. **b.** GWAS- and conjunctional FDR-significant variants showing fully concordant effect directions between each of the brain ventricles and globularity. **c.** Discordant (red) and concordant (black) pleiotropic variants for the glymphatic system and each of the brain ventricles, derived via conjunctional FDR. **d.** Significant correlations of globularity with the fourth and lateral ventricles as well as the glymphatic system, according to their patterns of concordance and discordance with other traits in our database. Values are scaled and *winsorized* for the purpose of visualization. **e.** Correlations between the glymphatic system and each of the brain ventricles. The upper triangle represents *Spearman* correlations based on these phenotypes’ degrees of concordance and discordance with other traits in our pool. The lower triangle depicts genetic correlations. Only relations that remain significant after Benjamini-Hochberg correction are marked with an asterisk (*).

Moreover, globularity and ventricular volume measures tended to remain concordant in their patterns of association with other traits (Fig. 2d, Suppl. Table 13). Specifically, applying Spearman correlations to a quantification of their degrees of convergence and divergence with each of 24 other phenotypes in our effect analysis (*n concordant variants – n discordant variants;* cf. Methods), we saw that globularity is positively and non-trivially correlated with the fourth (*r* = 0.697, *p* = 0.0009) and lateral (*r* = 0.837, *p* = 5.35E-05) ventricles, as well as nominally significantly correlated with the inferior lateral ventricle (*r* = 0.589, unadjusted *p* = 0.0128). The ventricles themselves were either nominally or significantly positively correlated between each other (Fig. 2e, Suppl. Table 13).

The literature linking ventricular enlargements to glymphatic dysfunction (e.g., Wu et al., 2025; Li et al., 2023) prompted us to examine relations between the ventricles, globularity, and one of the glymphatic function correlates (DTI-ALPS; Huang *et al*., 2025) in our GWAS pool. Though scarce, data for globularity hinted at discordances between this phenotype and glymphatic fluid dynamics. More concretely, rs12146713 (*NUAK1*; T > C), a GWAS-significant variant for both globularity and the DTI-ALPS index, showed positive effect sizes for the former trait (*beta* = 1.0173), but negative ones for the latter (*beta* = −0.091). In addition, lead SNPs for globularity (rs62056161, rs11860417, rs4843539, rs10841789) that emerged as enriched for the glymphatic system via conjunctional FDR also displayed inverse impacts (Suppl. Table 11).

Beyond revealing extensive pleiotropic enrichments between ventricular morphology and the DTI-ALPS index (e.g., 806 independent significant signals associated with the lateral ventricle; cf. Suppl. Table 12), conjunctional FDR uncovered robustly discordant effect directions for the volumes of all four ventricles with respect to our glymphatic function measure: across 1283 unique SNPs significantly associated with the ventricular and glymphatic datasets, 98.29% were divergent in their effects (Fig. 2c, Suppl. Table 12), confirming patterns already revealed by GWAS-significant variants at the intersection between ventricular size and glymphatic function (Suppl. Table 12). Genetic correlations added support for the inverse relation between glymphatic function and ventricular volume measures (Fig. 2e, Suppl. Table 14), whereas Spearman correlations indicated that the glymphatic system’s discordance with globularity and the brain ventricles holds consistently across their patterns of association to other brain-related phenotypes (Fig. 2d,e; Suppl. Table 13).

### Selective homogeneous effects

We next examined effect relations between globularity, ventricular volumes, and individual scores attributed to each of 23 distinct regional components studied in a recent GWAS on cerebellar size (Moberget *et al*., 2026), together with a mean score summarizing global effects on the volume of the cerebellum across components. Focusing on those components where genome-wide significant variants for all four ventricles and globularity seemed to be coordinated in their effect relations, we detected systematically convergent (e.g., 03, 08, 14) and steadily divergent (e.g., 11, 15, 19) components (Suppl. Figure 1). After adding conjunctional FDR-significant variants to our inspection, component 03 remained robustly concordant with globularity and ventricular effects, whereas component 02 increased its concordance (both 11/12 = 91.67% variants; Fig. 3a, Suppl. Table 15). Consistent with the predominance of variants related to the fourth ventricle (7/8) in our conjunctional FDR-derived overlaps with cerebellar grey matter volume, these two components displayed significantly positive genetic correlations only with the fourth ventricle (*r* = 0.21-0.26; *p* = 0.004-0.00002; Fig. 3b, Suppl. Table 14), but not the other ventricles, pointing to tighter connections between the size of this ventricle and the volume of cerebellar regions spanning portions of Crus II and lobules VIIb/VIIIa (components 02-03).

**Fig. 3.**
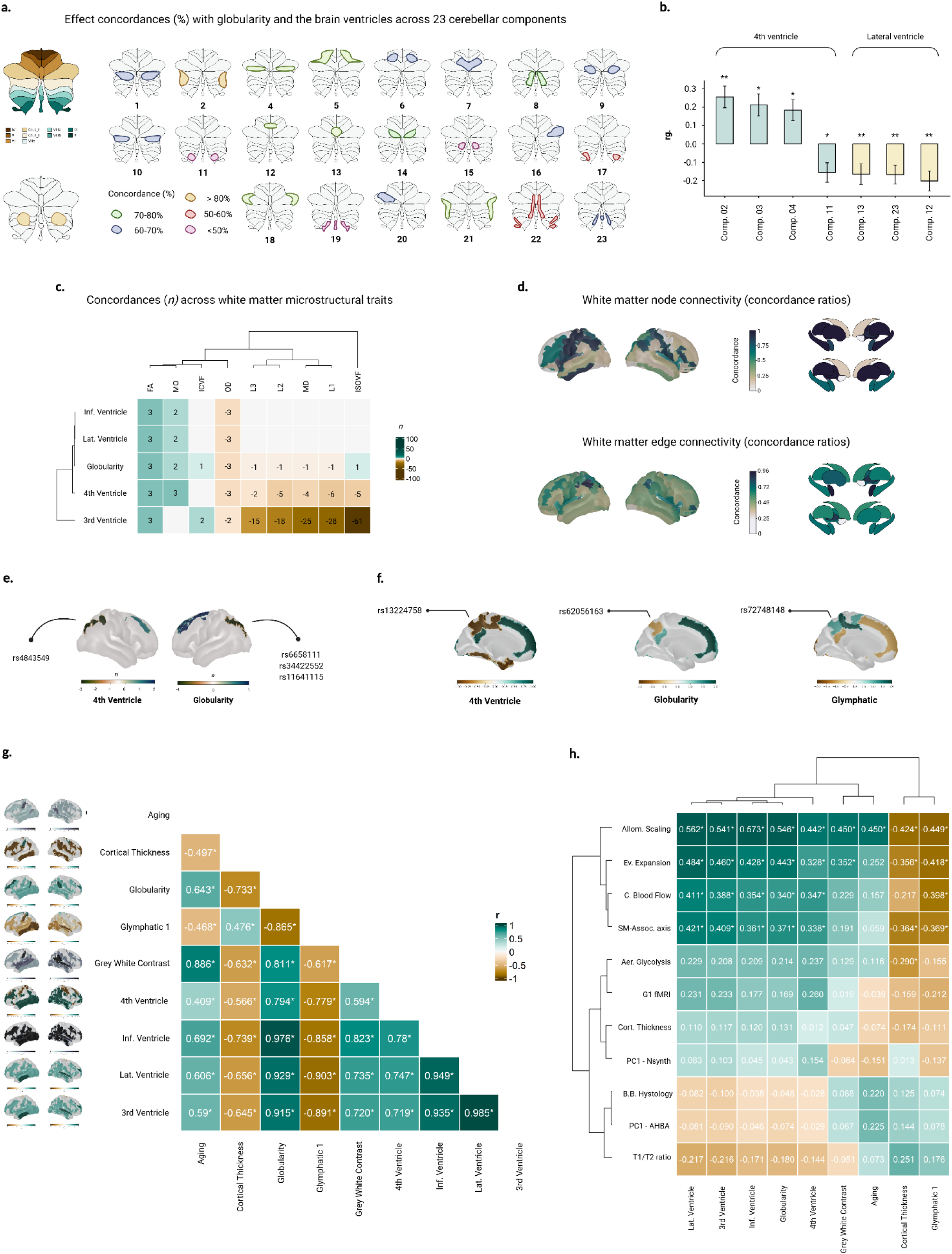
Selective homogeneous effects. **a.** Distributions of effect concordances (%) with globularity and the four brain ventricles across 23 cerebellar components. Percentages were computed from GWAS- and conjunctional FDR-significant variants. **b.** Significant genetic correlations between the brain ventricles and 23 cerebellar components after Benjamini-Hochberg correction. **c.** Counts (*n*) of concordant and discordant variants with globularity and the brain ventricles separate white matter microstructural traits into an architectural cluster and a diffusivity cluster. **d.** Distribution of concordance ratios with globularity and the brain ventricles accross brain regions, according to their corresponding parameters of white matter *node* (top) and e*dge* (bottom) connectivity. **e.** Globularity- and ventricular volume-related variants exhibit concurrent selective impacts on intraparietal depth. **f.** Discordances between globularity, ventricular volume-, and sulcal width-related SNPs specifically target the internal parietal sulcus. **g.** Pairwise correlations between traits according to their associated variants’ spatial patterns of concordance with sulcal width effects. **h.** Spatial correlations for the distribution of sulcal width concordances linked to each of nine traits with respect to eleven sensorimotor-association axis (Sydnor e*t al.*, 2021) features.

Turning to white matter traits, we observed more circumscribed concordant effects similar to what we found for the cerebellum. In particular, microstructural information gathered for seven language-relevant tracts from Mekki (2022) (cf. Methods) revealed concurrent impacts on metrics of white matter integrity (fractional anisotropy, FA), neurite density (intracellular volume fraction, ICVF) and tract directionality (mode of anisotropy, MO), as opposed to dissonant effects on measures of average (mean diffusivity, MD), axial (first DTI eigenvalue, L1) and radial (second DTI eigenvalue, L2; third DTI eigenvalue, L3) diffusivity. Indices of freely diffusing water movement (isotropic volume fraction; ISOVF) and fiber orientation dispersion (OD) also displayed discordant directions of impact (Suppl. Table 16; Fig. 3c). Furthermore, assessing effect relations by tract (Suppl. Fig. 2), we noted that concordant measures (FA, MO, ICVF) held most strongly for the frontal aslant and fronto-insular tracts, which also concentrated the observed discordances with OD. Instead, the corpus callosum (MO, OD), the left hemisphere’s frontal superior longitudinal tract (MO) and the right superior longitudinal fasciculus (ICVF) were more prone to exceptional or weaker correspondences.

To attain a more complete perspective on white matter effect associations, we resorted to *macrostructural* measures of *edge-* and *node-*level connectivity (Sha *et al*., 2023). *Edge-*level connectivity reflects the number of streamlines connecting a pair of regions adjusted by the combined volume of the two given areas, whereas *node*-level connectivity is quantified as the sum, for a certain region, of all the existing volume-weighted edges linking it to other nodes. For each of the 851 edges and 90 nodes under research, we averaged congruency scores across variants, by dataset (*globularity,* as well as *fourth, third, lateral* and *inferior ventricles*). Subsequently, we computed the mean concordance of each node across the phenotypes considered, so as to extract *node-* level congruencies that hold most strongly across globularity and its related traits. To this, we added an *edge-*level consistency score, calculated in terms of the average concordance exhibited by a node based on the degree of convergence of its associated edges.

For *edge-*based congruencies, we found that almost half (47.59%) of the inspected region connections bear concordance rates over 80% (Suppl. Table 17). Among these, we noticed that nodes such as the left Heschl gyrus (95.56%), the right amygdala (93.81%), the parietal inferior gyri (92%) or the right pallidum (84.44%) show the greatest proportions of concordant edges, suggesting a link between increased density of streamlines in a substantial part of their connections to other regions and positive effects on globularity and ventricular size (Suppl. Table 17, Fig. 3d). Regarding *node-*level effects, concordant directions were found in subcortical and temporal structures like the right hippocampus, the left pallidum, the left para-hippocampal, the left and right putamen, and the left and right thalamus, which exhibit 100% rates of convergence with globularity and ventricular effects. Previously highlighted nodes, such as the left Heschl gyrus, maintained relatively high rates of concordance (83.33%), whereas formerly unnoticed nodes emerged with equal (e.g., left superior temporal pole) or higher (e.g., left postcentral and precentral gyri, left inferior temporal gyrus) congruity scores (Suppl. Table 18; Fig. 3d).

Since variations in the organization of white matter *micro-* and *macro-*structural architectures have been linked to the remodeling and formation of brain sulci along specific morphological characteristics (e.g., Roll, 2024; Shan *et al*., 2026), we followed by extending our *variant-level* analyses to effect relations between globularity- or ventricular volume-related SNPs and parameters of sulcal depth and width. Our depth inspection was mainly restricted to independent GWAS- or conjunctional FDR-significant SNPs for globularity and ventricular volumes in overlap with conjunctional FDR-significant variants from the summary statistics of 22 sulci supplied by Lemaitre *et al*. (2023). To these, we added sulci with associated GWAS-significant variants from Sun *et al*. (2022), from which we also extracted data for our evaluation of sulcal width.

Exploring effect associations between globularity and sulcal depth, we unraveled generally sparse relations across most sulci: as captured through spatial permutation tests (cf. Methods), neither the brain ventricles nor globularity converged on a common, correlated spatial distribution of concordances or discordances with sulcal depth (Suppl. Table 19). Despite this global variability, consistently discordant effects were found for the (right) intraparietal sulcus, with most implicated variants linked to *C16orf95* (Suppl. Table 19, Fig. 3e). This sulcus yields nominally significant, negative genetic correlations with the brain ventricles (concretely, third: *r* =-0.13, unadjusted *p* = 0.005), and a nominally significant, positive genetic correlation with glymphatic function (*r* = 0.118, *p* = 0.01) (Suppl. Table 14).

As for variants influencing sulcal width (Sun *et al*., 2022), concordance proportions initially indicated them to be relatively highly congruent (60-100%) in their effect directions with globularity and ventricular volumes (Suppl. Table 20). However, convergent and divergent effect sizes coexisting within a common variant or across SNPs related to the same trait suggested heterogeneity to be explained. Therefore, to understand whether such variability is meaningfully organized throughout the brain, we quantified each sulcus’ degree of congruence with globularity and the brain ventricles according to the effects of its associated variants (*n concordant – n discordant SNPs*), and situated the derived scores in the brain using the BrainVISA sulcal atlas (Perrot *et al*., 2011). Applying *spin* tests to the resulting brain maps, we confirmed that the spatial distribution of convergent and divergent impacts is highly positively and non-trivially correlated between all of the ventricles and globularity (Fig. 3g; Suppl. Table 23).

### Distribution of sulcal width concordances

Positive correlations hinted at a potential coherence between the spatial patterns of overlap with sulcal width presented by globularity- and ventricular volume-related variants. Hence, we sought to further define the spatial distribution of such genetic associations with sulcal width, trying to determine whether they are organized along established anatomical, functional or evolutionary hierarchies of cortical organization. With that aim, we resorted to Sydnor *et al*.’s (2021) *sensorimotor-association archetypal axis* (SMA). This axis has been proposed to capture the spatiotemporal patterning of cortical maturation throughout humans’ prolonged brain development, characterizing the organizational gradients —from lower-order, primary, unimodal cortices to higher-order, transmodal, association cortices— alongside which such developmental program unfolds.

Through spatial correlations under 10,000 *spin* tests between our maps of globularity- or ventricle volume-related concordances with sulcal width and the eleven components of the sensorimotor-association axis, we obtained positive, significant correlations with three associatively-oriented SMA features (Fig. 3h): *allometric scaling* (*r* = 0.44 – 0.57, *p_spin_* = 0.0018-0.0054), which reflects the relative extent of areal scaling with scaling of total brain size (Reardon *et al*., 2018); *cerebral blood flow* (*r* = 0.34 – 0.41, *p_spin_* = 0.01 – 0.04), manifesting rates of blood delivery to the brain quantified through arterial spin labelling (Satterthwaite *et al*., 2014); and *evolutionary expansion* (*r* = 0.32 – 0.48, *p_spin_* = 0.032 – 0.005), which represents the most evolutionarily expanded areas of the human cortex as inferred by comparing human and macaque monkey cerebral cortices (Hill *et al*., 2010). Significant positive correlations were also found for the archetypal sensorimotor-association axis rank (*r* = 0.34 – 0.42, *p_spin_* = 0.033 – 0.01), which incorporates all other ten features to capture divergences between sensorimotor and association cortices (Sydnor *et al.,* 2021) (Suppl. Table 24). Overall, the positive correlations noticed suggest that sulcal width concordances for globularity might tend to concur on multimodal association cortices, more than unimodal sensorimotor regions.

In light of this, we conjectured that further neurobiological links of globularity might be delineated through the associative signature of its genetic relations to sulcal width. To test this intuition, research linking changes in sulcal width and ventricular size to white matter integrity and aging (e.g., Yue *et al*., 1997; Kwon *et al*., 2014) led us to retrieve data for *local brain aging* (Kim *et al*., 2026), *grey matter-white matter intensity contrast* and *cortical thickness* (Kim *et al*., 2025) phenotypes from our GWAS pool. Next, we examined the effect directions of those of their related variants that are also pleiotropically connected to sulcal width, globularity, ventricular volumes and, for contrastive purposes, glymphatic function (Suppl. Tables 21).

A first inspection of concordance ratios indicated substantial heterogeneity in these traits’ interactions (Suppl. Table 21). Since the observed concordance ratios of *aging, grey-white* contrast or *cortical thickness* effect directions with respect to those of *globularity-, glymphatic-* and *ventricular* volume-related SNPs seemed to be limited by the small size of their overlaps, we decided to focus on the spatial distributions of their congruences with sulcal width effects, grounded on more substantial variant intersections, to gain an understanding of how such variability may be organized. Through this approach, we first identified variants driving global sulcal width consistencies (100% concordance) for *grey-white contrast* and *aging*, as well as notable discrepancies (82.25% discordance) for *cortical thickness* (Suppl. Table 22).

Thereafter, we mapped, for each of these phenotypes, the extent of their variants’ effects agreement (*n concordant – n discordant*) with the direction of widening impacts on each of the sulci under investigation. Turning to spatial permutations, we tested for any possible correlations between the derived spatial distributions of effect congruences. Doing this revealed non-trivial, highly positive associations between the spatial arrangements of *grey-white contrast* and *aging* congruency scores (*r* = 0.886, *p_spin_* = 0.0001), which were also positively linked to the four ventricles (*r* = 0.41 – 0.82, *p_spin_* ≤ 0.0002) and globularity (*r* = 0.64 – 0.81, *p_spin_* = 0.0001). Instead, displaying positive correlations to the glymphatic system (*r* = 0.48, *p_spin_* = 0.0004), *cortical thickness* significantly deviated from all of these phenotypes (*r* = -0.497 – -0.74; *p_spin_* ≤ 0.0002) with respect to their patterns of congruence with sulcal widening effects (Fig. 3g; Suppl. Table 23).

Lastly, we asked whether the detected correlations may also be reflected in further associations with the features characterizing Sydnor *et al*.’s (2021) sensorimotor-association axis. We found that the spatial configuration of *aging’*s sulcal width concordances is meaningfully correlated with the *allometric scaling* attribute (*r* = 0.45, *p_spin_*= 0.002), whereas the organization of *grey-white contrast* concordances is both positively connected to *allometric scaling* (*r* = 0.45, *p* = 0.0002) and *evolutionary expansion* (*r* = 0.35, *p_spin_*= 0.04). Conversely, the distribution of convergences between *cortical thickness* and sulcal width shows inverse associations to the last two traits (*allometric scaling, evolutionary expansion*), as well as to the sensorimotor-association rank component (*r* =-0.36, *p_spin_*= 0.01) (Sydnor *et al*., 2021) (Fig. 3h; Suppl. Table 24).

### Specific parietal effects

Contrasting with the observed trend of associatively-oriented, generally congruent effects between sulcal width- and globularity- or ventricular volume-related variants, we identified an exception picking out the parietal lobe: namely, the internal parietal sulcus arose as discordant both for globularity (rs62056163, *C16orf95*) and for the fourth ventricle (rs13224758, *PRKAG2*), also opposing concordant effects for these traits at the sub-parietal sulcus (rs12146713, *NUAK1*) (Fig. 3f). Together with the previously noted discordances converging on the depth of the (right) intraparietal sulcus (consistent with Gregory *et al*., 2017), such parietal-specific impacts seemed prominent: reassuringly, in consonance with the recurrently proposed links between the globular endocranium and the parietal lobe in the literature (Bruner, 2004; Lombard, 2025a; Andirkó *et al*., 2022; Neubauer *et al*., 2018; Gunz *et al*., 2019; Zeloni *et al*., 2026, Bruner & Jacobs, 2013; Bruner *et al*., 2026; Bruner & Gallareto-Sande, 2025), parietal structures also regularly featured in our analysis. This may serve as ground truth proof that, despite being representative of contemporary variation within *Homo sapiens*, the globularity effects examined in this work are of the same nature as the globularity effects from cross-species comparisons at the heart of endocast research.

As for discordant sulcal width effects, their concentration on the internal parietal sulcus suggests an association between globularity and a narrower morphology of this sulcus, which comprises much of the precuneus territory and therefore offers a complementary view to the areal expansion that has been usually linked to this area (Bruner *et al*., 2017; Bruner & Gallareto-Sande, 2025). Particularly, our results may indicate that precuneus expansion could have been accompanied by a reduction in width, overall suggesting a tighter, more compact folding architecture for this sulcus in relation to globularity. Moreover, according to our data, such a morphological configuration could have been accompanied by a relaxation and widening of the sub-parietal sulcus, together with thinning of the intraparietal sulcus. Interestingly, the intraparietal and sub-parietal sulci have been respectively connected to the dorsal and ventral regions of the precuneus, which have been stressed for their structural differences (Bruner *et al*., 2026; Yamaguchi & Jitsuishi, 2024). Overall, the observed patterns of effect may be interpreted as hinting at a redistribution of cortical geometry within the parietal lobe, whereby genetic influences towards an expanded, potentially narrower precuneus could have been paired with genetic effects yielding wider and displaced, or thinner sulci at its ventral and dorsal ends, respectively (Fig. 3e-f).

### Insights from transcriptomics

Armed with these results, we reasoned that, if context-driven trait-specificity can arise from variants’ regulation of pleiotropic gene expression at particular tissues or moments throughout the lifespan (Spence *et al*., 2026), a spatiotemporal inquiry into transcriptomic information may improve our understanding of globularity’s genetic synergies with other traits. Hence, we retrieved normalized microarray datasets from the Allen Brain Atlas (Hawrylycz *et al*., 2012) and processed them via Abagen (Markello *et al*., 2021) in order to obtain regional data for gene expression levels in the adult brain according to Schaefer *et al*.’s (2018) 400 *local-global* parcellation. Furthermore, we extracted RNA-sequencing data from BrainSpan (Li *et al*., 2018) with a view to inspecting developmental trajectories of gene expression linked to our datasets. To further biologically characterize the patterns noticed, we continued capitalizing on the sensorimotor-association archetypal axis (Sydnor *et al*., 2021) for spatial analyses. Additionally, we took advantage from data associated with metabolic pathways (Pourmajidian *et al*., 2026) to understand what energetic costs may be implied in the spatiotemporal interplays observed for globularity, which Molz *et al*. (2024) suggest have been concurrently shaped by co-evolutionary pathways impacted by changes in factors such as energy needs.

For each set of genes related to 31 phenotypes of interest, we began by averaging regional expression levels along Schaefer *et al*.’s (2018) 400-parcellation of the brain, thereby achieving spatial maps for each of our traits’ transcriptomic data. We then carried out pairwise correlations between these brain maps under 10,000 *spin* tests, so as to unravel closer or more distant correspondences among them. Except for some moderately negative correlations (e.g., *ventricular morphology 1 – glymphatic 1: r* = −0.40, *p_spin_* = 0.0087), this procedure did not reveal marked oppositions between our datasets (Suppl. Table 25; Suppl. Fig. 3), possibly in consonance with the formerly observed continuity and interconnectedness of their gene-level intersections.

In order to determine if the derived *gene expression brain maps* contain associations between the phenotypes under research and features of cortical organization, we employed *spin tests* to perform correlations with the components integrating Sydnor *et al*.’s (2021) sensorimotor-association archetypal (SMA) axis (Suppl. Table 26). While correlations were null for some of the traits inspected (e.g., *cortical thickness*, *grey-white contrast, white matter connectome*), 15 of them seemed to align coherently with either sensorimotor or associative features of the axis. Applying multiscale bootstrap resampling with 5000 replicates via *pvclust* to their correlation values with SMA components, the split of these 15 GWAS into a sensorimotor-oriented and an association-oriented cluster was supported (AU = 100%) (Fig. 4b; Suppl. Fig. 4-5). The identified *sensorimotor cluster* tends to comprise traits positively and significantly correlated with SMA components where primary, unimodal cortices rank highest, while negatively and non-trivially correlated with components where primary, unimodal cortices rank lowest. The *association cluster* follows an inverse correlational profile.

**Fig. 4.**
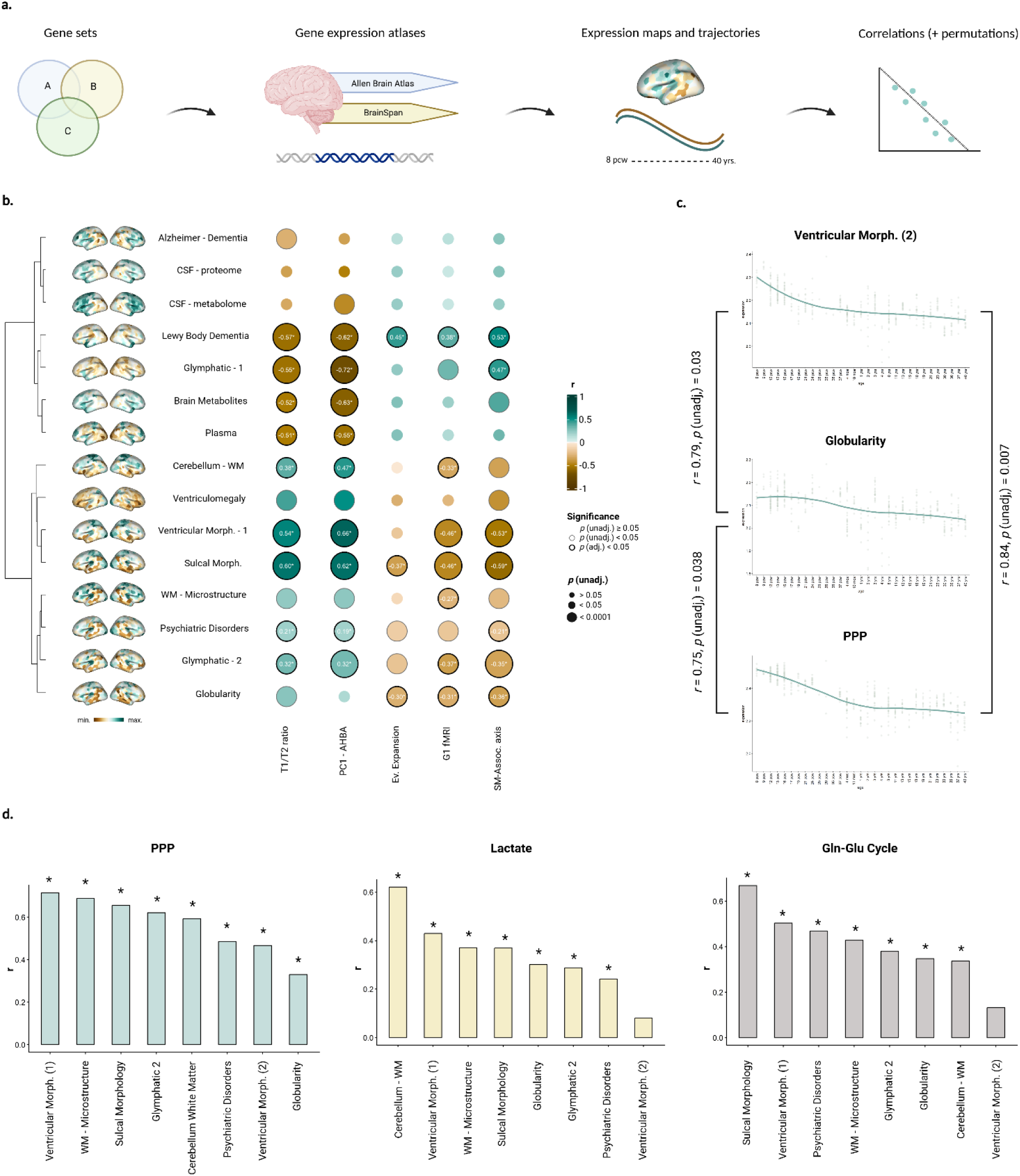
Transcriptomic analyses. **a.** Overview of spatial and temporal gene expression analyses. **b.** 15 datasets bear spatial gene expression patterns that can be dissociated into a *sensorimotor* and an *associative* cluster, according to their correlations with eleven features of cortical organization (5 shown in the heatmap; cf. Suppl. Fig. 4). **c.** Ventricular Morphology (2) and the pentose phosphate pathway constitute the only nominally significant, positive correlates of globularity in terms of their temporal transcriptomic trajectories, characterized by prenatal peaks (cf. Suppl. Tables 27, 29). **d.** The pentose phosphate pathway, lactate and the glutamate-glutamine cycle yield the most consistently significant positive correlations across sensorimotor cluster and ventricular volume-related datasets.

As shown in Fig. 4b, globularity lies within the *sensorimotor* cluster, largely on the basis of its negative spatial correlations with associatively-oriented components of the SMA axis. Among the inversely correlated features for globularity are those indexing cortical areas most evolutionarily expanded from macaque to human (*evolutionary expansion; r* = −0.297, *p_spin_* = 0.04), with greatest relative extents of areal scaling with respect to scaling of total brain size (*allometric scaling*; *r* =-0.301, *p_spin_* = 0.0066), or with largest rates of cerebral blood flow (*CBF*; *r* = −0.288, *p_spin_*= 0.03). Additional correlations are observed for Margulies *et al*.’s (2016) first functional gradient (*G1fMRI*; *r =* −0.308, *p_spin_* = 0.011), where highest scores suggest a progression towards regions functionally involved in abstract, internally-focused cognitive processes, and for the integrative *sensorimotor-association axis* component (*r =* −0.36, *p_spin_* = 0.0038), which unites all other features of the axis within a common gradient. While seemingly in tension with the associative orientation of globularity’s sulcal width concordances, we interpret these results to point to heterochronous interplays of the globular endocranium with different traits throughout distinct stages of brain maturation and aging (cf. Discussion).

Using pre-processed RNA-sequencing data extracted from BrainSpan (Li *et al*., 2018), we proceeded by deriving the temporal expression signature corresponding to each of our datasets’ prioritized genes, averaging their normalized individual trajectories across donors, and then structures (Suppl. Fig. 6, cf. Methods). Datasets with less than four genes represented in BrainSpan were excluded from the analysis. This rendered 30 temporal transcriptomic trajectories consisting of 31 aggregated gene expression values across 31 developmental periods (8 pcw – 40 yrs), which we correlated under the Spearman method to assess their degree of similarity in a pairwise manner. To reduce the likelihood of inflated *p* values, permutations were used to compare the empirical results obtained with 10,000 null models, generated by drawing and correlating temporal expression profiles from random lists of genes of the same size and degree of overlap of the lists under inspection. Focusing on globularity, we saw that its strongest and only nominally significant correlate concerns a second GWAS on structural characteristics and volumes of the brain ventricles (*r* = 0.79, unadjusted *p* = 0.03), with which it shares a decreasing signature from early prenatal (8pcw) peaks of expression (Suppl. Table 27; Fig. 4c).

The transcriptomic profiles of genes prioritized for ventricular morphology-related metrics was steadily connected to those of genes prioritized for the globular braincase across both our temporal (cf. above) and spatial (e.g., *ventricular morphology 2 r* = 0.57, *ventricular morphology 1 r* = 0.38; *p_spin_* = 0.0001) inspections (Suppl. Tables 25, 27; Suppl. Fig. 3). Moreover, by linking their gene expression patterns to primary, unimodal cortical structures (pertaining to the SMA sensorimotor cluster) or prenatal periods, our spatiotemporal exploration implies an early developmental time window for the emergence of these traits’ specific signatures.

In an attempt to sharpen the biological characterization of such a timeframe, we first retrieved curated gene sets associated with 23 metabolic pathways from Pourmajidian *et al*. (2026), and derived spatial expression maps for them using Abagen-preprocessed (Markello *et al*., 2021) data from the Allen Human Brain Atlas (Hawrylycz *et al*., 2012). Computing univariate correlations between these and our transcriptomic maps under 10,000 *spin tests*, we obtained consistently significant associations for the pentose phosphate pathway across all of our sensorimotor-cluster, ventricular volume- (*r* = 0.47-0.72, *p_spin_* = 0.0001) and globularity-related (*r* = 0.38, *p_spin_* = 0.0001) phenotypes (Fig. 4d; Suppl. Table 28). Pathways such as lactate or the glutamate-glutamine cycle also yielded consistent positive correlations with sensorimotor-cluster traits, but failed to achieve such associations with our second ventricular morphology dataset (Fig. 4d; Suppl. Table 28).

We proceeded to ask whether globularity’s relations to metabolic pathways hold for temporal expression profiles. Specifically, we performed correlations between the derived temporal expression trajectories of our initial pool of GWAS and those of our pathways of interest, using BrainSpan data. Following the procedure described above, one dataset and one pathway were excluded from analyses due to insufficient representation of their genes in BrainSpan, yielding a total of 22*30 = 660 pairwise correlations. Out of these, the pentose phosphate pathway emerged as the strongest and only nominally significant positive association for *globularity* (*r* = 0.75, unadjusted *p* = 0.038) and our second ventricular morphology GWAS (*r* = 0.84, unadjusted *p* = 0.007) (Fig. 4c, Suppl. Table 29).

### An evolutionary framing for globularity

We finally looked for variants in our collection of GWAS that are known to be distributed at high frequency in contemporary populations, as compiled by a catalog of single nucleotide changes distinguishing present-day humans from archaic hominins (Kuhlwilm & Boeckx, 2019). In addition to characterizing their effects, we mapped the identified SNPs in terms of their times of evolutionary emergence. Our rationale was that, given the observation of cross-phenotype genetic interplays engaging globularity, evidence of single nucleotide changes impacting brain and cognitive traits may supply indirect, but valuable insights into the evolutionary context that might have shaped the emergence of the globular brain shape.

Suppl. Table 30 summarizes the overlaps reached (142 variants for 25 datasets), with their distributions across lists of modern and archaic SNPs. Among the most extensive intersections with Kuhlwilm & Boeckx’s (2019) catalog, we observe those yielded by network measures of intrinsic brain activity (Bell *et al*., 2022) (Suppl. Table 31). Inquiring into the effect directions of their overlapping variants, we note that *global network efficiency*-related SNPs display a ratio of 100% (4/4) positive impacts, although *resting state fluctuation amplitude* (RSFA), an important vascular component underlying *global network efficiency* (Bell *et al*., 2022), implies greater heterogeneity in the effects related to this trait (6/10 = 60% positive impacts). In between, variants intersecting with DMN and subcortical-cerebellar networks present 62.5% (5/8) positive effects (Fig. 5a).

**Fig. 5.**
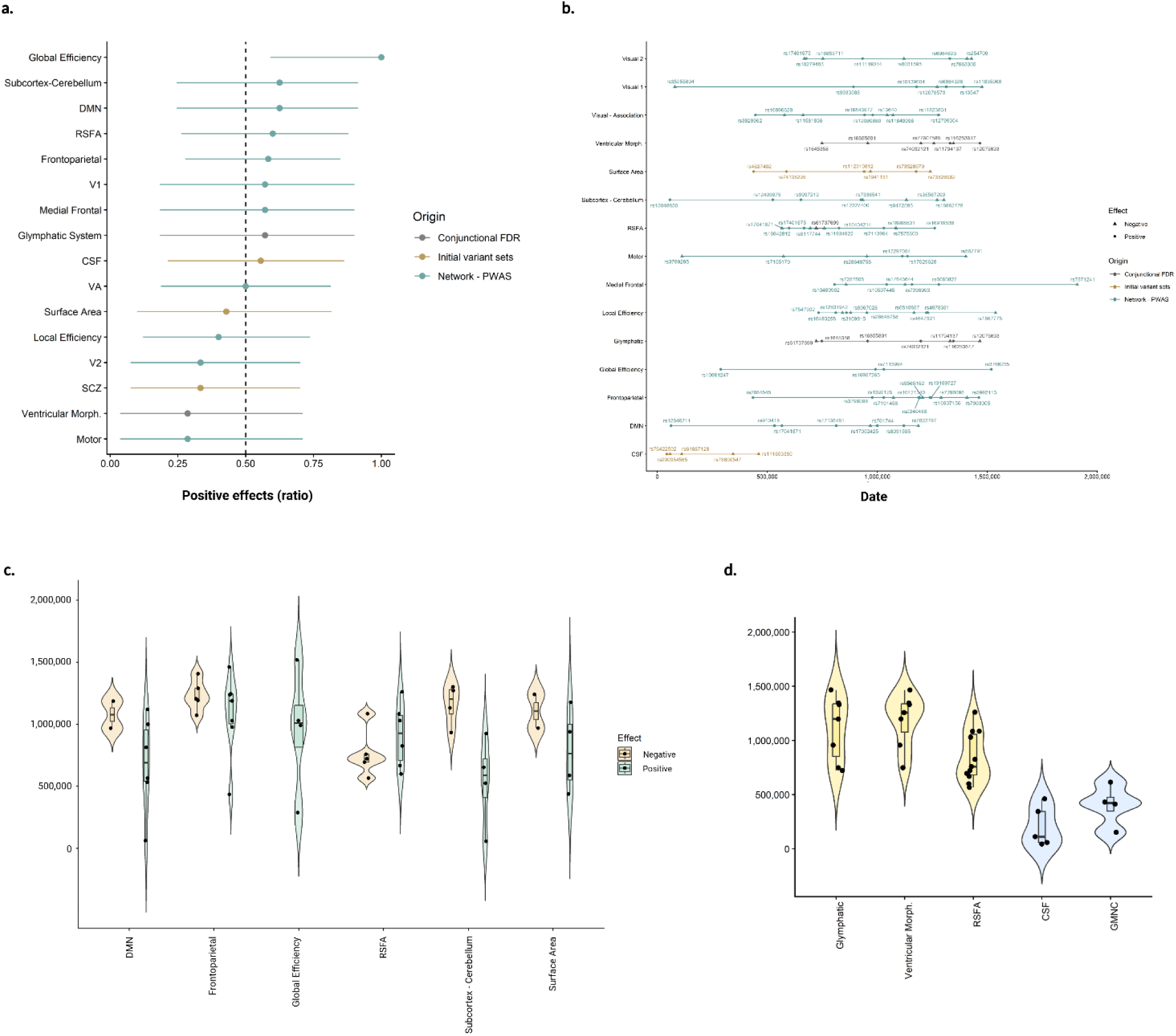
An evolutionary framework for the emergence of globularity. **a.** Proportions of positive effect directions for phenotypes with the largest overlaps (≥ 7) with evolutionary annotations. **b.** Emergence times of *Homo sapiens*-distinctive high-frequency variants associated with brain-related traits. **c.** Temporal distribution of variants with positive and negative effects for network metrics of brain activity and measures of surface area. **d.** Parallel, modern times of emergence for CSF and *GMNC* high-frequency variants. Timeframes for *Glymphatic, RSFA* and *Ventricular Morphology* single nucleotide changes are shown for comparison.

Noteworthy overlaps of nine variants also obtain for the cerebrospinal fluid proteome (Western *et al*., 2024) and Sha *et al*.’s (2025) SNPs for pleiotropies between psychiatric disorders and brain morphological characteristics, where we see that they mostly converge on schizophrenia (8/9 = 88.89%) and surface area (7/9 = 77.78%) (Suppl. Table 32). Moreover, boosting our chances of variant discovery beyond the genome-wide significance threshold, conjunctional FDR allows us to detect eleven single nucleotide changes enriched for the glymphatic (Huang *et al*., 2025) (4/7 = 57.14% positive impacts) and/or ventricular systems (2/7 = 28.57% positive impacts) (Ge *et al*., 2024) (Suppl. Table 33).

To examine the temporal makeup of our data, we then use a collection of times of emergence (Andirkó *et al*., 2022) for variants in Kuhlwilm & Boeckx’s (2019) catalog as predicted by the Genealogical Estimation of Variant Age method (Albers & McVean, 2020) (Suppl. Fig. 7; Suppl. Table 34). Through this method, we observe that variants pleiotropically linked to the glymphatic and ventricular systems are traced back to periods substantially prior to the emergence of the globular brain shape, approximately from 1.5 Mya to 700 Kya. Similar results obtain for most network metrics of brain function, with the exception of some related variants that expand to more recent evolution (Fig. 5b). Those linked to frontoparietal, default-mode and subcortico-cerebellar measures exhibit progressively more positive impacts as they do so (Fig. 5c), suggesting that their overall positive orientation (Fig. 5a) may be partly related to recent changes in their evolutionary history.

Remarkably, whereas networks like the subcortico-cerebellar or the default-mode reach modern times of emergence for their latest SNPs, we notice that variants linked to the cerebrospinal fluid (CSF) proteome are consistently mapped to a period comprising the last 500 Ky of human evolution (Fig. 5b,d). Hence, the CSF proteome features prominently for concentrating all of its mapped high-frequency single nucleotide changes in a recent evolutionary period of known significance for the emergence of the globular endocranium.

Noteworthy in this context are high-frequency, *sapiens*-derived variants linked to *GMNC*. Through an examination of the effects of non-coding Neanderthal-specific variants in a large human cohort, this gene has been recently highlighted for its broad impacts on brain morphological features from cortical and sulcal measurements to white matter diffusion as well as ventricular, choroid plexus and subcortical volumes (Zeloni *et al*., 2026). Thus, under the hypothesis that they may be connected to brain shape differences between *Homo sapiens* and archaic hominins, we detect that “modern” *GMNC*-associated variants follow a recent evolutionary profile, seemingly concurrent with that of SNPs linked to the CSF proteome (Fig. 5d).

### Modern links to the cerebrospinal fluid proteome

The finding of *Homo sapiens*-distinctive high-frequency single nucleotide changes in the CSF proteome arising more recently in our lineage, in parallel with the relevant timeframe for the evolution of globularity, bears on recently proposed connections between the globular endocranium and CSF-driven hydrostatic pressure in the brain ventricles (Molz *et al*., 2024), as well as between the cerebrospinal fluid secretome and embryonic brain development (Courtney *et al*., 2025). It also bears on the hypothesis that cerebral blood flow rate increased disproportionately faster than brain size during human evolution (Seymour *et al*., 2016; Harrington *et al*., 2019). This motivates us to conduct a more detailed inspection of signals connected to the CSF proteome.

As for recently emerging, high-frequency variants overlapping with Kuhlwilm & Boeckx’s (2019) catalog, we first observe that their major alleles are associated with upregulated levels of proteins that are predominantly intracellular or membrane-bound: GSTP1, CTSD, T-cell antigen CD7, BTN2A2, and B4GALT2. By contrast, downregulated proteins (PSPN, MSMB, HSPG2, GCPII) tend to be secreted or extracellular matrix proteins, in some cases (PSPN, HSPG2) related to growth factors. Both upregulated and downregulated proteins have been implicated in relevant processes for the development of traits that our results relate to globularity (cf. Discussion), such as neurite initiation throughout the maturation of the cerebral cortex (GSTP1; Liu *et al*., 2021), perivascular fluid transport and brain clearance (HSPG2; Singh *et al*., 2025) or, through glycan synthesis (B4GALT2), the inflation of the brain ventricles (Lowery & Sive, 2009), neurite outgrowth, axon guidance and synaptogenesis (Bonfanti, 2006; Gouveia *et al*., 2012; Parkinson *et al*., 2013; Noel *et al*., 2025).

Notably, beyond modern high-frequency single nucleotide changes, we notice further links between the cerebrospinal fluid proteome and recent evolutionary events potentially linked to globularity. In particular, our initial hypergeometric tests reveal a significant variant-level overlap (*p* = 7.6E-07) between CSF proteome-related SNPs (Western *et al*., 2024) and independent variants prioritized for the volume of the cerebellum (Moberget *et al*., 2026), a structure with strong associations to globularity (Gunz *et al*., 2019; McCoy *et al*., 2017; Buisan *et al*., 2022; Kochiyama *et al*., 2018; Takagi & Kondo, 2025). The shared variants are mostly concordant in their effects with all of the 23 cerebellar components studied (Suppl. Table 35; 5/7 variants with > 86.96% concordances), suggesting an association between increased levels of their corresponding proteins in the CSF and general increases in cerebellar size.

### *GNA12, NUAK1, C16orf95, GMNC* and the 8p23.1 cytogenetic region

Throughout hypergeometric, conjunctional FDR or evolutionary analyses, our results regularly pick out variants mapped to *GNA12, NUAK1, C16orf95* and *GMNC*. These genes are among the most frequent across datasets in our initial pool of GWAS (Suppl. Table 8), which suggests their recurrent links to globularity-related phenotypes. In light of this, recent associations of *GMNC, C16orf95* and *GNA12* with alleles increasing ventricular volumes and causing CSF-specific molecular signatures, as well as with high expression in cilia-abundant tissues (e.g., testis, fallopian tubes), deserve mention, as they insinuate mechanistic links between coordinated ciliary beating in ependymal surfaces, CSF flow and ventricular sizes (García-González *et al*., 2026). Especially through a highly pleiotropic variant frequently featuring in our data (rs12146713), *NUAK1* has also been linked to CSF flow and ventricular volume increases (García-González *et al*., 2026), as well as with neurocranial shape (Goovaerts *et al*., 2025). Additionally, all four genes have been usually implicated in Alzheimer’s disease (Lasagna-Reeves *et al*., 2016; Timsina *et al*., 2026), consistent with our observation of concordances between globularity and brain aging. In the case of *C16orf95*, connections to neurodegeneration and relevant pathways (*lipid signaling, cilia, microtubule*) may also be drawn from or for its neighboring gene *MAP1LC3B,* as several studies (Handrigan *et al*., 2013; Zhang *et al*., 2025; Timsina *et al*., 2026) have suggested common regulatory variants to equally impact both *C16orf95* and *MAP1LC3B*. A known regulator of autophagic flux (Chen *et al*., 2025; Yao *et al*., 2025), the latter gene makes relations between autophagy and the regulation of neural stem cell proliferation and differentiation (Leonardi *et al*., 2025) worthy of notice in the context of our findings.

From an evolutionary perspective, it is also interesting that these genes, or regions regulating their expression, feature prominently in various lists derived from paleogenomic analyses, suggesting that evolutionary variants may have tuned in the globularization phase in a similar way to those common variants highlighted in our inspections. *C16orf95* is associated with a differentially methylated region in *Homo sapiens* compared to other hominins (Gokhman *et al*., 2020), pointing to reduced expression of the gene in *sapiens.* This differentially methylated region contains the single virtually fixed variant associated with *C16orf95* in Kuhlwilm & Boeckx (2019). Consistently with the expression-decreasing effects of this region, we find via GTEx that several *C16orf95* variants positively affecting globularity in Molz *et al*. (2024) are linked to a lower expression of this gene (e.g., rs9933149, rs1108048, rs35427507, rs4843578). Regarding *GMNC*, the variants mapped above (Fig. 5d) fall within a regulatory island (Moriano *et al*., 2025), i.e., a regulatory region depleted of mutations in other hominins. Moreover, this island corresponds to a region with diverged 3D architecture in McArthur *et al*.’s (2025) comparison of hominin genomes. Similarly to *GMNC,* we note that *NUAK1* is the predicted target of several regulatory islands, which furthermore rank among those in Moriano *et al*. (2025) exhibiting the largest number of unique transcription factor binding motif modification. As for the *AMZ1*/*GNA12* region, it exhibits an excess of high frequency mutations in Neanderthal/Denisovan genomes compared to *sapiens* in Kuhlwilm & Boeckx (2019), indicating another point of accentuated divergence.

Finally, beyond recurrent associations to these genes, our results also frequently associate globularity and brain-related phenotypes to the cytogenetic region 8p23.1. This region, reported in Molz *et al*. (2024), harbors relevant loci concordantly linking either the fourth brain ventricle or globularity to autism spectrum disorders in our analyses (Suppl. Table 36). Moreover, similar to the paleogenomic signals found for *GNA12, NUAK1, C16orf95* and *GMNC*, this region harbors variants that, apart from distinguishing modern and archaic hominins, have been shown to bear active regulatory activity in neural progenitor cells (Weiss *et al*., 2021).

## Discussion

Our work aimed to investigate the neurobiological underpinnings of the human globular endocranium, a *sapiens*-specific trait thought to result from an early-life apomorphic growth trajectory of the brain (Hublin *et al*., 2015; Gunz *et al*., 2010). In the absence of fossil brain tissue or experimental models that can reliably mirror the complete maturational timeline leading to a globular brain shape (currently outside the reach of organoid models), GWAS were used to explore the genetic foundations of globularity in relation to relevant brain and cognitive phenotypes, on evolutionary and developmental timescales. Conjunctional FDR, neuroimaging techniques, evolutionary annotations and transcriptomic data were combined to try to overcome inherent limitations of GWAS-based research: namely, the influences of trait-irrelevant factors on gene prioritization and discovery (Spence *et al*., 2026), the conservatism of genome-wide significance thresholds for low-frequency variants (Fadista *et al*., 2016; Chen *et al*., 2021; Mendes *et al*., 2026), or the difficulty to capture bidirectional dimensions of genetic overlap by global genetic correlations (Sha *et al*., 2025), which nevertheless helped us confirm homogeneous relations between traits.

While not exempt of limitations, our study broadens our view of the globular phenotype. We show that value can be gained from the *variant*-level’s advantageousness for characterizing specificities within and between phenotypes, coupled with the widespread pleiotropies expectable for complex, polygenic, brain-related traits (Watanabe *et al*., 2019) under stabilizing selection (Zhu *et al*., 2026). Particularly, leveraging *variant*-level overlaps of multiple globularity-related traits through GWAS, we are able to attain insights at higher levels of resolution. For instance, both earlier (Gunz *et al*., 2019) and later (Molz *et al*., 2024) inquiries into globularity have stressed effects on genes involved, among others, in myelination pathways and white matter microstructure: here, we separate concordant (FA, MO, ICVF) from discordant white matter *microstructural* parameters (L1, L2, L3, MD, OD, ISOVF), seemingly pointing to a link of globularity with the formation of denser, more organized connectivity networks, with the particular engagement of regions such as the left Heschl gyrus, the pallidum, the right amygdala or the parietal inferior gyri at the *macrostructural* level.

In agreement with Sha *et al*.’s (2025) endeavor to capture the full dimensionality of genetic overlap between phenotypes, our use of conjunctional FDR further allows us to understand underlying neurobiological connections between globularity and related traits through a comprehensive lens. As an example, we accentuate our method’s capacity to identify highly concordant and discordant cerebellar components with respect to globularity, refining previously observed associations between increased endocranial globularity and somewhat conflicting signals of increased or decreased grey matter throughout different cerebellar regions (Gunz *et al*., 2019). In addition, beyond distinguishing between globally and more specifically distributed homogeneous effects between globularity and brain-related traits, our approach actually detects specific variants contributing to such pleiotropic relations, together with relevant genes (e.g., *C16orf95*) or cytogenetic regions (8p23.1) to which these variants are consistently mapped. Ultimately, our focus on concordant and discordant dimensions of variant-level overlaps (either genome-wide or conjunctional FDR-significant) also allows us to recognize temporal patterns in globularity-related evolutionary timelines, whereby, for instance, impacts on subcortico-cerebellar and default-mode network connectivity are increasingly positive as more recent evolution is reached.

Our study likewise contributes an expanded view of how a globular brain may emerge and age —how the globular phenotype might manifest throughout the lifespan. Specifically, we are able to reconstruct a developmental timeline for globularity-related phenotypic impacts by reconciling tensions arising from our data: namely, an ‘associative cortex’ orientation for sulcal width and aging effects, alongside a ‘sensorimotor cortex’ orientation for the spatial expression of globularity-prioritized genes.

These genes also display prenatal peaks of expression, consistent with the notion that gestational stages of brain development might already be critical for globularity well before the visible emergence of a globular neurocranium. Regarding their predominant spatial expression in sensorimotor, primary cortices, it points to early-developed unimodal cortical regions as likely targets of globularity-associated alterations in neural architectures (Gunz *et al*., 2019). Congruently with this, our inspection of white matter connectivity effects stressed concordances with cortical regions such as the left Heschl gyrus (primary auditory cortex) and the postcentral and precentral gyri (primary somatosensory and primary motor cortices), together with subcortical structures (e.g., thalamus, putamen, hippocampus, amygdala, pallidum). The early ontogeny of these subcortical and primary cortical structures’ connectivity during gestational stages might be an important factor in the shaping of the globular brain.

In this context, the transcriptomic- and effect-level alignments between *ventricular volumes* and *globularity* indicates a potentially relevant role for the embryonic formation of an evolutionarily expanded ventricular system from the lumen of the neural tube, supporting the relevance of the neuroepithelium for brain evolution (Benito-Kwiecinski *et al*., 2021). One might speculate that a globularity-associated increase in ventricular size in initial developmental stages could be linked to an equally expanded germinal neuroepithelium lining the ventricles, thus allowing a rise in the pool of stem cells that would go on to generate the vast majority of CNS cells (Fame & Lehtinen, 2020).

Importantly, our results also stress a possible role of the embryonic cerebrospinal fluid (eCSF). Secreted by the neuroepithelium to inflate the ventricles, this fluid has been claimed to regulate neuronal proliferation and determination, whereas contact with it and the factors it contains have been suggested as prerequisites for the production of early, multipotent neuroepithelial cells (Lowery & Sive, 2009; Shen, 2018). Differently from previous implications of the cerebrospinal fluid in brain development (Desmond *et al*., 2005) and, particularly, the shaping of the globular endocranium through its hydrostatic pressure on the ventricles (Molz *et al*., 2024), our research provides evidence that changes in its proteome could have occurred in recent evolution. Since proteomic alterations in the CSF, which is richer in pre-/peri-natal stages, have been linked to the stimulation of neural progenitors lining the ventricles and shifts in the developmental trajectory of the brain (Zappaterra *et al*., 2007; Parada *et al*., 2006; Courtney *et al*., 2025; Tanabe *et al*., 2025), it seems worth accentuating the potential developmental import of the cerebrospinal fluid proteomic changes observed in our work. Indeed, some of the *sapiens*-distinctive, high frequency modifications we find imply the upregulation of proteins that have been involved in relevant processes like neurite initiation (e.g., GSTP1; Liu *et al*., 2021) and outgrowth (B4GALT2; Noel *et al*., 2025). It may thus be hypothesized that an expansion of the ventricles related to increased cerebrospinal fluid pressure throughout development (Molz *et al*., 2024), paired with changes in the proteomic composition of the CSF, could be a relevant mechanism for the conformation of the aforementioned globularity-associated characteristics of white matter connectivity. Such an evolutionary relevance of changes in CSF substrates would cohere with current research on the *ADSL* gene, for which a high-frequency haplotype has been identified to modify CSF concentrations of succinyladenosine (Ju *et al*., 2025).

The transcriptomic alignments between globularity, the brain ventricles and the pentose phosphate metabolic pathway also encourage further investigation of the latter’s roles in brain development. Relevantly, *PPP* has been traditionally involved in cellular biosynthetic processes of potential significance for the maintenance of redox homeostasis (Pourmajidian *et al*., 2026; Camandola *et al*., 2017), which could be important for contexts of rapid cell proliferation, likely to generate cellular stress (Stincone *et al*., 2014), as presumable for a prenatally increased pool of neuroepithelial cells. More recently, *PPP* has been linked to axonal extension and regeneration (Zhang *et al*., 2025; Song *et al*., 2026), as well as the modulation (preventing premature transitions) of radial glial cell specification (Mil *et al*., 2025). Interestingly, TKTL1, a gene belonging to the PPP, is known to harbor a protein-coding mutation distinguishing most contemporary humans from archaic hominins. Although *in-vitro* experimental work points to a possible effect of this mutation on brain development (Pinson *et al*., 2022), phenotypic data from contemporary humans carrying the ancestral allele question the overall impact of this mutation (Molz *et al*., 2025).

On the other hand, the distribution of interactions between globularity and parameters like sulcal width leads us to infer that globularity may also impact the later development of higher order, transmodal association cortices throughout postnatal life. Indeed, the neonatal period has been said to constitute a critical phase of brain shape formation, most strongly represented by measures of fractal dimensionality, gyrification and surface area (Krohn *et al*., 2026). During this period, sulci that emerge later prenatally show faster deepening than earlier emerging sulci, becoming wider, thicker, denser with tissue, and less concave (Tung *et al*., 2026). Crucially, later emerging sulci have been localized towards the associative cortex in a bimodal, linear-to-complex taxonomy of adult brain morphology related to timing of fetal sulcation (Snyder *et al*., 2024). In line with this, our positive correlation between globularity’s sulcal width concordance scores and the associative features of the sensorimotor-association archetypal axis (Sydnor *et al*., 2021) might point to early postnatal development, coinciding with the globularization phase, as a relevant developmental time window for the widening of transmodally-oriented, later developed sulci. In this sense, our results diverge from nominal, not fully consistent relations between globularity and sulcal measurements already hinted at in the literature (Zeloni *et al*., 2026) and advance the current state of the art by pointing to a consistent, spatially organized convergence of globularity-related variants on sulcal width effects (opposed to more selective relations with the depth of sulci such as the intraparietal sulcus).

Widening impacts of the sort noticed have been associated with brain aging (Yue *et al*., 1997) in adult life, yet, developmentally, they have been linked to neural reorganizations aimed at optimizing transmission efficiency (Shan *et al*., 2026). In this sense, the associative pattern of our observed width effects might provide an indirect, complementary view of globularity-related influences on brain connectivity. Particularly, whereas the early connectivity signature arising from our data predominantly involves long-range connections to and from subcortical as well as primary cortical structures, postnatal sulcal widening may be linked to later modifications in brain connectivity. For instance, sulcal widening has been hypothesized to accommodate the maturation of short-range U-fibers (Sowell *et al*., 2004), marking a shift from generalized to specialized neural connectivity (Petanjek *et al*., 2011; Shan *et al*., 2026). Such an account could fit the mismatch between our connectivity and our sulcal width results, by relating widening effects to later developments in neurotransmission rather than to the formation of dense neural connections influencing earlier-developed structures.

Our work, moreover, illuminates the relation between globularity, the glymphatic system and the volume of the brain ventricles. Previous research had hypothesized links between globularity, increased ventricular volumes, and hydrostatic pressure from the cerebrospinal fluid during development (Molz *et al*., 2024); nonetheless, despite its known negative associations with ventricular enlargements (Wu *et al*., 2025; Li *et al*., 2023) and its role supporting a constant influx of CSF driving the export of brain metabolic waste (Hablitz & Nedergaard, 2021), the glymphatic system had remained unexplored in relation to the globular brain shape. With our approach, we contribute strong evidence for variant effect concordances between globularity- and ventricle-related traits, as well as discordances between these and the DTI-ALPS metric of glymphatic function.

While such discordances are consistent with literature indicating ventricular expansions (Barron *et al*., 1976; Nestor *et al*., 2008; Ferrarini *et al*., 2006; Apostolova *et al*., 2012) and glymphatic dysfunction (Hsu *et al*., 2023; Huang *et al*., 2024) in aging and neurodegenerative diseases, recent studies have also suggested that, in developmental conditions where the ventricles are dilated, glymphatic dysfunction may also be implicated. This could relevantly be the case of pediatric hydrocephalus (Zhao *et al*., 2025; Emmert *et al*., 2019). Hence, since we have argued for ontogenetically early interactions between globularity and size-increases of the brain ventricles, we may hypothesize that the evolutionary emergence of the globular braincase might have lent decreased baseline rates of glymphatic efficiency already at initial developmental stages, such as neonatal periods when glymphatic flow has been described to mature dynamically (Lin *et al*., 2024; Peng, 2025). Coherently with the proposed postnatal coupling between the *globularization* phase and effects on sulcal width —another phenotype departing from glymphatic clearance in our results—, this brings us to ask what the developmental impacts of a globularity-related reduction in glymphatic efficiency could be.

Two possibilities offer themselves to us. First, in light of the glymphatic system’s roles in the elimination of soluble proteins and metabolites from the central nervous system, as well as in the distribution of compounds such as glucose, lipids, amino acids, growth factors and neuromodulators (Jessen *et al*., 2015), a reduction in glymphatic fluid mobility could be speculated to allow a longer retention of necessary nutrients and products for the metabolically demanding periods of postnatal brain development (Kuzawa *et al*., 2014). In that sense, evidence pointing to connections between higher accumulations of lipids in human neurons and their prolonged synaptic maturation (Rava *et al*., 2026) is relevant: from such a standpoint, reduced glymphatic fluid dynamics could be suggested to contribute to lipid retention needed to fulfil the substantial metabolic demands of protracted neuronal maturation. Astrocytes may be relevant in this regard, as they have been implicated in human brain neoteny (Ciuba *et al*., 2026), the timing of neuron maturation (Steiner *et al*., 2026) and, through Aquaporin-4 (AQP4) water channels at their perivascular endfeet (Gomolka *et al*., 2023; Mestre *et al*., 2018), glymphatic function.

Second, given the recently uncovered negative correlations between cerebral blood volume and glymphatic influx (Li *et al*., 2026), it may be hypothesized that a decrease in ontogenetic baseline levels of glymphatic flow could have been paired with a corresponding increase in baseline levels of cerebral blood volume, and thus cerebral blood flow, again throughout the metabolically expensive periods of brain development after birth (Kuzawa *et al*., 2014). According to Li *et al*. (2026), the mechanistic explanation for such a relation would lie on a facilitatory effect of cerebral vasoconstriction towards increased CSF influx into the brain parenchyma, as opposed to an effect of cerebral vasodilation towards decreased CSF influx into the brain parenchyma. This account fits the positive correlation between globularity’s sulcal width concordances and the cerebral blood flow feature of Sydnor et al.’s (2021) sensorimotor-association axis, negatively related to glymphatic-associated sulcal width concordances. If so, it would suggest that changes in the shape of the brain may be coupled with modifications in blood supply, a trait claimed to have undergone faster increases than brain volume during human evolution (Seymour *et al*., 2016).

Additionally, through their convergence with sulcal width effects and their interactions with ventricle- and glymphatic-related phenotypes, our results seem to establish an association between *globularity* and *brain aging*. Namely, we note that *globularity* and *aging* bear a common pattern of concordant associations with ventricular volumes and sulcal width, as well as discordant associations with glymphatic function. This is consistent with research that has identified ventricular enlargements (Coffey *et al*., 1992), sulcal widening (Kochunov *et al*., 2005) and glymphatic system disruptions (Xiong *et al*., 2024) in aging-related conditions. Additionally, it may be in line with our finding of a high-frequency variant (rs114959832) positively impacting brain aging (i.e., related to an older brain phenotype) in modern human populations and mapped to ZEB2, a key regulator of a cell shape transition mechanism driving evolutionary human brain expansion (Benito-Kwiecinski *et al*., 2021).

Notably, the established link between globularity and aging seems to conform to an associative cortex orientation. This is suggested by the positive spatial correlations obtained between their distribution of sulcal width concordances throughout the brain and associative features of the archetypal sensorimotor-association axis (Sydnor *et al*., 2021). Specifically, *aging* is positively associated with an *allometric scaling* component, which could indicate an increased vulnerability to *age*-related structural and functional changes in cortical regions displaying greater degrees of areal expansion relative to overall brain size. This seems in agreement not only with evidence of regionally selective susceptibilities to brain aging (Pandya & Patani, 2021; Kim *et al*., 2026), but also with research pointing to a human-specific association between strong aging effects and large relative cortical expansion, especially in higher-order cognitive regions (Vickery *et al*., 2024; Filippi *et al*., 2023).

More broadly, the relation between allometrically scaled cortical areas and other associative features of the archetypal sensorimotor-association axis (Sydnor *et al*., 2021), such as *cerebral blood flow, aerobic glycolysis* or *evolutionary expansion*, suggests that aging vulnerability may lie on highly evolutionarily expanded and metabolically demanding brain regions. These are known to be regions with most protracted developmental timelines in human ontogeny (Casey *et al*., 2005; Baum *et al*., 2020; Sydnor *et al*., 2021), which may be connected to a long-standing idea associating greater age-related susceptibility to oxidative stress and neurodegeneration with increased aerobic metabolism and plasticity of (“neotenous”) brain regions retaining juvenile characteristics for longer (Bufill *et al*., 2011).

Beyond extending our view of globularity from gestational developmental stages to brain aging, our research also bears on the globular endocranium and its associated brain phenotypes from an evolutionary perspective. Concretely, by identifying and temporally mapping high-frequency GWAS hits related to CNS traits, we reveal changes accumulating in variable directions throughout the last 2 My of human evolution, consistent with the subtle, multifactorial differences suggested to have driven the emergence of the ‘modern’ human condition (Caporale *et al*., 2025; Bennett & Fu, 2024; Scerri *et al*., 2018; Scerri & Will, 2023; Kuhlwilm & Boeckx, 2019). Furthermore, by offering insights into the phenotypic consequences of changes undergone at different times, we complement previous efforts to reconstruct an evolutionary history of our species using genetic data. Particularly, in relative agreement with Andirkó *et al*.’s (2022) bimodal distribution of *Homo sapiens*-derived alleles, we distinguish an earlier concentration of variants, approximately, from 1.5 Mya to 500 Kya, and a later concentration of variants, approximately, throughout the last 500 Ky. We hypothesize that the earliest changes may be partly associated with the accelerated brain growth (Jerison, 2012; Gómez-Robles *et al*., 2024) and significant neocortical expansions (Barton, 2007) experienced in the human lineage over the past 2 million years (Libedinsky *et al*., 2025), as suggested by increasingly positive effects on networks linked to the evolutionary expansion of the cortex (*frontoparietal, DMN:* Mantini *et al*., 2013; Wei *et al*., 2019) and predominantly positive effects on the glymphatic system —whose efficient functioning has been related to cortical folding (Kameya *et al*., 2024). Instead, more recent changes seem to stress the significance of the cerebrospinal fluid proteome throughout evolutionary periods that have been most closely linked to the emergence of the globular brain shape (e.g., last 100 Ky: Neubauer *et al*., 2018; 300 – 120 Ky: Molz *et al*., 2024).

Finally, our study’s contributions also concern the cognitive implications of the neurobiological interplays uncovered for globularity and multiple brain-related traits. Beginning with early developmental effects, our results might be able to illuminate heterogeneous associations observed in the literature for ventricular volume measures. Concordantly linked to the globular endocranium, these have been genetically positively correlated with verbal numerical reasoning and general cognitive function with nominal significance (Ge *et al*., 2024), but phenotypically associated with decreased frontotemporal connectivity in relation to language, crucially in adults (Amelink *et al*., 2025). Instead, our regional connectivity patterns for globularity or the brain ventricles highlight concordances with subcortical structures, such as the hippocampus, the pallidum, the amygdala, the putamen and the thalamus. Some of these are involved in language-responsive (e.g., thalamus) or language-selective (e.g., amygdala, hippocampus) extended networks beyond the ‘core’ left frontal and temporal areas (Wolna *et al*., 2026), which may suggest, alongside their *edge*- and *node*-level concordances, a link between globularity and an ‘extension’ of cognitive networks to the subcortex. This may be attributed to different functions traditionally associated with subcortical structures, like learning and memory (hippocampus: Jarrard, 1993; Eichenbaum *et al*., 1996) or the integration of CNS signals (thalamus: Shine *et al*., 2023), which hypothetically may have lent the subcortex enhanced roles in global brain function.

Further, the emphasis in structures such as the amygdala suggests impacts on social cognition (Adolphs, 2010; Phelps, 2006). Relevantly, the basolateral nucleus of the amygdala has been found to present the most prolonged plasticity among social brain areas, all following extended developmental trajectories (Cerrito *et al*., 2024). Moreover, the lateral nucleus of the amygdala is larger in humans compared to other apes (Barger *et al*., 2007), a change claimed to have convergently reduced reactive aggressiveness in both domesticated and cooperatively breeding species, potentially facilitating the emergence of proactive prosociality in humans (Cerrito & Burkart, 2023). Thus, pointing to notable socio-cognitive effects, high degrees of concordance between globularity, ventricular volumes and the amygdala’s *edge-* and *node*-level connectivity might be consistent with those found for cortical areas like the insula, which has been involved in the interface between sensation, emotion and cognition through its contribution to interoceptive and exteroceptive sensory processing (Zhang *et al*., 2024), emotionality (Gasquoine, 2014) and motivation (Namkung *et al*., 2017).

Other highly concordant cortical areas stressed in our connectivity inspections seem to reflect, again, language-related influences beyond frontotemporal regions traditionally linked to linguistic cognition. In this regard, asymmetries concerning the Heschl gyrus (*edge-level* left = 95.56% vs. right = 23.33%) and superior temporal pole (*node-level* left = 83.33% vs. right = 23.33%) are especially noteworthy, as the left counterparts of these areas have been deemed key for auditory and semantic processing (Roll, 2024).

From our connectivity analyses a picture emerges where early globularity-related influences seem to support the formation of networks serving linguistic (particularly, auditory and semantic), social and global cognitive function. Given the early ontogenetic (specifically, prenatal) signature insinuated for these effects, they may be related to evidence pointing to fetal stages of development as a period of rapid white matter maturation foundational for future cognitive function, such that *edge-centric* network controllability properties (indicative of the ease with which the brain moves between distinct mental states based on white matter connectivity) arising throughout the third trimester of gestation and exhibited at birth are predictive of social engagement and language outcomes in toddlerhood (Sun *et al*., 2023, 2025, 2026).

Such social and linguistic impacts may be echoed by our results for cerebellar size. Specifically, the concordances observed between cerebellar volumes and globularity accentuate a component (*component 03*) in apparent overlap with a cerebellar subregion recently shown to be highly selective for language (Casto *et al*., 2026). Indeed, regions through which the third component spans, such as Crus II and lobules VIIb/VIIIa, have also been implicated in the ‘extended language network’ (Wolna *et al*., 2026), similarly to subcortical structures highlighted by our inquiry into white matter connectivity.

As for globularity-related cognitive implications with later developmental signatures, we may emphasize those of the geometric redistribution of the parietal cortices delineated through our data’s specific parietal effects. These first point to a thinning of the intraparietal sulcus, especially in the right hemisphere. Interestingly, interruptions in this sulcus, related to decreases in sulcal depth (Roll, 2024), have been associated with improved performance on memory and language tasks (Santacroce *et al*., 2024). Moreover, our parietal morphology influences are recurrently related to the precuneus, whose expansion has been often linked to visuo-spatial integration, memory, autonoetic consciousness, body awareness and imagination (Bruner & Gallareto-Sande, 2025; Bruner *et al*., 2014). In this way, whereas our initial developmental impacts seem to relate to connectivity-based scaffolds on semantic, auditory and emotional processing of potential import for early human ontogeny, our later postnatal effects additionally seem to tackle higher-order cognition linked to inventiveness and the construction of the self.

### Limitations

Our approach contributes insights into the genetic architecture of the globular endocranium, its overlaps with the underlying neurobiology of other brain-related traits, and its evolutionary and developmental implications for the human brain and cognition. Nonetheless, there exist constraints on the generalizability of our results and the extent to which they support some of our proposed mechanistic explanations.

First, a substantial part of our project has been devoted to comparing *variant*-level pleiotropies in both their concordant and discordant dimensions. While this strategy has allowed overcoming limitations of methods traditionally used to assess commonalities in phenotypes’ genetic architectures (e.g., pleiotropy obscured by genetic correlations due to bidirectional impacts on traits; Sha *et al*., 2025), it is still restricted in several ways. For instance, attaining a quantitative measure that allows accounting for simultaneous opposed effects between two phenotypes remains problematic: while our respective attribution of positive and negative scores to concordant and discordant effect directions has permitted disentangling spatial distributions of contrasting genetic influences, these oppositions are not properly explained when they converge on a common brain region. Moreover, whereas analyses at the *variant*-level seem to allow higher degrees of granularity, they have also implied working with reduced observations, which has importantly hampered statistical inference. More broadly, despite advantages previously outlined, the *variant*-level has been recently shown to bear several constraints. In that sense, like other methods requiring intersections of variants across multiple studies (e.g., multi-trait colocalization, GWAS meta-analyses), our approach may be limited by current challenges in GWAS integration and fine-mapping for variant interpretation: from heterogeneity in SNP coverage due to variations in genotyping arrays and imputation panels, to LD dropout-driven hindrances in statistical fine-mapping, among others (Ahmed *et al*., 2026).

Further work should therefore be committed to developing methods that enhance prioritization, quantification and hypothesis testing at the *variant*-level. In this sense, we should also acknowledge challenges in the validation of conjunctional FDR-significant loci due to the lack of further SNP-wise statistics, such as beta coefficients or standard errors, derived from this method (Sha *et al*., 2025). To that extent, despite their own limitations, traditional approaches such as genetic correlations confer relevant complementary insights into the statistical significance of two trait’s genetic commonalities, at least, when unidirectionally associated. Together with conjunctional FDR, this approach has been importantly limited in our work by the unavailability of summary statistics for globularity (Molz *et al*., 2024). Hence, expanding and validating associations indirectly proposed for globularity (e.g., on the grounds of links to ventricular volume and glymphatic function measures) stays a prospect to accomplish once summary statistics for the globular brain shape GWAS become available.

Beyond constraints in assessing the extent and direction of genetic overlap at the variant-level, limitations of our research arise more broadly from its reliance on variably powered GWAS. While our methods (e.g., hypergeometric tests) partly account for disparity in the number of genes and variants pertaining to each trait, there is still the possibility that variance in the strength of the associations reported reflects influences of ancestry, sample size or subject age in each GWAS, rather than variability in our phenotypes’ actual biological relations to globularity. Moreover, standing on multiple genetic studies to enhance our understanding of the globular brain shape, our work contextualizes the polygenic bases of this trait in a purely relational manner, thus falling short in supplying empirical evidence for causal or temporal connections suggested. This may serve to underscore the hypothetical nature of many of the mechanistic interpretations discussed, but also to highlight the avenues opened for subsequent experimental inquiries by the articulation of such hypotheses. Moreover, it may serve to point to the multiple testing burden potentially impacting our inspection: applying Benjamini-Hochberg correction to individual tests, our work may suffer from a global false-discovery issue driven by the cumulative number of statistical comparisons run through it.

Finally, apart from aspects that remain unaddressed in our research (e.g., cell type- or tissue-specificity as a relevant contextual driver of trait-specificity beyond brain areas or developmental periods targeted by gene expression), it cannot be stressed enough that our work paradoxically uses genotype-phenotype associations in primarily adult, modern human, European populations as a proxy to reach developmental and evolutionary inferences about a *Homo sapiens*-universal trait. The need to assess the generalizability of our findings in more diverse populations, both in terms of age and ancestry, must therefore be emphasized, just as it is necessary to further validate the evolutionary reliability of the signals uncovered as a window into the emergence of the *sapiens*-distinctive globular head shape.

## Methods

We based our research on publicly available data (cf. Data availability), mainly, from genomic, proteomic and metabolomic association studies. Given previously noticed differences in the degree of specificity to which GWAS-prioritized *genes* and *variants* may reflect trait biology (Spence *et al*., 2026), we extracted information at both levels (STX) to form our pool of datasets. At the *gene* level, HGNC gene symbols or Ensembl Gene IDs were obtained from GWAS data; through the *biomaRt* R package (version 3.23; Smedley *et al*., 2009; https://doi.org/doi:10.18129/B9.bioc.biomaRt), correspondences between both types of identifiers were attained. To maintain consistency in the strategies defining our trait-associated gene sets, retrieval of significant gene lists from MAGMA gene-based analyses was prioritized. However, due to methodological variability in our GWAS of interest, genes derived using alternative approaches (e.g., FUMA annotations, GWGAS-reported genes) were also included when necessary. At the *variant-*level, either lead SNPs (with statistically most significant trait-associations within a chromosomal region) or all independent significant SNPs were considered, depending on availability. For effect inspections, significant variants from conjunctional FDR (cf. below) were incorporated as well. Information extracted encompassed *rsIDs* (for hypergeometric testing) and, when accessible, effect and non-effect alleles, effect sizes (*beta*), effect allele frequencies, standard errors, sample sizes and *p* values, as required by the Two Sample MR package (Hemani *et al*., 2017, 2018; https://mrcieu.github.io/TwoSampleMR/index.html) to harmonize data for effect comparisons. To maximize the number of datasets available for this purpose, the following procedures were applied to achieve reliable estimations in the event of missing values:

a. retrieval of allele frequencies using the Ensembl VEP tool (1000 Genomes data; web interface: https://www.ensembl.org/Tools/VEP),
b. estimation of standard errors from effect allele frequencies, sample sizes and z-scored effect values: 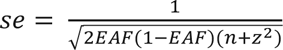
c. estimation of *beta* effect sizes from *z*-scored effects and standard errors (*beta* = *z * se*).

Overall, we established a pool of genetic association studies linked to (i) *31 traits for analyses of gene-level overlaps (hypergeometric tests) and transcriptomic profiles, (ii) 29 traits for inspections of variant-level overlaps (hypergeometric tests), and (iii) 50 traits for conjunctional FDR and genetic correlations* (Suppl. Tables 1-3). Intersections with a catalog of high-frequency single nucleotide changes considered variants derived from both conjunctional FDR and our initial GWAS sets undergoing hypergeometric testing.

Additionally, for transcriptomic inspections, we complemented GWAS-retrieved data with publicly available gene lists pertaining to 23 energy metabolism pathways from Pourmajidian *et al*.’s (2026) GitHub repository (accessible through https://github.com/netneurolab/pourmajidian_metabolism-genes/). As indicated by Pourmajidian *et al*. (2026), the downloaded gene sets were curated using Gene Ontology (GO) (Ashburner *et al*., 2000) and Reactome Knowledge base (Croft *et al*., 2014), databases which provide a unified platform for evidence-based and cross-referenced gene functional classification and pathway annotation. Concretely, for each of the 23 pathways researched, corresponding GO biological processes (Ashburner *et al*., 2000) and Reactome pathway IDs (Croft *et al*., 2014) were detected, such that gene sets involved in each pathway could be extracted using *biomaRt* (https://doi.org/doi:10.18129/B9.bioc.biomaRt; Smedley *et al*., 2009) and *GO.db* packages (https://bioconductor.org/packages/GO.db/). The curated energy metabolism gene sets employed in our research eventually comprised genes consistently annotated in both databases.

### Gene expression data

Lifespan and cortex-wise transcriptomic datasets were deployed to define the spatiotemporal patterns of expression linked to our GWAS and metabolic pathway gene sets.

In particular, for our temporal transcriptomic analysis, we leveraged publicly available RNA-sequencing data from BrainSpan (https://www.brainspan.org/static/download.html; Li *et al*., 2018). Using conditional quantile normalization (*Gencode v10 summarised to genes*), we pre-processed it into normalized Reads Per Kilobase of transcript per Million (RPKM) (Li *et al*., 2018), from which an expression matrix with normalized RPKM values for 52,376 genes across 524 samples resulted. We further cleaned the sample-by-gene expression matrix by (a) retaining regions with at least 1 sample in each age group, (b) removing duplicate genes, and (c) maintaining genes with an RPKM value ≥ 1 in 80% of the samples at each spatiotemporal point (Parikshak *et al*., 2013; Pourmajidian *et al*., 2026). Step (b) reduced data to 47,808 genes, whereas (c) yielded a final dataset containing expression values for 7,923 genes in 498 samples and 16 unique brain regions, including: amygdaloid complex, posterior (caudal) superior temporal cortex (area 22c), anterior (rostral) cingulate (medial prefrontal) cortex, dorsolateral prefrontal cortex, orbital frontal cortex, inferolateral temporal cortex (area TEv, area 20), hippocampus, ventrolateral prefrontal cortex, primary auditory cortex, primary visual cortex (striate cortex, area V1/17), striatum, primary motor cortex (area M1, area 4), posteroventral (inferior) parietal cortex, primary somatosensory cortex (area S1, areas 3, 1, 2), cerebellar cortex, and mediodorsal nucleus of thalamus.

Temporal expression values were subsequently log_2_ transformed and normalized in compliance with the upper quartile (UC) method (Bullard *et al*., 2010; Dillies *et al*., 2013; Miller *et al*., 2014). Scaled by their 75^th^ percentile expression value, each donor’s data was multiplied by the average 75^th^ percentile value across all donors (Pourmajidian *et al*., 2026). From the derived upper quartile-normalized data, trait- and pathway-specific lifespan transcriptomic trajectories were extracted by averaging expression values of genes within GWAS and metabolic pathway gene sets across Li *et al*.’s (2018) 31 age periods: 8 pcw, 9 pcw, 12 pc2, 13 pcw, 16 pcw, 17 pcw, 19 pcw, 21 pcw, 24 pcw, 25 pcw, 26 pcw, 35 pcw, 37 pcw, 4 months, 10 months, 1 year, 2 years, 3 years, 4 years, 8 years, 11 years, 13 years, 15 years, 18 years, 19 years, 21 years, 23 years, 30 years, 36 years, 37 years, 40 years. Because of constraints in the genes represented in the BrainSpan dataset or overcoming our quality controls, the final number of genes for which expression trajectories could be extracted in relation to each phenotype or pathway may diverge from the original input lists. In that sense, lists with less than four genes represented in BrainSpan were excluded from the analysis, such that 30 gene sets were eventually subjected to our inspection. For plotting purposes, the LOESS method was used to produce smoothed curves, whereas correlations were based on mean expression values, first averaged across donors and then across brain structures, at each lifespan period (cf. below).

As for our spatial transcriptomic exploration, regional microarray expression data were obtained from 6 post-mortem brains (1 female, ages 24.0-57.0 ± 13.38) provided by the Allen Human Brain Atlas (AHBA, https://human.brain-map.org; Hawrylycz *et al*., 2012). Data were processed with the *abagen* toolbox (version 0.1.3; https://github.com/rmarkello/abagen; Markello *et al*., 2021), using Schaefer *et al*.’s (2018) 400-region volumetric atlas in MNI space for parcellation.

Microarray probes were reannotated using data provided by Arnatkeviciute *et al*. (2019); probes not matched to a valid Entrez ID were discarded. Next, probes were filtered based on their expression intensity relative to background noise (Quackenbush, 2002), such that probes with intensity less than the background in ≥ 50.00% of samples across donors were discarded, yielding 31,729 probes. When multiple probes indexed the expression of the same gene, we selected and used the probe with the most consistent pattern of regional variation across donors (i.e., differential stability; Hawrylycz *et al*., 2015). This was calculated with 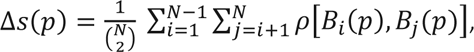 where *rho* (*ρ*) is Spearman’s rank correlation of the expression of a single probe, *p*, across regions in two donors *B_i_* and *B*_j_, and N is the total number of donors. Here, regions correspond to the structural designations provided in the ontology from the AHBA.

The MNI coordinates of tissue samples were updated to those generated via non-linear registration using the Advanced Normalization Tools (ANTs; https://github.com/chrisfilo/alleninf). To increase spatial coverage, tissue samples in the left hemisphere were mirrored into the right hemisphere (Romero-Garcia *et al*., 2018). Samples were assigned to brain regions in the provided atlas if their MNI coordinates were within 2 mm of a given parcel. To reduce the potential for misassignment, sample-to-region matching was constrained by hemisphere and gross structural divisions (i.e., cortex, subcortex/brainstem, and cerebellum, such that e.g., a sample in the left cortex could only be assigned to an atlas parcel in the left cortex; Arnatkeviciute *et al*., 2019). If a brain region was not assigned a tissue sample based on the above procedure, every voxel in the region was mapped to the nearest tissue sample from the donor in order to generate a dense, interpolated expression map. The average of these expression values was taken across all voxels in the region, weighted by the distance between each voxel and the sample mapped to it, in order to estimate the parcellated expression values for the missing region.

Inter-subject variation was addressed by normalizing tissue sample expression values across genes using a robust sigmoid function (Fulcher *et al*., 2013): 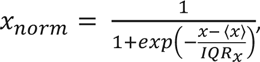 where 〈x〉 is the median and IQR_x_ is the normalized interquartile range of the expression of a single tissue sample across genes. Normalized expression values were then rescaled to the unit interval: 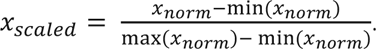 Gene expression values were then normalized across tissue samples using an identical procedure. All available tissue samples were used in the normalization process regardless of whether they were assigned to a brain region. Tissue samples not matched to a brain region were discarded after normalization. Samples assigned to the same brain region were averaged separately for each donor and then across donors, yielding a regional expression matrix with 400 rows, corresponding to brain regions, and 15,677 columns, corresponding to the retained genes.

GWAS and metabolic pathway gene sets were then employed to retrieve trait- or pathway-specific gene expression matrices. Given limitations in the genes present in the AHBA dataset and passing *abagen* quality controls, the number of genes for each phenotype or pathway may diverge from the original input lists. In that sense, lists with less than four genes represented in the Allen Human Brain Atlas were excluded from the analysis, such that 30 gene sets were eventually subjected to our inspection. Spatial expression values averaged across all genes within each resulting expression matrix were used to generate trait- or pathway-specific brain maps with the *surfplot* package (Gale *et al*., 2021; Vos de Wael *et al*., 2020), as well as to perform spatial correlation analyses via spatial permutation tests.

### The archetypal sensorimotor-association axis

To discover meaningful axes of cortical organization along which our *variant effect* and *gene expression* data may be arranged, we capitalized on 11 cortical gradients that have been proposed to manifest 10 long-range features defining the cortex along evolutionary, anatomical and functional hierarchies, as well as their integration into a unified archetypal sensorimotor-association axis (Sydnor *et al*., 2021).

Values corresponding to each gradient were extracted from Sydnor *et al*.’s (2021) Archetypal Sensorimotor-Association Axis GitHub repository (accessible at https://github.com/PennLINC/S-A_ArchetypalAxis). Concretely, for spatial correlations with our transcriptomic brain maps, data already parcellated according to Schaefer *et al*.’s (2018) *local-global* atlas (400 parcels – 17 Networks) was downloaded (https://github.com/PennLINC/S-A_ArchetypalAxis/tree/main/Schaefer400_17Network), as this directly matched the regional divisions and coordinate space (MNI) employed to process and obtain our cortex-wise gene expression distributions. Instead, in the absence of data directly corresponding to the cortical parcellation of the BrainVISA atlas (Perrot *et al*., 2011) or its original coordinate space (MNI), values supplied in *fsLR* (HCP) surface vertices (Van Essen *et al*., 2012) were extracted (https://github.com/PennLINC/S-A_ArchetypalAxis/tree/main/FSLRVertex) to perform spatial correlations with our measures of sulcal depth and width concordances (converted from MNI to *fsLR* space).

### Evolutionary annotations

To identify modern human derived high-frequency (≥ 90%) variants in our pool of GWAS, we took advantage of a publicly available dataset that capitalizes on Neanderthal and Denisovan genomes to compile a genome-wide catalogue of single nucleotide changes distinguishing modern humans from archaic hominins (Kuhlwilm & Boeckx, 2019) (accessible at https://doi.org/10.6084/m9.figshare.8184038). As indicated in the original paper, this catalog comprises “archaic”-specific variants as well as all varying loci within modern human populations. It derives frequencies from 1000 Genomes Project and ExAc data, and it uses the human genome version *hg19* as reference. Beyond the full catalogue, a subset of data is provided including derived variants at a ≥ 90% global frequency cutoff (Kuhlwilm & Boeckx, 2019). Variants we report correspond to intersections between our lists of GWAS- or conjunctional FDR-significant SNPs and subsets of the catalog encompassing “archaic” (Neanderthal/Denisovan) variants and high-frequency “modern” (*sapiens*) variants.

As for the temporal mapping of both kinds of single nucleotide changes (“modern” and “archaic”), it was based on a dataset compiling times of emergence for the catalog’s (Kuhlwilm & Boeckx, 2019) variants, as predicted by the Genealogical Estimation of Variant Age (GEVA) tool (Albers & McVean, 2020; Andirkó *et al*., 2022). This tool deploys a hidden Markov model approach to infer the location of ancestral haplotypes relative to a given variant, then inferring time to the most recent ancestor in pairwise comparisons by coalescent-based clock models. From a combination of the resulting pairwise information, a posterior probability measure of variant age is derived. Thus, the annotations we used for temporal mapping consist of dating information for the alleles in Kuhlwilm & Boeckx’s (2019) catalog, as extracted from the bulk summary information of GEVA age predictions (Andirkó *et al*., 2022).

### Brain atlases

To achieve spatial accounts for the *variant effect* and *gene expression* patterns in our data, we leveraged several publicly available brain atlases.

Namely, for white matter *edge-* and *node-*level connectivity data (Sha *et al*., 2023), we made use of the Automated Anatomical Labelling Atlas (Tzourio-Mazoyer *et al*., 2002; https://www.gin.cnrs.fr/en/tools/aal/), which consists of 45 regions per hemisphere comprising both cerebral cortical and subcortical structures. Its total 90 regions constitute the *nodes* for which Sha *et al*. (2023) supply variant effect information. Regarding data for *edge*-level connections, it was acquired by Sha *et al*. (2023) via deterministic tractography procedures applied to quality-controlled diffusion MRI (dMRI) data obtained from Siemens Skyra 3 T scanners, generating tens of thousands of streamlines to reconstruct individual white matter connectivity matrices on the grounds of the AAL atlas (Tzourio-Mazoyer *et al*., 2002). The atlas in question is originally in MNI coordinate space.

For *sulcal morphology* effects, we took advantage of the BrainVISA Sulci atlas (BSA, Perrot *et al*., 2011). This was originally a probabilistic atlas based on a coherent Bayesian framework that allows automatic recognition and labelling of the cortex’s folding patterns (sulci). As a probabilistic 4-dimensional atlas, the initial NIFTI file downloaded in MNI coordinate space from https://brainvisa.info/web/morphologist.html provided, for each of its voxels, a numeric vector representing the probabilities that the voxel was assigned to each of the anatomical landmarks (brain structures/sulci) considered in the atlas. However, to be able to map variant effect sizes from Sun *et al*. (2022) and Lemaitre *et al*. (2023) to specific sulci, we converted the original atlas to a deterministic, 3-dimensional one by assigning each voxel to its highest-probability argument.

Finally, for our transcriptomic data, Schaefer *et al*.’s (2018) *local-global* parcellation of the brain’s cortex (https://github.com/ThomasYeoLab/CBIG/tree/master/stable_projects/brain_parcellation/Schaefer2018_LocalGlobal) was consulted. This division of the cerebral cortex uses task-fMRI and rs-fMRI to propose a gradient-weighted Markov Random Field (gwMRF) model integrating previous *local gradient* (detecting abrupt transitions in functional connectivity patterns) and *global similarity* (clustering similar functional connectivity patterns regardless of spatial proximity) approaches. Among the multiple resolution (100 to 1000 parcels) and coordinate space system (*fsaverage, fsLR 32k, MNI*) options at which the atlas is available, we retrieved a 400 parcellation in MNI space (1 mm resolution; 17 Networks) for our transcriptomic examinations.

Although initially in MNI coordinate system, both white matter connectivity values and data from the BrainVISA Sulci Atlas (Perrot *et al*., 2011) had to be converted to *fsLR 32k* space with neuromaps (Markello *et al*., 2022; Buckner *et al*., 2011; Wu *et al*., 2018), either to conduct correlations with Sydnor *et al*.’s (2021) sensorimotor-association axis or for the purpose of visualization with *nilearn* (https://doi.org/10.5281/zenodo.8397156). For cerebellar and subcortical structures, plotting could be achieved using the *subcortex visualization* package instead (python version; Bryant, 2026), whereas *cortex-wise* transcriptomic maps could be generated with the *surfplot* package (Gale *et al*., 2021; Vos de Wael *et al*., 2020).

### Intersection-based analyses

We first sought to determine the extent to which our traits of interest may be biologically associated. Using R software (Posit team, 2025), we performed pairwise intersections between our original lists of GWAS-prioritized genes and variants (31 and 29 sets, respectively). Initially, intersections allowed us to quantify the number of genes or variants in overlap between two traits, as shown in Suppl. Tables 4-5. Then, hypergeometric tests (*phyper*) were conducted for each overlap, employing the hypergeometric distribution function to compute the probability of having drawn an intersection of length *L* independently and without replacement from two distinct sets, A and B, composed of objects of *n* categories and pertaining to a global population or universe of size *N*, with *a* and *b* number of draws, respectively (Kalinka, 2013). Multiple hypothesis testing was corrected by means of the Benjamini-Hochberg (BH) procedure, such that overlaps with BH-adjusted *p* values under 0.05 were considered statistically significant.

For *gene-*level inspections, intersections and hypergeometric tests were conducted with Ensembl IDs and gene symbols, so as to control for existing variability in the correspondences between these two types of identifiers. In both cases, a universe of *N* = 20,781 (based on Wagstyl *et al*., 2024) was designated as a simplifying assumption for the total number of protein-coding genes from which our input sets and output intersections could be drawn. Since outcomes were notably similar across *Ensembl ID* and *gene symbol* analyses (Suppl. Table 6), we continued our explorations with the latter, reporting results for them.

As for *variant-*level assessments, they were grounded on extracted lists of rsIDs. On this occasion, a model universe of *N* = 8,000,000 variants was assumed on the basis of the total SNPs usually studied by GWAS in our pool, which tended to converge on values between 8 and 9 million variants (Suppl. Table 2).

We subsequently tried to understand how our genetic intersections were organized by detecting phenotype clusters or, especially, traits more closely related to globularity on the basis of *gene-* and *variant-*level overlaps. We therefore applied the *Louvain clustering* method for community detection (Blondel *et al*., 2008) to *gene-* and *variant-*level matrices of negative log10-transformed *p* values derived from our pairwise hypergeometric tests. This procedure, employed via *igraph* (Csárdi & Nepusz, 2006) in our research, allows reliable extraction of community structure within large networks through a heuristic strategy grounded on modularity optimization. Modularity constitutes a scalar value between −1 and 1 measuring the density of links inside communities as compared to links between communities, thus providing a measure of the quality of the clusters or partitions recognized. Negative modularity scores imply poor partitions, whereas scores ≥ 0.3 are deemed indicative of highly compartmentalized and structured networks with meaningful, distinguishable communities. In between, low positive modularity scores (0.0 – 0.3) suggest weak, rather overlapping community structures, with faint boundaries within highly integrated, broad networks.

Based on our intersection results, we ultimately wanted to comprehend whether the GWAS-significant variants encountered in our overlaps were more closely associated to trait-specificity in either of two ways: (a) by acting through trait-specific genes, or (b) by exerting context-specific influences on pleiotropic genes (Spence *et al*., 2026). Under (a), genes mapped to variants would be expected to be found less frequently across datasets linked to different phenotypes, given their trait-specificity; under (b), genes mapped to variants would be predicted to occur more frequently across lists of GWAS-prioritized genes associated to different traits, in light of their pleiotropy.

Hence, we considered that variations in the cross-dataset frequencies of genes mapped to variants in our intersections could provide a proximate measure for the degree of pleiotropy of such genes, and thus, the degree to which variants shared between globularity and brain-related phenotypes were trait-specific via *context-* or *gene-*specificity. To test this, frequencies of occurrence were first computed for all of the variants and genes in our data, such that every additional association of a variant or gene to a distinct phenotype in our pool of GWAS counted as an occurrence. Then, variants were mapped to genes using *gProfiler* (Kolberg *et al*., 2023; accessible at https://biit.cs.ut.ee/gprofiler/snpense), and to compare the frequencies of genes related to globularity-intersecting variants with respect to other genes in our pool, *z-scores* (*z = frequency - mean(frequency) / sd(frequency)*) were computed as an indicator of how many standard deviations our gene frequencies were from the mean across phenotypes (Suppl. Tables 8-9). Positive *z-scores* ≥ 2.0 were considered to indicate substantially higher frequencies than the mean.

### Variant effect inspections

Having identified asymmetries in globularity’s intersections with other brain-related phenotypes at the gene- and variant-levels, we went on to investigate the latter, given evidence that it guarantees greater trait-specificity (Spence *et al*., 2026). Particularly, we reasoned that maximizing our discovery of variants (likely) pleiotropically shared between globularity-related phenotypes, as well as contrasting their effects across traits, could offer us a window into both concordant and discordant dimensions of genetic overlap (Sha *et al*., 2025) with globularity, together with principles organizing them, at high degrees of resolution.

Under this rationale, conjunctional FDR analysis (Andreassen *et al*., 2013) was first conducted with the R package *cfdr.pleio* (Ploner & Frei, 2026) to inquire into shared genetic variants between different brain-related phenotypes. This strategy enhances the statistical power to discover pleiotropic loci between phenotypes by taking advantage of cross-trait SNP enrichment within an empirical Bayesian framework (Sha *et al*., 2025).

The method was applied to summary statistics of 50 phenotypes (Suppl. Table 3), comprising measures of glymphatic function, ventricular volumes, sulcal depth, cerebellar shape or brain activity networks, among others. At each iteration, estimates of genetic overlap were computed for a pair of traits *A* and *B.* Our code was programmed so that most of the 1,225 possible pairwise combinations of phenotypes were allowed to undergo the analysis; excluded combinations comprised those involving summary statistics from the same study (e.g., depth measures for sulci like the left intraparietal and the right superior frontal, from Lemaitre *et al*., 2023).

In total, 913 pairwise combinations of phenotypes (74.53%) underwent our conjunctional FDR analysis. For each pair of phenotypes *A* and *B*, conditional FDR re-ranked the test statistics of trait *A* conditioned on its association with trait *B* (i.e., FDR(*A*|*B*)), such that conditional FDR values concerning the significance of the association with trait *A* could be reached. Then, the same conditional FDR process was performed reversely (i.e., FDR(*B*|*A*)), so that the significance of SNP associations with trait *B* could be evaluated. From this, a conjunctional FDR value was determined as the maximum of the two conditional FDR values in both directions (FDR(*A*|*B*) and FDR(*B*|*A*)), thus providing a conservative estimate of the posterior probability that a genetic variant has no association with either phenotype, considering that its *p* values for association with both traits are as small(er) than the identified *p* values (Sha *et al*., 2025).

We deemed conjunctional FDR scores < 0.05 significant. From this, independent significant signals for each pair of phenotypes were derived by clumping filtered sets of conjunctional FDR-significant variants with PLINK (v. 1.9; Chang *et al*., 2015; https://www.cog-genomics.org/plink/). Concretely, linkage disequilibrium clumping was performed with the *1000 Genomes Project Phase 3* reference panel, defining index variants under a significance threshold of *p* < 0.05, and assigning to the same clump variants in LD (*r*^2^ ≥ 0.2) or within a 250-kb window with respect to the index variant, such that the most significantly associated variant within each LD block could be retained as the index SNP.

Pairwise comparisons of pleiotropic variant effect sizes were then carried out for datasets providing sufficient effect-related information for the purposes of harmonization and analysis. Researched datasets therefore included all of our 50 conjunctional FDR phenotypes, to the extent that they yielded genetic overlaps with other traits undergoing the inspection, and the following traits with genome-wide significant variants: *brain aging* (Kim *et al*., 2026), *white matter edge-* and *node-*level connectivity (Sha *et al*., 2023), *sulcal morphology* (Sun *et al*., 2022), *globularity* (Molz *et al*., 2024), *cortical thickness* and *grey-white contrast* (Kim *et al*., 2025), *idiopathic normal pressure hydrocephalus* (Räsänen *et al*., 2024), *Lewy body dementia* (Zhu *et al*., 2025), *cerebrospinal fluid proteome* (Western *et al*., 2024), *perivascular space burden* (Duperron *et al*., 2023), *language white matter connectome microstructure* (Mekki *et al*., 2022), and *cerebellar morphology* (Moberget *et al*., 2026). In total, more than 300 pairs of traits were assessed in terms of their shared variants’ impacts.

To attain sound comparisons for each pair of phenotypes, effects of overlapping variants were harmonized to the same reference allele with the *harmonise_dat* function of the TwoSampleMR package (Hemani *et al*., 2017, 2018). Deployed in its default mode (*action* = 2), this function allowed inferring ambiguous stands on the basis of allele frequency information when aligning effect sizes to a common allele. This boosted the number of variants that could be reliably harmonized, while dropping SNPs when their effects could not be properly aligned.

Once harmonization was completed, degrees of concordance or discordance between traits in each pair were calculated under several metrics. Namely, for each pleiotropic variant shared between two phenotypes, effect directions were compared, considering them concordant if both positive or negative, but discordant if opposed. From this, counts of concordances (where *concordant SNPs* = +1, vs. *discordant SNPs* = 0) and discordances (where *discordant SNPs* = +1, vs. *concordant SNPs* = 0) were obtained, as well as their proportions with respect to the total of variants in overlap between two traits. However, these quantifications were not fully apt to account for both congruent and discrepant dimensions of genetic overlap simultaneously, as they equated concordant or discordant effect directions to an absence of impacts (i.e., 0). To therefore achieve a better measure of the discordant and concordant influences possibly acting on two phenotypes, the former were counted as –1, whereas the latter were taken as +1, in line with Sha *et al*. (2025). In this way, both congruence counts (e.g., −1 discordant with +2 concordant = 1) and their corresponding ratios (e.g., 1/3 = 33.33%) were computed. While not exempt of limitations, this strategy permitted a continuous and integrated account of where greater degrees of convergence and divergence between two traits may lie, which was relevant for spatial inspections. Moreover, it proved to lend consistent results with separate quantifications of mean effect sizes, as well as separate measures of concordance and discordance counts or proportions (Suppl. Fig. 8).

Beyond concordance scores or ratios, this last quantification was also used for the performance of spatial correlations, looking at the distribution of convergent or divergent effects throughout the brain, and for correlations examining two phenotypes’ patterns of association (*concordance/discordance*) to other traits. Spearman was the method used in both cases (cf. below).

### Correlation-based analyses

To better understand the organization of our transcriptomic and variant effect data across the human cortex and lifespan, we performed spatial and temporal correlations with the Spearman method. Temporal correlations were based on the transcriptomic trajectories associated to 30 distinct gene lists, as derived by aggregating their corresponding gene expression values across 31 developmental periods in BrainSpan (8 pcw – 40 yrs). Averaging was first conducted across donors, and then across structures. As already indicated, datasets with insufficiently represented genes (<4) on BrainSpan were excluded from the analysis. To reduce the likelihood of inflated *p* values, permutations were used to compare the empirical results obtained with 10,000 null models, generated by drawing and correlating temporal expression profiles from random lists of genes of the same size and degree of overlap of the lists under inspection.

As for spatial correlations, *spatial permutation (spin) tests* (Alexander-Bloch *et al*., 2018; Váša *et al*., 2018; Burt *et al*., 2020) were used to control the inherent distance-dependent spatial autocorrelation of both structural and functional measures of the brain. Intrinsically, such spatial autocorrelation would make cortex-wise *gene expression* or *variant effect* data not independent and identically distributed —as commonly required for sound statistical inference—, but similar in nearby voxels or regions due to both biological and technical factors (Markello *et al*., 2021; Pourmajidian *et al*., 2026). Accounting for these spatial dependencies in brain data, *spin tests* permute voxels or parcels while maintaining the spatial structure and auto-correlation, hence allowing a more reliable framework for hypothesis testing. In light of this, the *netneurotools* package (https://netneurolab.github.io/netneurotools/) was used in *python* to generate and implement spatial permutation tests along the *Váša method* (Váša *et al*., 2018). This procedure produces rotations of parcellated brain data based on parcel centroid coordinates, ensuring that reassignments are not duplicated, while keeping the spatial covariance structure of the brain. Therefore, it offers a true null distribution, against which empirical brain data can be contrasted to test for the statistical significance of correlations. We did so across 10,000 iterations for *spin tests* generated for Schaefer *et al*.’s (2018) and Perrot *et al*.’s (2011) centroid coordinates, having previously converted spatial maps under inspection to a common coordinate system and parcellation. The non-parametric *p* values obtained as a measure of statistical significance are commonly referred to as *p*_spin_.

When spatial correlations seemed to unravel relevant clusters of phenotypes in our data, we resorted to the R package *pvclust* (Suzuki *et al*., 2006; https://github.com/shimo-lab/pvclust) to evaluate their strength. This package conducts hierarchical clustering and, for each cluster contained in the clustering of the original data, uses bootstrap resampling techniques to automatically calculate two probability (*p*) values: an *approximately unbiased* (AU) *p* value, and a *bootstrap probability* (BP) *p* value. Among these, AU is considered superior as an approximation to unbiased *p* value (Suzuki *et al*., 2006). Thus, clusters with AU values > 95% were judged strongly supported by our data.

### Evolutionary inspection

We finally took advantage of our realization of extensive variant-level pleiotropies engaging globularity to seek an evolutionary characterization of the emergence of the globular phenotype, framing it within the context of the evolution of multiple brain and cognitive traits. Our reasoning was that, given the observation of genetic interplays with globularity across multiple brain-related phenotypes, evidence of single nucleotide changes impacting brain and cognitive traits could indirectly provide insights into the evolutionary context that might have shaped the emergence of the globular brain shape.

Intersections were conducted between our lists of variants and Kuhlwilm & Boeckx’s (2019) subsets of “archaic” and “modern” single nucleotide changes. Since the size of the overlaps reached was not sufficiently large to yield significant hypergeometric testing results or to perform statistical inference analyses, variants were explored descriptively in terms of their effect sizes, phenotypic impacts and times of emergence. Our logic was that such information could also prove valuable for an understanding of evolutionary dynamics preceding and concurring with the emergence of the globular endocranium.

Effects were harmonized to the major allele, so as to understand phenotypic impacts fixed at high-frequency across modern human populations. Ratios of variants with positive effects with respect to the total were then computed for each phenotype’s overlap with the “modern” subset of Kuhlwilm & Boeckx’s (2019) catalog. As for high-frequency variants’ times of emergence, they were assessed using the aforementioned temporal annotations derived from GEVA. These allowed mapping most of the SNPs in our intersections (137/158 = 86.71%), except for 21 exceptional variants without an assigned date of emergence (Suppl. Table 34).

## Acknowledgments

We thank all researchers who made available the computational tools and genetic datasets on which this work relies. Amaia Carrión-Castillo provided constructive comments that helped shape the presentation of the research reported here. Figures were further edited and adapted using BioRender.

## Author contributions

Conceptualization: C. B., J. C. R.; methodology: C. B., J. C. R; data curation: J. C. R.; software: J. C. R.; formal analysis: J. C. R.; visualization: J. C. R.; investigation: C. B., J. C R.; writing – original draft: C. B., J. C. R; writing – review and editing: C. B., J. C. R; supervision: C. B.; funding acquisition: C. B.

## Data and code availability

All datasets used in this study are publicly available. Scripts and data used to perform the analyses will be made available at the article’s GitHub repository: https://github.com/jcriu/Globularity.

## Supplementary tables

**Supplementary Table 1.** Index of studies for genome-wide significant gene-level analyses.

**Supplementary Table 2.** Index of studies for genome-wide significant variant-level analyses.

**Supplementary Table 3.** Reference summary statistics for conjunctional false discovery rate (FDR) and genetic correlation analyses.

**Supplementary Table 4.** Number of genes in each pairwise intersection between datasets (top: gene symbols; bottom: Ensembl gene IDs).

**Supplementary Table 5.** Number of variants in each pairwise intersection between datasets.

**Supplementary Table 6.** BH-corrected *p* values for gene-level pairwise hypergeometric tests between datasets (top: gene symbols; bottom: Ensembl gene IDs).

**Supplementary Table 7.** BH-corrected p values for variant-level pairwise hypergeometric tests between datasets.

**Supplementary Table 8.** Cross-phenotype gene frequencies (*gene symbols*).

**Supplementary Table 9.** Cross-phenotype variant frequencies.

**Supplementary Table 10.** Effect relations for pleiotropic variants shared between Globularity and Ventricular Morphology 1.

**Supplementary Table 11.** Effect relations for pleiotropic variants shared between Globularity and Glymphatic DTI-ALPS 1.

**Supplementary Table 12.** Conjunctional FDR results for the four brain ventricles (Ventricular Morphology 1) and Glymphatic DTI-ALPS 1.

**Supplementary Table 13.** Spearman correlations between 26 brain-related phenotypes according to their patterns of concordance and discordance.

**Supplementary Table 14.** Genetic correlation results for the brain ventricles (Ventricular Morphology 1) and glymphatic system (Glymphatic DTI-ALPS 1).

**Supplementary Table 15.** Effect relations for pleiotropic variants shared between Globularity/Ventricular Morphology 1 and Cerebellum GM Volume Components.

**Supplementary Table 16.** Effect relations for pleiotropic variants shared between Globularity/Ventricular Morphology 1 and White Matter Microstructural Parameters.

**Supplementary Table 17.** Ratios of concordant edges across globularity and ventricular morphology (1) data for each node inspected.

**Supplementary Table 18.** Ratios of concordant variants across globularity and ventricular morphology (1) data for node-level connectivity effects.

**Supplementary Table 19.** Effect relations for pleiotropic variants shared between Globularity/Ventricular Morphology 1 and Sulcal depth.

**Supplementary Table 20.** Effect relations for pleiotropic variants shared between Globularity/Ventricular Morphology-1 and Sulcal width.

**Supplementary Table 21.** Effect relations for variants in overlap between Grey-White Contrast / Cortical Thickness / Local Brain Aging and Glymphatic DTI-ALPS 1 / Ventricular Morphology 1 / Globularity

**Supplementary Table 22.** Effect relations for pleiotropic variants shared between Local Brain Aging/Grey-White Contrast/Cortical Thickness and Sulcal width.

**Supplementary Table 23.** Pairwise spatial correlations between ventricular volumes, globularity, glymphatic function, grey matter-white matter contrast, local brain aging and cortical thickness according to their patterns of concordance with sulcal width.

**Supplementary Table 24.** Spatial correlations between the sensorimotor-association axis features and the distribution of sulcal width concordances related to ventricular volumes, globularity, glymphatic function (DTI-ALPS 1), grey matter-white matter contrast, local brain aging and cortical thickness.

**Supplementary Table 25.** Pairwise correlations between traits according to their genes’ spatial patterns of expression.

**Supplementary Table 26.** Correlations between features in the sensorimotor-association axis and traits in our GWAS pool according to their genes’ spatial patterns of expression.

**Supplementary Table 27.** Pairwise correlations between traits according to their genes’ temporal patterns of expression.

**Supplementary Table 28.** Correlations between metabolic pathways and traits in our GWAS pool according to their genes’ spatial patterns of expression.

**Supplementary Table 29.** Correlations between metabolic pathways and traits in our GWAS pool according to their genes’ temporal patterns of expression.

**Supplementary Table 30.** Summary of overlaps with Kuhlwilm & Boeckx’s (2019) catalog of high-frequency single-nucleotide changes distinguishing modern humans from archaic hominins.

**Supplementary Table 31.** Modern and archaic variants (Kuhlwilm & Boeckx, 2019) intersecting with network measures of brain activity.

**Supplementary Table 32.** Modern and archaic variants (Kuhlwilm & Boeckx, 2019) intersecting with our initial pool of genetic studies.

**Supplementary Table 33.** Modern and archaic variants (Kuhlwilm & Boeckx, 2019) intersecting with our conjunctional FDR-significant SNPs.

**Supplementary Table 34.** Temporal mapping of variants in overlap with Kuhlwilm & Boeckx’s (2019) catalog.

**Supplementary Table 35.** Effect relations for pleiotropic variants shared between the CSF proteome and Cerebellum GM Volume Components.

**Supplementary Table 36.** Variants within the 8p23.1 cytogenetic region commonly significant for ASD and either Globularity or the Fourth Ventricle.

## Supplementary figures

**Supplementary Figure 1.** Effect direction relations for each cerebellar component and the average across components with respect to globularity, ventricular morphology, and glymphatic function measures.

**Supplementary Figure 2.** Concordance scores for Mekki’s (2022) white matter microstructural parameters, by tract.

**Supplementary Figure 3.** Pairwise correlations between traits according to their genes’ spatial patterns of expression.

**Supplementary Figure 4.** Correlations between our datasets’ spatial patterns of gene expression and the sensorimotor association axis components (Sydnor et al., 2021).

**Supplementary Figure 5.** Datasets significantly correlated with Sydnor *et al*.’s (2021) sensorimotor association axis components are significantly split into a sensorimotor and an association cluster through multiscale bootstrap resampling with 5000 replicates via *pvclust*.

**Supplementary Figure 6.** Temporal trajectories of gene expression associated to globularity and brain-related traits.

**Supplementary Figure 7.** Complete temporal mapping of the times of emergence associated with variants from Kuhlwilm & Boeckx’s (2019) catalog intersecting with our datasets.

**Supplementary Figure 8.** All (concordance/discordance) counts lend consistent results with other measures of convergence and divergence between traits, as reflected by the relations between the glymphatic system’s patterns of concordance with sulcal width and Sydnor et al.’s (2021) SMA axis features.

## References

Adolphs R. (2010). What does the amygdala contribute to social cognition?. Annals of the New York Academy of Sciences, 1191(1), 42–61. 10.1111/j.1749-6632.2010.05445.x

Ahmed, O., Saravanan, N., Rovsing, A., Simpson, D., Devarajan, A., Gunn, S., Singh, T., Lappalainen, T., Snjana, N. Current challenges in GWAS integration and fine-mapping for variant interpretation. bioRxiv. 10.64898/2026.07.04.736511

Albers, P. K., & McVean, G. (2020). Dating genomic variants and shared ancestry in population-scale sequencing data. PLoS biology, 18(1), e3000586. 10.1371/journal.pbio.3000586

Alexander-Bloch, A. F., Shou, H., Liu, S., Satterthwaite, T. D., Glahn, D. C., Shinohara, R. T., Vandekar, S. N., & Raznahan, A. (2018). On testing for spatial correspondence between maps of human brain structure and function. NeuroImage, 178, 540–551. 10.1016/j.neuroimage.2018.05.070

Amelink, J. S., Soheili-Nezhad, S., Alagöz, G., Llera, A., Schijven, D., Wang, M.-Y., Haak, K., Fisher, S. E., Beckmann, C. F., Francks, C. Integrating brain structure and function for the neurobiology and genetics of language. bioRxiv. 10.64898/2025.12.17.694832

Andirkó, A., Moriano, J., Vitriolo, A., Kuhlwilm, M., Testa, G., & Boeckx, C. (2022). Temporal mapping of derived high-frequency gene variants supports the mosaic nature of the evolution of Homo sapiens. Scientific reports, 12(1), 9937. 10.1038/s41598-022-13589-0

Andreassen, O. A., Thompson, W. K., Schork, A. J., Ripke, S., Mattingsdal, M., Kelsoe, J. R., Kendler, K. S., O’Donovan, M. C., Rujescu, D., Werge, T., Sklar, P., Psychiatric Genomics Consortium (PGC), Bipolar Disorder and Schizophrenia Working Groups, Roddey, J. C., Chen, C. H., McEvoy, L., Desikan, R. S., Djurovic, S., & Dale, A. M. (2013). Improved detection of common variants associated with schizophrenia and bipolar disorder using pleiotropy-informed conditional false discovery rate. PLoS genetics, 9(4), e1003455. 10.1371/journal.pgen.1003455

Apostolova, L. G., Green, A. E., Babakchanian, S., Hwang, K. S., Chou, Y. Y., Toga, A. W., & Thompson, P. M. (2012). Hippocampal atrophy and ventricular enlargement in normal aging, mild cognitive impairment (MCI), and Alzheimer Disease. Alzheimer disease and associated disorders, 26(1), 17–27. 10.1097/WAD.0b013e3182163b62

Arnatkeviciute, A., Fulcher, B. D., & Fornito, A. (2019). A practical guide to linking brain-wide gene expression and neuroimaging data. NeuroImage, 189, 353–367. 10.1016/j.neuroimage.2019.01.011

Ashburner, M., Ball, C. A., Blake, J. A., Botstein, D., Butler, H., Cherry, J. M., Davis, A. P., Dolinski, K., Dwight, S. S., Eppig, J. T., Harris, M. A., Hill, D. P., Issel-Tarver, L., Kasarskis, A., Lewis, S., Matese, J. C., Richardson, J. E., Ringwald, M., Rubin, G. M., & Sherlock, G. (2000). Gene ontology: tool for the unification of biology. The Gene Ontology Consortium. Nature genetics, 25(1), 25–29. 10.1038/75556

Balzeau, A., Bardinet, E., Bardo, A., Bernat, A-L., Didier, M., Filippo, A., Gilissen, E., Giolland, V., Gómez-Olivencia, A., Gómez-Robles, A., Heuzé, Y., Hublin, J-J., Hui, J., Kubicka, A. M., Labra, N., Leprince, Y., Mangin, J.-F., Masi, S., Pouydebat, E., Prima, S., Rivière, D. (2025). The brain of Homo neanderthalensis. 15th Annual Meeting of the European Society for the study of Human Evolution. Hal Science. https://hal.science/hal-05491688v1

Barger, N., Stefanacci, L., & Semendeferi, K. (2007). A comparative volumetric analysis of the amygdaloid complex and basolateral division in the human and ape brain. American journal of physical anthropology, 134(3), 392–403. 10.1002/ajpa.20684

Barron, S. A., Jacobs, L., & Kinkel, W. R. (1976). Changes in size of normal lateral ventricles during aging determined by computerized tomography. Neurology, 26(11), 1011–1013. 10.1212/wnl.26.11.1011

Barton R. A. (2007). Evolutionary specialization in mammalian cortical structure. Journal of evolutionary biology, 20(4), 1504–1511. 10.1111/j.1420-9101.2007.01330.x

Barton, R. A., & Venditti, C. (2014). Rapid evolution of the cerebellum in humans and other great apes. Current biology : CB, 24(20), 2440–2444. 10.1016/j.cub.2014.08.056

Baum, G. L., Cui, Z., Roalf, D. R., Ciric, R., Betzel, R. F., Larsen, B., Cieslak, M., Cook, P. A., Xia, C. H., Moore, T. M., Ruparel, K., Oathes, D. J., Alexander-Bloch, A. F., Shinohara, R. T., Raznahan, A., Gur, R. E., Gur, R. C., Bassett, D. S., & Satterthwaite, T. D. (2020). Development of structure-function coupling in human brain networks during youth. Proceedings of the National Academy of Sciences of the United States of America, 117(1), 771–778. 10.1073/pnas.1912034117

Bell, S., Tozer, D. J., & Markus, H. S. (2022). Genome-wide association study of the human brain functional connectome reveals strong vascular component underlying global network efficiency. Scientific reports, 12(1), 14938. 10.1038/s41598-022-19106-7

Benito-Kwiecinski, S., Giandomenico, S. L., Sutcliffe, M., Riis, E. S., Freire-Pritchett, P., Kelava, I., Wunderlich, S., Martin, U., Wray, G. A., McDole, K., & Lancaster, M. A. (2021). An early cell shape transition drives evolutionary expansion of the human forebrain. Cell, 184(8), 2084–2102.e19. 10.1016/j.cell.2021.02.050

Bennett, E. A., & Fu, Q. (2024). Ancient genomes and the evolutionary path of modern humans. Cell, 187(5), 1042–1046. 10.1016/j.cell.2024.01.047

Blondel, V., Guillaume, J.-L., Lambiotte, R., Lefebvre, E. (2008). Fast unfolding of communities in large networks. 10.48550/arXiv.0803.0476

Boeckx C. (2013). Biolinguistics: forays into human cognitive biology. Journal of anthropological sciences = Rivista di antropologia: JASS, 91, 63–89. 10.4436/jass.91009

Boeckx C. (2017). The language-ready head: Evolutionary considerations. Psychonomic bulletin & review, 24(1), 194–199. 10.3758/s13423-016-1087-5

Boeckx, C., & Benítez-Burraco, A. (2014a). Globularity and language-readiness: generating new predictions by expanding the set of genes of interest. Frontiers in psychology, 5, 1324. 10.3389/fpsyg.2014.01324

Boeckx, C., & Benítez-Burraco, A. (2014b). The shape of the human language-ready brain. Frontiers in psychology, 5, 282. 10.3389/fpsyg.2014.00282

Bonfanti L. (2006). PSA-NCAM in mammalian structural plasticity and neurogenesis. Progress in neurobiology, 80(3), 129–164. 10.1016/j.pneurobio.2006.08.003

Bruner E. (2004). Geometric morphometrics and paleoneurology: brain shape evolution in the genus Homo. Journal of human evolution, 47(5), 279–303. 10.1016/j.jhevol.2004.03.009

Bruner, E. (2010). Morphological differences in the parietal lobes within the human genus: a neurofunctional perspective. Current Anthropology, 51(S1), S77–S88. 10.1086/650729

Bruner, E., & Gallareto-Sande, R. (2025). Growth and development of the precuneus in humans from birth to early adulthood. Cerebral cortex (New York, N.Y. : 1991), 35(12), bhaf305. 10.1093/cercor/bhaf305

Bruner, E., & Jacobs, H. I. (2013). Alzheimer’s disease: the downside of a highly evolved parietal lobe?. Journal of Alzheimer’s disease: JAD, 35(2), 227–240. 10.3233/JAD-122299

Bruner, E., Battaglia-Mayer, A., & Caminiti, R. (2023). The parietal lobe evolution and the emergence of material culture in the human genus. Brain structure & function, 228(1), 145–167. 10.1007/s00429-022-02487-w

Bruner, E., Gallareto-Sande, R., López-González, F. J., Zhang, L., Sánchez Martín, C., Grothe, M. J., & Sánchez-Juan, P. (2026). Morphological variations and cortical atrophy of the precuneus in normal aging and Alzheimer’s disease. Journal of anatomy, 248(1), 62–70. 10.1111/joa.70027

Bruner, E., Preuss, T. M., Chen, X., & Rilling, J. K. (2017). Evidence for expansion of the precuneus in human evolution. Brain structure & function, 222(2), 1053–1060. 10.1007/s00429-015-1172-y

Bruner, E., Rangel de Lázaro, G., de la Cuétara, J. M., Martín-Loeches, M., Colom, R., & Jacobs, H. I. (2014). Midsagittal brain variation and MRI shape analysis of the precuneus in adult individuals. Journal of anatomy, 224(4), 367–376. 10.1111/joa.12155

Bryant, A. (2026). Subcortex visualization: A toolbox for custom data visualization in the subcortex and cerebellum. bioRxiv. 10.64898/2026.01.23.699785

Buckner, R. L., Krienen, F. M., Castellanos, A., Diaz, J. C., & Yeo, B. T. (2011). The organization of the human cerebellum estimated by intrinsic functional connectivity. Journal of neurophysiology, 106(5), 2322–2345. 10.1152/jn.00339.2011

Bufill, E., Agustí, J., & Blesa, R. (2011). Human neoteny revisited: The case of synaptic plasticity. American journal of human biology: the official journal of the Human Biology Council, 23(6), 729–739. 10.1002/ajhb.21225

Buisan, R., Moriano, J., Andirkó, A., & Boeckx, C. (2022). A Brain Region-Specific Expression Profile for Genes Within Large Introgression Deserts and Under Positive Selection in *Homo sapiens*. Frontiers in cell and developmental biology, 10, 824740. 10.3389/fcell.2022.824740

Bullard, J. H., Purdom, E., Hansen, K. D., & Dudoit, S. (2010). Evaluation of statistical methods for normalization and differential expression in mRNA-Seq experiments. BMC bioinformatics, 11, 94. 10.1186/1471-2105-11-94

Camandola, S., & Mattson, M. P. (2017). Brain metabolism in health, aging, and neurodegeneration. The EMBO journal, 36(11), 1474–1492. 10.15252/embj.201695810

Caporale, N., Leonardi, O., Villa, C. E., Vitriolo, A., Boeckx, C., & Testa, G. (2025). Tile by tile: capturing the evolutionary mosaic of human conditions. Current opinion in genetics & development, 90, 102297. 10.1016/j.gde.2024.102297

Caro, I., Western, D., Namba, S., Sun, N., Kawaguchi, S., He, Y., Fujita, M., Roshchupkin, G., D’Aoust, T., Duperron, M. G., Sargurupremraj, M., Tsuchida, A., Koido, M., Ahmadi, M., Yang, C., Timsina, J., Ibanez, L., Matsuda, K., Suzuki, Y., Oda, Y., … Debette, S. (2025). Proteogenomics in cerebrospinal fluid and plasma reveals new biological fingerprint of cerebral small vessel disease. Nature aging, 5(12), 2514–2531. 10.1038/s43587-025-01006-w

Casey, B. J., Tottenham, N., Liston, C., & Durston, S. (2005). Imaging the developing brain: what have we learned about cognitive development?. Trends in cognitive sciences, 9(3), 104–110. 10.1016/j.tics.2005.01.011

Casto, C., Poliak, M., Tuckute, G., Small, H., Sherlock, P., Wolna, A., Lipkin, B., D’Mello, A. M., & Fedorenko, E. (2026). The cerebellar components of the human language network. Neuron, 114(8), 1504–1523.e11. 10.1016/j.neuron.2025.12.030

Chang, C. C., Chow, C. C., Tellier, L. C., Vattikuti, S., Purcell, S. M., & Lee, J. J. (2015). Second-generation PLINK: rising to the challenge of larger and richer datasets. GigaScience, 4, 7. 10.1186/s13742-015-0047-8

Cerrito, P., & Burkart, J. M. (2023). Human Amygdala Volumetric Patterns Convergently Evolved in Cooperatively Breeding and Domesticated Species. Human nature (Hawthorne, N.Y.), 34(3), 501–511. 10.1007/s12110-023-09461-3

Cerrito, P., Gascon, E., Roberts, A. C., Sawiak, S. J., & Burkart, J. M. (2024). Neurodevelopmental timing and socio-cognitive development in a prosocial cooperatively breeding primate (*Callithrix jacchus*). Science advances, 10(44), eado3486. 10.1126/sciadv.ado3486

Chen, Z., Boehnke, M., Wen, X., & Mukherjee, B. (2021). Revisiting the genome-wide significance threshold for common variant GWAS. G3 (Bethesda, Md.), 11(2), jkaa056. 10.1093/g3journal/jkaa056

Chen, Z., Chen, Z., Mo, J., Chen, Y., Chen, L., & Deng, C. (2025). m6A RNA methylation modulates autophagy by targeting Map1lc3b in bisphenol A induced Leydig cell dysfunction. Journal of hazardous materials, 485, 136748. 10.1016/j.jhazmat.2024.136748

Ciuba, K., Figiel, I., Duński, E., Virmani, G., Dehingia, B., Abbas, M., Hemmatvand, K., Piotrowska, A., Borsuk, E., Hofman, B., Rava, V., Taverna, E., Włodarczyk, J., Pękowska, A. (2025). Primate Astrocyte Evolution Controls the Tempo of Neuronal Development. bioRxiv. 10.64898/2026.07.08.736743

Coffey, C. E., Wilkinson, W. E., Parashos, I. A., Soady, S. A., Sullivan, R. J., Patterson, L. J., Figiel, G. S., Webb, M. C., Spritzer, C. E., & Djang, W. T. (1992). Quantitative cerebral anatomy of the aging human brain: a cross-sectional study using magnetic resonance imaging. Neurology, 42(3 Pt 1), 527–536. 10.1212/wnl.42.3.527

Courtney, Y., & Lehtinen, M. K. (2025). Apocrine secretion by the choroid plexus. Fluids and barriers of the CNS, 22(1), 75. 10.1186/s12987-025-00684-3

Croft, D., Mundo, A. F., Haw, R., Milacic, M., Weiser, J., Wu, G., Caudy, M., Garapati, P., Gillespie, M., Kamdar, M. R., Jassal, B., Jupe, S., Matthews, L., May, B., Palatnik, S., Rothfels, K., Shamovsky, V., Song, H., Williams, M., Birney, E., … D’Eustachio, P. (2014). The Reactome pathway knowledgebase. Nucleic acids research, 42(Database issue), D472–D477. 10.1093/nar/gkt1102

Csárdi, G., & Nepusz, T. (2006). The Igraph Software Package for Complex Network Research. InterJournal, Complex Systems, 1695, 1–9. http://igraph.org

Desmond, M. E., Levitan, M. L., & Haas, A. R. (2005). Internal luminal pressure during early chick embryonic brain growth: descriptive and empirical observations. The anatomical record. Part A, Discoveries in molecular, cellular, and evolutionary biology, 285(2), 737–747. 10.1002/ar.a.20211

Dillies, M. A., Rau, A., Aubert, J., Hennequet-Antier, C., Jeanmougin, M., Servant, N., Keime, C., Marot, G., Castel, D., Estelle, J., Guernec, G., Jagla, B., Jouneau, L., Laloë, D., Le Gall, C., Schaëffer, B., Le Crom, S., Guedj, M., Jaffrézic, F., & French StatOmique Consortium (2013). A comprehensive evaluation of normalization methods for Illumina high-throughput RNA sequencing data analysis. Briefings in bioinformatics, 14(6), 671–683. 10.1093/bib/bbs046

Duperron, M. G., Knol, M. J., Le Grand, Q., Evans, T. E., Mishra, A., Tsuchida, A., Roshchupkin, G., Konuma, T., Trégouët, D. A., Romero, J. R., Frenzel, S., Luciano, M., Hofer, E., Bourgey, M., Dueker, N. D., Delgado, P., Hilal, S., Tankard, R. M., Dubost, F., Shin, J., … Debette, S. (2023). Genomics of perivascular space burden unravels early mechanisms of cerebral small vessel disease. Nature medicine, 29(4), 950–962. 10.1038/s41591-023-02268-w

Eichenbaum, H., Schoenbaum, G., Young, B., & Bunsey, M. (1996). Functional organization of the hippocampal memory system. Proceedings of the National Academy of Sciences of the United States of America, 93(24), 13500–13507. 10.1073/pnas.93.24.13500

Emmert, A. S., Iwasawa, E., Shula, C., Schultz, P., Lindquist, D., Dunn, R. S., Fugate, E. M., Hu, Y. C., Mangano, F. T., & Goto, J. (2019). Impaired neural differentiation and glymphatic CSF flow in the *Ccdc39* rat model of neonatal hydrocephalus: genetic interaction with *L1cam*. Disease models & mechanisms, 12(11), dmm040972. 10.1242/dmm.040972

Fadista, J., Manning, A. K., Florez, J. C., & Groop, L. (2016). The (in)famous GWAS P-value threshold revisited and updated for low-frequency variants. European journal of human genetics : EJHG, 24(8), 1202–1205. 10.1038/ejhg.2015.269

Fame, R. M., & Lehtinen, M. K. (2020). Emergence and Developmental Roles of the Cerebrospinal Fluid System. Developmental cell, 52(3), 261–275. 10.1016/j.devcel.2020.01.027

Ferrarini, L., Palm, W. M., Olofsen, H., van Buchem, M. A., Reiber, J. H., & Admiraal-Behloul, F. (2006). Shape differences of the brain ventricles in Alzheimer’s disease. NeuroImage, 32(3), 1060–1069. 10.1016/j.neuroimage.2006.05.048

Filippi, M., Cividini, C., Basaia, S., Spinelli, E. G., Castelnovo, V., Leocadi, M., Canu, E., & Agosta, F. (2023). Age-related vulnerability of the human brain connectome. Molecular psychiatry, 28(12), 5350–5358. 10.1038/s41380-023-02157-1

Fulcher, B. D., Little, M. A., & Jones, N. S. (2013). Highly comparative time-series analysis: the empirical structure of time series and their methods. Journal of the Royal Society, Interface, 10(83), 20130048. 10.1098/rsif.2013.0048

Gale, D. J., Vos de Wael, R., Benkarim, O., Bernhardt, B. (2021). Surfplot: publication-ready brain surface figures. https://zenodo.org/record/5567926

García-González, P., Puerta, R., Dehairs, J., Yang, C., Wang, C., Timsina, J., de Rojas, I., Olivé, C., Valenzuela, A., Bayón-Buján, P., Rovira, M., Montrreal, L., Capdevila, M., Muñoz-Morales, Á., Calm, B., Valero, S., Alegret, M., Marquié, M., The Global Neurodegeneration Proteomics Consortium (GNPC), Morris, J. C., Schindler, S. E., Holtzmann, D. M., Sanz, P., Tárraga, L., Khan, A., Sáez, M. E., Smets, B., Orellana, A., Montalbán, X., Boada, M., Cano, A., Liu, M., Ali, M. Cruchaga, C., Swinnen, J. V., Fernández, V., Cabrera-Socorro, A., Ruiz, A. (2026). CSF turnover reshapes biomarker interpretation in neurodegeneration studies. MedRxiv. 10.64898/2026.02.02.26345363

Gasquoine P. G. (2014). Contributions of the insula to cognition and emotion. Neuropsychology review, 24(2), 77–87. 10.1007/s11065-014-9246-9

Ge, Y. J., Wu, B. S., Zhang, Y., Chen, S. D., Zhang, Y. R., Kang, J. J., Deng, Y. T., Ou, Y. N., He, X. Y., Zhao, Y. L., Kuo, K., Ma, Q., Banaschewski, T., Barker, G. J., Bokde, A. L. W., Desrivières, S., Flor, H., Grigis, A., Garavan, H., Gowland, P., … Yu, J. T. (2024). Genetic architectures of cerebral ventricles and their overlap with neuropsychiatric traits. Nature human behaviour, 8(1), 164–180. 10.1038/s41562-023-01722-6

Gokhman, D., Nissim-Rafinia, M., Agranat-Tamir, L., Housman, G., García-Pérez, R., Lizano, E., Cheronet, O., Mallick, S., Nieves-Colón, M. A., Li, H., Alpaslan-Roodenberg, S., Novak, M., Gu, H., Osinski, J. M., Ferrando-Bernal, M., Gelabert, P., Lipende, I., Mjungu, D., Kondova, I., Bontrop, R., … Carmel, L. (2020). Differential DNA methylation of vocal and facial anatomy genes in modern humans. Nature communications, 11(1), 1189. 10.1038/s41467-020-15020-6

Gómez-Robles, A., Nicolaou, C., Smaers, J. B., & Sherwood, C. C. (2024). The evolution of human altriciality and brain development in comparative context. Nature ecology & evolution, 8(1), 133–146. 10.1038/s41559-023-02253-z

Gomolka, R. S., Hablitz, L. M., Mestre, H., Giannetto, M., Du, T., Hauglund, N. L., Xie, L., Peng, W., Martinez, P. M., Nedergaard, M., & Mori, Y. (2023). Loss of aquaporin-4 results in glymphatic system dysfunction via brain-wide interstitial fluid stagnation. eLife, 12, e82232. 10.7554/eLife.82232

Goovaerts, S., Naqvi, S., Hoskens, H., Herrick, N., Yuan, M., Shriver, M. D., Shaffer, J. R., Walsh, S., Weinberg, S. M., Wysocka, J., & Claes, P. (2025). Enhanced insights into the genetic architecture of 3D cranial vault shape using pleiotropy-informed GWAS. Communications biology, 8(1), 439. 10.1038/s42003-025-07875-6

Gouveia, R., Schaffer, L., Papp, S., Grammel, N., Kandzia, S., Head, S. R., Kleene, R., Schachner, M., Conradt, H. S., & Costa, J. (2012). Expression of glycogenes in differentiating human NT2N neurons. Downregulation of fucosyltransferase 9 leads to decreased Lewis(x) levels and impaired neurite outgrowth. Biochimica et biophysica acta, 1820(12), 2007–2019. 10.1016/j.bbagen.2012.09.004

Gregory, M. D., Kippenhan, J. S., Kohn, P., Eisenberg, D. P., Callicott, J. H., Kolachana, B., & Berman, K. F. (2021). Neanderthal-Derived Genetic Variation is Associated with Functional Connectivity in the Brains of Living Humans. Brain connectivity, 11(1), 38–44. 10.1089/brain.2020.0809

Gunz, P., Neubauer, S., Maureille, B., & Hublin, J. J. (2010). Brain development after birth differs between Neanderthals and modern humans. Current biology: CB, 20(21), R921–R922. 10.1016/j.cub.2010.10.018

Gunz, P., Tilot, A. K., Wittfeld, K., Teumer, A., Shapland, C. Y., van Erp, T. G. M., Dannemann, M., Vernot, B., Neubauer, S., Guadalupe, T., Fernández, G., Brunner, H. G., Enard, W., Fallon, J., Hosten, N., Völker, U., Profico, A., Di Vincenzo, F., Manzi, G., Kelso, J., … Fisher, S. E. (2019). Neandertal Introgression Sheds Light on Modern Human Endocranial Globularity. Current biology: CB, 29(1), 120–127.e5. 10.1016/j.cub.2018.10.065

Hablitz, L. M., & Nedergaard, M. (2021). The Glymphatic System: A Novel Component of Fundamental Neurobiology. The Journal of neuroscience : the official journal of the Society for Neuroscience, 41(37), 7698–7711. 10.1523/JNEUROSCI.0619-21.2021

Handrigan, G. R., Chitayat, D., Lionel, A. C., Pinsk, M., Vaags, A. K., Marshall, C. R., Dyack, S., Escobar, L. F., Fernandez, B. A., Stegman, J. C., Rosenfeld, J. A., Shaffer, L. G., Goodenberger, M., Hodge, J. C., Cain, J. E., Babul-Hirji, R., Stavropoulos, D. J., Yiu, V., Scherer, S. W., & Rosenblum, N. D. (2013). Deletions in 16q24.2 are associated with autism spectrum disorder, intellectual disability and congenital renal malformation. Journal of medical genetics, 50(3), 163–173. 10.1136/jmedgenet-2012-101288

Harrington, A. R., Kuzawa, C. W., & Boyer, D. M. (2019). Carotid foramen size in the human skull tracks developmental changes in cerebral blood flow and brain metabolism. American journal of physical anthropology, 169(1), 161–169. 10.1002/ajpa.23809

Hawrylycz, M. J., Lein, E. S., Guillozet-Bongaarts, A. L., Shen, E. H., Ng, L., Miller, J. A., van de Lagemaat, L. N., Smith, K. A., Ebbert, A., Riley, Z. L., Abajian, C., Beckmann, C. F., Bernard, A., Bertagnolli, D., Boe, A. F., Cartagena, P. M., Chakravarty, M. M., Chapin, M., Chong, J., Dalley, R. A., … Jones, A. R. (2012). An anatomically comprehensive atlas of the adult human brain transcriptome. Nature, 489(7416), 391–399. 10.1038/nature11405

Hawrylycz, M., Miller, J. A., Menon, V., Feng, D., Dolbeare, T., Guillozet-Bongaarts, A. L., Jegga, A. G., Aronow, B. J., Lee, C. K., Bernard, A., Glasser, M. F., Dierker, D. L., Menche, J., Szafer, A., Collman, F., Grange, P., Berman, K. A., Mihalas, S., Yao, Z., Stewart, L., … Lein, E. (2015). Canonical genetic signatures of the adult human brain. Nature neuroscience, 18(12), 1832–1844. 10.1038/nn.4171

Hemani, G., Tilling, K., & Davey Smith, G. (2017). Orienting the causal relationship between imprecisely measured traits using GWAS summary data. PLoS genetics, 13(11), e1007081. 10.1371/journal.pgen.1007081

Hemani, G., Zheng, J., Elsworth, B., Wade, K. H., Haberland, V., Baird, D., Laurin, C., Burgess, S., Bowden, J., Langdon, R., Tan, V. Y., Yarmolinsky, J., Shihab, H. A., Timpson, N. J., Evans, D. M., Relton, C., Martin, R. M., Davey Smith, G., Gaunt, T. R., & Haycock, P. C. (2018). The MR-Base platform supports systematic causal inference across the human phenome. eLife, 7, e34408. 10.7554/eLife.34408

Hill, J., Inder, T., Neil, J., Dierker, D., Harwell, J., & Van Essen, D. (2010). Similar patterns of cortical expansion during human development and evolution. Proceedings of the National Academy of Sciences of the United States of America, 107(29), 13135–13140. 10.1073/pnas.1001229107

Hoffmann, D. L., Standish, C. D., García-Diez, M., Pettitt, P. B., Milton, J. A., Zilhão, J., Alcolea-González, J. J., Cantalejo-Duarte, P., Collado, H., de Balbín, R., Lorblanchet, M., Ramos-Muñoz, J., Weniger, G. C., & Pike, A. W. G. (2018). U-Th dating of carbonate crusts reveals Neandertal origin of Iberian cave art. Science (New York, N.Y.), 359(6378), 912–915. 10.1126/science.aap7778

Hsu, J. L., Wei, Y. C., Toh, C. H., Hsiao, I. T., Lin, K. J., Yen, T. C., Liao, M. F., & Ro, L. S. (2023). Magnetic Resonance Images Implicate That Glymphatic Alterations Mediate Cognitive Dysfunction in Alzheimer Disease. Annals of neurology, 93(1), 164–174. 10.1002/ana.26516

Huang, S. Y., Ge, Y. J., Ren, P., Wu, B. S., Gong, W., Du, J., Chen, S. D., Kang, J. J., Ma, Q., Bokde, A. L. W., Desrivières, S., Garavan, H., Grigis, A., Lemaitre, H., Smolka, M. N., Hohmann, S., IMAGEN Consortium, Feng, J. F., Zhang, Y. R., Cheng, W., … Yu, J. T. (2025). Genome-wide association study unravels mechanisms of brain glymphatic activity. Nature communications, 16(1), 626. 10.1038/s41467-024-55706-9

Hublin, J. J., Neubauer, S., & Gunz, P. (2015). Brain ontogeny and life history in Pleistocene hominins. Philosophical transactions of the Royal Society of London. Series B, Biological sciences, 370(1663), 20140062. 10.1098/rstb.2014.0062

Jarrard L. E. (1993). On the role of the hippocampus in learning and memory in the rat. Behavioral and neural biology, 60(1), 9–26. 10.1016/0163-1047(93)90664-4

Jerison H. 2012. Evolution of the brain and intelligence. Elsevier

Jessen, N. A., Munk, A. S., Lundgaard, I., & Nedergaard, M. (2015). The Glymphatic System: A Beginner’s Guide. Neurochemical research, 40(12), 2583–2599. 10.1007/s11064-015-1581-6

Ju, X. C., Lee, S. Y., Ågren, R., Machado, L. C., Xing, J., Azama, C., Roy, M. C., Endo, T., Huttner, W., Siepel, A., Fukunaga, I., Zeberg, H., & Pääbo, S. (2025). The activity and expression of adenylosuccinate lyase were reduced during modern human evolution, affecting brain and behavior. Proceedings of the National Academy of Sciences of the United States of America, 122(32), e2508540122. 10.1073/pnas.2508540122

Kalinka, A. T. (2013). The probability of drawing intersections: extending the hypergeometric distribution. arXiv. https://arxiv.org/abs/1305.0717

Kameya, N., Sakai, I., Saito, K., Hamabe-Horiike, T., Shinmyo, Y., Nakada, M., Okuda, S., & Kawasaki, H. (2024). Evolutionary changes leading to efficient glymphatic circulation in the mammalian brain. Nature communications, 15(1), 10048. 10.1038/s41467-024-54372-1

Kim, N. J., Chowdhury, N. F., Buetow, K. H., Thompson, P. M., & Irimia, A. (2025). Genetic Insights into Brain Morphology: a Genome-Wide Association Study of Cortical Thickness and T1-Weighted MRI Gray Matter-White Matter Intensity Contrast. Neuroinformatics, 23(2), 26. 10.1007/s12021-025-09722-9

Kim, N. J., Mishra, A., Chowdhury, N. F., Anderson, S. D., Vega, O. M., Chaudhari, N. N., Buetow, K. H., Thompson, P. M., & Irimia, A. (2026). Deep neural networks and genome-wide associations reveal the polygenic architecture of local brain aging. GeroScience, 48(3), 3729–3753. 10.1007/s11357-025-02046-1

Kochiyama, T., Ogihara, N., Tanabe, H. C., Kondo, O., Amano, H., Hasegawa, K., Suzuki, H., Ponce de León, M. S., Zollikofer, C. P. E., Bastir, M., Stringer, C., Sadato, N., & Akazawa, T. (2018). Reconstructing the Neanderthal brain using computational anatomy. Scientific reports, 8(1), 6296. 10.1038/s41598-018-24331-0

Kochunov, P., Mangin, J. F., Coyle, T., Lancaster, J., Thompson, P., Rivière, D., Cointepas, Y., Régis, J., Schlosser, A., Royall, D. R., Zilles, K., Mazziotta, J., Toga, A., & Fox, P. T. (2005). Age-related morphology trends of cortical sulci. Human brain mapping, 26(3), 210–220. 10.1002/hbm.20198

Kolberg, L., Raudvere, U., Kuzmin, I., Adler, P., Vilo, J., & Peterson, H. (2023). g:Profiler-interoperable web service for functional enrichment analysis and gene identifier mapping (2023 update). Nucleic acids research, 51(W1), W207–W212. 10.1093/nar/gkad347

Krohn, S., Romanello, A., von Schwanenflug, N., Rasmussen, J. M., Buss, C., Valk, S. L., Madan, C. R., & Finke, C. (2026). Fractal analysis of brain shape formation predicts age and genetic similarity in human newborns. Nature neuroscience, 29(1), 171–185. 10.1038/s41593-025-02107-w

Kuhlwilm, M., & Boeckx, C. (2019). A catalog of single nucleotide changes distinguishing modern humans from archaic hominins. Scientific reports, 9(1), 8463. 10.1038/s41598-019-44877-x

Kuzawa, C. W., Chugani, H. T., Grossman, L. I., Lipovich, L., Muzik, O., Hof, P. R., Wildman, D. E., Sherwood, C. C., Leonard, W. R., & Lange, N. (2014). Metabolic costs and evolutionary implications of human brain development. Proceedings of the National Academy of Sciences of the United States of America, 111(36), 13010–13015. 10.1073/pnas.1323099111

Kwon, Y. H., Jang, S. H., & Yeo, S. S. (2014). Age-related changes of lateral ventricular width and periventricular white matter in the human brain: a diffusion tensor imaging study. Neural regeneration research, 9(9), 986–989. 10.4103/1673-5374.133152

Lasagna-Reeves, C. A., de Haro, M., Hao, S., Park, J., Rousseaux, M. W., Al-Ramahi, I., Jafar-Nejad, P., Vilanova-Velez, L., See, L., De Maio, A., Nitschke, L., Wu, Z., Troncoso, J. C., Westbrook, T. F., Tang, J., Botas, J., & Zoghbi, H. Y. (2016). Reduction of Nuak1 Decreases Tau and Reverses Phenotypes in a Tauopathy Mouse Model. Neuron, 92(2), 407–418. 10.1016/j.neuron.2016.09.022

Lemaitre, H., Le Guen, Y., Tilot, A. K., Stein, J. L., Philippe, C., Mangin, J. F., Fisher, S. E., & Frouin, V. (2023). Genetic variations within human gained enhancer elements affect human brain sulcal morphology. NeuroImage, 265, 119773. 10.1016/j.neuroimage.2022.119773

Leonardi, O., Lewerissa, E., Aprile, D., Barakat, T. S., Culotta, L., Deng, R., Haazen, L., Linda, K., Majoie, M., Mordelt, A., Ortale, A., Oudakker, A., Prazzoli, F., Puvogel, S., Scheefals, N., Schelhaas, H. J., Schoenmaker, C., van Hughte, E., van Bokhoven, H., …, Nadif Kasri, N. (2025). Human-Specific Regulation of *CHD2* Controls the Tempo of Cortical Development through Lysosomal Homeostasis. Biorxiv. 10.1101/2025.01.21.634145

Li, Z., Xu, J., & Li, Y. (2026). The dual role of the pentose phosphate pathway in homeostasis and regeneration. Cell metabolism, 38(3), 449–451. 10.1016/j.cmet.2026.02.002

Li, M., Santpere, G., Imamura Kawasawa, Y., Evgrafov, O. V., Gulden, F. O., Pochareddy, S., Sunkin, S. M., Li, Z., Shin, Y., Zhu, Y., Sousa, A. M. M., Werling, D. M., Kitchen, R. R., Kang, H. J., Pletikos, M., Choi, J., Muchnik, S., Xu, X., Wang, D., Lorente-Galdos, B., … Sestan, N. (2018). Integrative functional genomic analysis of human brain development and neuropsychiatric risks. Science (New York, N.Y.), 362(6420), eaat7615. 10.1126/science.aat7615

Li, Y., Zhou, Y., Zhong, W., Zhu, X., Chen, Y., Zhang, K., He, Y., Luo, Z., Ran, W., Sun, J., & Lou, M. (2023). Choroid Plexus Enlargement Exacerbates White Matter Hyperintensity Growth through Glymphatic Impairment. Annals of neurology, 94(1), 182–195. 10.1002/ana.26648

Libedinsky, I., Wei, Y., de Leeuw, C., Rilling, J. K., Posthuma, D., & van den Heuvel, M. P. (2025). The emergence of genetic variants linked to brain and cognitive traits in human evolution. Cerebral cortex (New York, N.Y. : 1991), 35(8), bhaf127. 10.1093/cercor/bhaf127

Lin, S., Guo, M., Liang, Q., Lin, X., Chen, S., Li, Y., Chen, P., & Qiu, Y. (2024). Evaluation of Glymphatic System Development in Neonatal Brain via Diffusion Analysis along the Perivascular Space Index. Annals of neurology, 96(5), 970–980. 10.1002/ana.27047

Liu, X., Blazejewski, S. M., Bennison, S. A., & Toyo-Oka, K. (2021). Glutathione S-transferase Pi (Gstp) proteins regulate neuritogenesis in the developing cerebral cortex. Human molecular genetics, 30(1), 30–45. 10.1093/hmg/ddab003

Lombard, M. (2025a). From Complex Techno-behaviour to Complex Attention Through the Genes of the Precuneus. Journal of Archaeological Method and Theory, 32(46). 10.1007/s10816-025-09716-6

Lombard, M. (2025b). Towards an archaeology of attention: a neuro-genetic exploration. Journal of Archaeological Science, 182, 106344. 10.1016/j.jas.2025.106344

Lowery, L. A., & Sive, H. (2009). Totally tubular: the mystery behind function and origin of the brain ventricular system. BioEssays : news and reviews in molecular, cellular and developmental biology, 31(4), 446–458. 10.1002/bies.200800207

Mantini, D., Corbetta, M., Romani, G. L., Orban, G. A., & Vanduffel, W. (2013). Evolutionarily novel functional networks in the human brain?. The Journal of neuroscience: the official journal of the Society for Neuroscience, 33(8), 3259–3275. 10.1523/JNEUROSCI.4392-12.2013

Margulies, D. S., Ghosh, S. S., Goulas, A., Falkiewicz, M., Huntenburg, J. M., Langs, G., Bezgin, G., Eickhoff, S. B., Castellanos, F. X., Petrides, M., Jefferies, E., & Smallwood, J. (2016). Situating the default-mode network along a principal gradient of macroscale cortical organization. Proceedings of the National Academy of Sciences of the United States of America, 113(44), 12574–12579. 10.1073/pnas.1608282113

Markello, R. D., Arnatkeviciute, A., Poline, J. B., Fulcher, B. D., Fornito, A., & Misic, B. (2021). Standardizing workflows in imaging transcriptomics with the abagen toolbox. eLife, 10, e72129. 10.7554/eLife.72129

Markello, R. D., Hansen, J. Y., Liu, Z. Q., Bazinet, V., Shafiei, G., Suárez, L. E., Blostein, N., Seidlitz, J., Baillet, S., Satterthwaite, T. D., Chakravarty, M. M., Raznahan, A., & Misic, B. (2022). neuromaps: structural and functional interpretation of brain maps. Nature methods, 19(11), 1472–1479. 10.1038/s41592-022-01625-w

McCoy, R. C., Wakefield, J., & Akey, J. M. (2017). Impacts of Neanderthal-Introgressed Sequences on the Landscape of Human Gene Expression. Cell, 168(5), 916–927.e12. 10.1016/j.cell.2017.01.038

Mekki, Y. N. (2022). The genetic architecture of the human language connectome. [Doctoral dissertation, Université Paris-Saclay]. HAL theses. https://theses.hal.science/tel-03649334v1

Mendes, M., Engchuan, W., Thompson, S., Zhou, X., Safarian, N., Zeya, D., Trost, B., Bautista Salazar, N., Ma, C., Thiruvahindrapuram, B., Vorstman, J., Scherer, S., Breetvelt, E. Genome-Wide Significance Reconsidered: Low-Frequency Variants and Regulatory Networks in Autism. MedRXiv. (2026). 10.64898/2026.02.11.26346090

Mestre, H., Hablitz, L. M., Xavier, A. L., Feng, W., Zou, W., Pu, T., Monai, H., Murlidharan, G., Castellanos Rivera, R. M., Simon, M. J., Pike, M. M., Plá, V., Du, T., Kress, B. T., Wang, X., Plog, B. A., Thrane, A. S., Lundgaard, I., Abe, Y., Yasui, M., … Nedergaard, M. (2018). Aquaporin-4-dependent glymphatic solute transport in the rodent brain. eLife, 7, e40070. 10.7554/eLife.40070

Mil, J., Soto, J. A., Matulionis, N., Krall, A., Day, F., Stiles, L., Montales, K. P., Azizad, D. J., Gonzalez, C. E., Nano, P. R., Antoni, A. M., Perez-Ramirez, C., Nguyen, C. V., Kan, R. L., Andrews, M. G., Christofk, H. R., Bhaduri, A. (2025). Metabolic Atlas of Early Human Cortex Identifies Regulators of Cell Fate Transitions. bioRxiv. 10.1101/2025.03.10.642470

Miller, J. A., Ding, S. L., Sunkin, S. M., Smith, K. A., Ng, L., Szafer, A., Ebbert, A., Riley, Z. L., Royall, J. J., Aiona, K., Arnold, J. M., Bennet, C., Bertagnolli, D., Brouner, K., Butler, S., Caldejon, S., Carey, A., Cuhaciyan, C., Dalley, R. A., Dee, N., … Lein, E. S. (2014). Transcriptional landscape of the prenatal human brain. Nature, 508(7495), 199–206. 10.1038/nature13185

Moberget, T., van der Meer, D., Bahrami, S., Roelfs, D., Frei, O., Kaufmann, T., Fernandez-Cabello, S., Kim, M., Wolfers, T., Diedrichsen, J., Smeland, O. B., Shadrin, A., Dale, A., Andreassen, O. A., & Westlye, L. T. (2026). The genetic architecture of human cerebellar morphology supports a key role for the cerebellum in human evolution and psychopathology. Communications biology, 9(1), 445. 10.1038/s42003-026-09664-1

Molz, B., Alberro, M. L., Eising, E., Schijven, D., Alagöz, G., Francks, C., & Fisher, S. E. (2025). Evaluating the effects of archaic protein-altering variants in living human adults. Science advances, 11(50), eads5703. 10.1126/sciadv.ads5703

Molz, B., Eising, E., Alagöz, G., Schijven, D., Francks, C., Gunz, P., Fisher, S. (2024). Imaging genomics reveals genetic architecture of the globular human braincase. bioRxiv. 10.1101/2024.03.20.585712

Moriano, J., Boeckx, C., & Kuhlwilm, M. (2025). Palaeogenomics: A window into the genetic basis of derived traits in *Homo sapiens*, in L. Raviv & C. Boeckx (eds.), The Oxford Handbook of Approaches to Language Evolution. Oxford Handbooks.

Namkung, H., Kim, S. H., & Sawa, A. (2017). The Insula: An Underestimated Brain Area in Clinical Neuroscience, Psychiatry, and Neurology. Trends in neurosciences, 40(4), 200–207. 10.1016/j.tins.2017.02.002

Nestor, S. M., Rupsingh, R., Borrie, M., Smith, M., Accomazzi, V., Wells, J. L., Fogarty, J., Bartha, R., & Alzheimer’s Disease Neuroimaging Initiative (2008). Ventricular enlargement as a possible measure of Alzheimer’s disease progression validated using the Alzheimer’s disease neuroimaging initiative database. Brain : a journal of neurology, 131(Pt 9), 2443–2454. 10.1093/brain/awn146

Neubauer, S., Hublin, J. J., & Gunz, P. (2018). The evolution of modern human brain shape. Science advances, 4(1), eaao5961. 10.1126/sciadv.aao5961

Noel, M., Zürcher, Y. M., Tulin, E. K. C., & Cummings, R. D. (2025). The Importance of N- and O-Glycosylation of Brain Cell Surface Glycoproteins. Glycobiology, 35(11), cwaf054. 10.1093/glycob/cwaf054

Ou, Y. N., Ge, Y. J., Wu, B. S., Zhang, Y., Jiang, Y. C., Kuo, K., Yang, L., Tan, L., Feng, J. F., Cheng, W., & Yu, J. T. (2023). The genetic architecture of fornix white matter microstructure and their involvement in neuropsychiatric disorders. Translational psychiatry, 13(1), 180. 10.1038/s41398-023-02475-6

Pandya, V. A., & Patani, R. (2021). Region-specific vulnerability in neurodegeneration: lessons from normal ageing. Ageing research reviews, 67, 101311. 10.1016/j.arr.2021.101311

Parada, C., Gato, A., Aparicio, M., & Bueno, D. (2006). Proteome analysis of chick embryonic cerebrospinal fluid. Proteomics, 6(1), 312–320. 10.1002/pmic.200500085

Parikshak, N. N., Luo, R., Zhang, A., Won, H., Lowe, J. K., Chandran, V., Horvath, S., & Geschwind, D. H. (2013). Integrative functional genomic analyses implicate specific molecular pathways and circuits in autism. Cell, 155(5), 1008–1021. 10.1016/j.cell.2013.10.031

Parkinson, W., Dear, M. L., Rushton, E., & Broadie, K. (2013). N-glycosylation requirements in neuromuscular synaptogenesis. Development (Cambridge, England), 140(24), 4970–4981. 10.1242/dev.099192

Phelps E. A. (2006). Emotion and cognition: insights from studies of the human amygdala. Annual review of psychology, 57, 27–53. 10.1146/annurev.psych.56.091103.070234

Peng, T., Lin, Y., Xu, X., Li, J., Liu, M., Zhang, C., Liao, X., Ji, X., Xiong, Z., Gu, Z., Cai, X., Tao, T., Zhang, Y., Zhu, L., Zhuang, D., Huang, X., Xiong, M., Zhang, P., Liu, J., & Cheng, G. (2025). Assessing neonatal brain glymphatic system development using diffusion tensor imaging along the perivascular space and choroid plexus volume. BMC medical imaging, 25(1), 126. 10.1186/s12880-025-01673-6

Pereira-Pedro, A. S., Bruner, E., Gunz, P., & Neubauer, S. (2020). A morphometric comparison of the parietal lobe in modern humans and Neanderthals. Journal of human evolution, 142, 102770. 10.1016/j.jhevol.2020.102770

Perrot, M., Rivière, D., & Mangin, J. F. (2011). Cortical sulci recognition and spatial normalization. Medical image analysis, 15(4), 529–550. 10.1016/j.media.2011.02.008

Petanjek, Z., Judaš, M., Šimic, G., Rasin, M. R., Uylings, H. B., Rakic, P., & Kostovic, I. (2011). Extraordinary neoteny of synaptic spines in the human prefrontal cortex. Proceedings of the National Academy of Sciences of the United States of America, 108(32), 13281–13286. 10.1073/pnas.1105108108

Pinson, A., Xing, L., Namba, T., Kalebic, N., Peters, J., Oegema, C. E., Traikov, S., Reppe, K., Riesenberg, S., Maricic, T., Derihaci, R., Wimberger, P., Pääbo, S., & Huttner, W. B. (2022). Human TKTL1 implies greater neurogenesis in frontal neocortex of modern humans than Neanderthals. Science (New York, N.Y.), 377(6611), eabl6422. 10.1126/science.abl6422

Ploner, A., & Frei, O. (2026). cfdr.pleio: Conditional and Conjunctional FDR for GWAS. R package version 0.0.0.9100.

Posit team. (2025). RStudio: Integrated Development Environment for R [Computer software]. Boston, MA. https://posit.co/

Pourmajidian, M., Hansen, J. Y., Shafiei, G., Misic, B., & Dagher, A. (2026). Five energy metabolism pathways show distinct regional distributions and lifespan trajectories in the human brain. PLoS biology, 24(1), e3003619. 10.1371/journal.pbio.3003619

Primus, S. A., Hoffstaedter, F., Raimondo, F., Eickhoff, S. B., Winkelmann, J., Oexle, K., & Patil, K. R. (2025). Beyond volume: Unraveling the genetics of human brain geometry. Science advances, 11(24), eadr1644. 10.1126/sciadv.adr1644

Quackenbush J. (2002). Microarray data normalization and transformation. Nature genetics, 32 *Suppl*, 496–501. 10.1038/ng1032

Ran, L., Fang, Y., Cheng, C., He, Y., Shao, Z., Kong, Y., Huang, H., Xu, S., Luo, X., Wang, W., Hao, X., & Wang, M. (2025). Genome-wide and phenome-wide studies provided insights into brain glymphatic system function and its clinical associations. Science advances, 11(3), eadr4606. 10.1126/sciadv.adr4606

Räsänen, J., Heikkinen, S., Mäklin, K., Lipponen, A., Kuulasmaa, T., Mehtonen, J., Korhonen, V. E., Junkkari, A., Grenier-Boley, B., Bellenguez, C., Oinas, M., Avellan, C., Frantzén, J., Kotkansalo, A., Rinne, J., Ronkainen, A., Kauppinen, M., von Und Zu Fraunberg, M., Lönnrot, K., Satopää, J., … Leinonen, V. (2024). Risk Variants Associated With Normal Pressure Hydrocephalus: Genome-Wide Association Study in the FinnGen Cohort. Neurology, 103(5), e209694. 10.1212/WNL.0000000000209694

Rava, V., Restelli, E., Mirabella, F., Silvestrini, L., Cannone, E., Ciuba, K., Abbas, M., Graziadei, A., Francolini, M., Pękowska, A., Taverna, E. (2026). Evolutionary diversification of lipid logistics shapes synaptic maturation in primates. bioRxiv. 10.64898/2026.07.08.734953

Reardon, P. K., Seidlitz, J., Vandekar, S., Liu, S., Patel, R., Park, M. T. M., Alexander-Bloch, A., Clasen, L. S., Blumenthal, J. D., Lalonde, F. M., Giedd, J. N., Gur, R. C., Gur, R. E., Lerch, J. P., Chakravarty, M. M., Satterthwaite, T. D., Shinohara, R. T., & Raznahan, A. (2018). Normative brain size variation and brain shape diversity in humans. Science (New York, N.Y.), 360(6394), 1222–1227. 10.1126/science.aar2578

Roebroeks, W., & Soressi, M. (2016). Neandertals revised. Proceedings of the National Academy of Sciences of the United States of America, 113(23), 6372–6379. 10.1073/pnas.1521269113

Roll M. (2024). Heschl’s gyrus and the temporal pole: The cortical lateralization of language. NeuroImage, 303, 120930. 10.1016/j.neuroimage.2024.120930

Romero-Garcia, R., Whitaker, K. J., Váša, F., Seidlitz, J., Shinn, M., Fonagy, P., Dolan, R. J., Jones, P. B., Goodyer, I. M., NSPN Consortium, Bullmore, E. T., & Vértes, P. E. (2018). Structural covariance networks are coupled to expression of genes enriched in supragranular layers of the human cortex. NeuroImage, 171, 256–267. 10.1016/j.neuroimage.2017.12.060

Santacroce, F., Cachia, A., Fragueiro, A., Grande, E., Roell, M., Baldassarre, A., Sestieri, C., & Committeri, G. (2024). Human intraparietal sulcal morphology relates to individual differences in language and memory performance. Communications biology, 7(1), 520. 10.1038/s42003-024-06175-9

Satterthwaite, T. D., Shinohara, R. T., Wolf, D. H., Hopson, R. D., Elliott, M. A., Vandekar, S. N., Ruparel, K., Calkins, M. E., Roalf, D. R., Gennatas, E. D., Jackson, C., Erus, G., Prabhakaran, K., Davatzikos, C., Detre, J. A., Hakonarson, H., Gur, R. C., & Gur, R. E. (2014). Impact of puberty on the evolution of cerebral perfusion during adolescence. Proceedings of the National Academy of Sciences of the United States of America, 111(23), 8643–8648. 10.1073/pnas.1400178111

Scerri, E. M. L., Thomas, M. G., Manica, A., Gunz, P., Stock, J. T., Stringer, C., Grove, M., Groucutt, H. S., Timmermann, A., Rightmire, G. P., d’Errico, F., Tryon, C. A., Drake, N. A., Brooks, A. S., Dennell, R. W., Durbin, R., Henn, B. M., Lee-Thorp, J., deMenocal, P., Petraglia, M. D., … Chikhi, L. (2018). Did Our Species Evolve in Subdivided Populations across Africa, and Why Does It Matter?. Trends in ecology & evolution, 33(8), 582–594. 10.1016/j.tree.2018.05.005

Scerri, E. M. L., & Will, M. (2023). The revolution that still isn’t: The origins of behavioral complexity in Homo sapiens. Journal of human evolution, 179, 103358. 10.1016/j.jhevol.2023.103358

Schaefer, A., Kong, R., Gordon, E. M., Laumann, T. O., Zuo, X. N., Holmes, A. J., Eickhoff, S. B., & Yeo, B. T. T. (2018). Local-Global Parcellation of the Human Cerebral Cortex from Intrinsic Functional Connectivity MRI. Cerebral cortex (New York, N.Y. : 1991), 28(9), 3095–3114. 10.1093/cercor/bhx179

Seymour, R. S., Bosiocic, V., & Snelling, E. P. (2016). Fossil skulls reveal that blood flow rate to the brain increased faster than brain volume during human evolution. Royal Society open science, 3(8), 160305. 10.1098/rsos.160305

Sha, Z., Schijven, D., Fisher, S. E., & Francks, C. (2023). Genetic architecture of the white matter connectome of the human brain. Science advances, 9(7), eadd2870. 10.1126/sciadv.add2870

Sha, Z., Warrier, V., Bethlehem, R. A. I., Schultz, L. M., Merikangas, A., Sun, K. Y., Gur, R. C., Gur, R. E., Shinohara, R. T., Gandal, M. J., Seidlitz, J., Almasy, L., Andreassen, O. A., & Alexander-Bloch, A. F. (2025). The overlapping genetic architecture of psychiatric disorders and cortical brain structure. Nature. Mental health, 3(9), 1020–1036. 10.1038/s44220-025-00475-7

Shan, Y., Qiao, H., He, Y., Chu, L., Zeng, D., Dong, X., Zhao, T., Liao, X., Chen, X., Xia, Y., Lei, T., Sun, L., Men, W., Chen, R., Ma, L., Ren, X., Wang, Y., Wang, D., Hu, M., Pan, Z., … Li, S. (2026). Linking changes in sulcal morphometry to cognitive development from childhood to adolescence. Communications biology, 9(1), 767. 10.1038/s42003-026-09956-6

Shen M. D. (2018). Cerebrospinal fluid and the early brain development of autism. Journal of neurodevelopmental disorders, 10(1), 39. 10.1186/s11689-018-9256-7

Shine, J. M., Lewis, L. D., Garrett, D. D., & Hwang, K. (2023). The impact of the human thalamus on brain-wide information processing. Nature reviews. Neuroscience, 24(7), 416–430. 10.1038/s41583-023-00701-0

Singh, A., Kaakinen, M., Elamaa, H., Kiviniemi, V., & Eklund, L. (2025). The glycosaminoglycan chains of perlecan regulate the perivascular fluid transport. Fluids and barriers of the CNS, 22(1), 48. 10.1186/s12987-025-00648-7

Smedley, D., Haider, S., Ballester, B., Holland, R., London, D., Thorisson, G., & Kasprzyk, A. (2009). BioMart--biological queries made easy. BMC genomics, 10, 22. 10.1186/1471-2164-10-22

Snyder, W. E., Vértes, P. E., Kyriakopoulou, V., Wagstyl, K., Williams, L. Z. J., Moraczewski, D., Thomas, A. G., Karolis, V. R., Seidlitz, J., Rivière, D., Robinson, E. C., Mangin, J. F., Raznahan, A., & Bullmore, E. T. (2024). A bimodal taxonomy of adult human brain sulcal morphology related to timing of fetal sulcation and trans-sulcal gene expression gradients. Neuron, 112(20), 3396–3411.e6. 10.1016/j.neuron.2024.07.023

Song, Y., Luengo-Gutierrez, L., Sagi-Kiss, V., Kong, G., Huang, H., Steinruecke, M., Zhou, L., Yuan, Z., De Virgiliis, F., Pap, I., Decourt, C., Yan, Y., Park, H. H., Zhang, H., Wei, J., Want, E., Tong, X., Takats, Z., & Di Giovanni, S. (2026). A glycolytic shunt via the pentose phosphate pathway is a metabolic checkpoint for nervous system sensory homeostasis and axonal regeneration. Cell, 189(4), 1211–1227.e25. 10.1016/j.cell.2025.12.003

Sowell, E. R., Thompson, P. M., Leonard, C. M., Welcome, S. E., Kan, E., & Toga, A. W. (2004). Longitudinal mapping of cortical thickness and brain growth in normal children. The Journal of neuroscience: the official journal of the Society for Neuroscience, 24(38), 8223–8231. 10.1523/JNEUROSCI.1798-04.2004

Spence, J. P., Mostafavi, H., Ota, M., Milind, N., Gjorgjieva, T., Smith, C. J., Simons, Y. B., Sella, G., & Pritchard, J. K. (2026). Specificity, length and luck drive gene rankings in association studies. Nature, 649(8098), 918–925. 10.1038/s41586-025-09703-7

Steiner, S., Foster, K., Chinn, R., Pratt, J., Fernandes, S., Sharma, A., Santos, R., Metallo, C., Marchetto, M. C., Gage, F. (2026). A Role for Astrocyte Metabolism in Species-Specific Neuronal Development. bioRxiv. 10.64898/2026.07.15.737608

Stincone, A., Prigione, A., Cramer, T., Wamelink, M. M., Campbell, K., Cheung, E., Olin-Sandoval, V., Grüning, N. M., Krüger, A., Tauqeer Alam, M., Keller, M. A., Breitenbach, M., Brindle, K. M., Rabinowitz, J. D., & Ralser, M. (2015). The return of metabolism: biochemistry and physiology of the pentose phosphate pathway. Biological reviews of the Cambridge Philosophical Society, 90(3), 927–963. 10.1111/brv.12140

Stringer C. (2016). The origin and evolution of Homo sapiens. Philosophical transactions of the Royal Society of London. Series B, Biological sciences, 371(1698), 20150237. 10.1098/rstb.2015.0237

Stringer C. (2022). The development of ideas about a recent African origin for Homo sapiens. Journal of anthropological sciences = Rivista di antropologia: JASS, 100, 5–18. 10.4436/JASS.10009

Sun, B. B., Loomis, S. J., Pizzagalli, F., Shatokhina, N., Painter, J. N., Foley, C. N., Biogen Biobank Team, Jensen, M. E., McLaren, D. G., Chintapalli, S. S., Zhu, A. H., Dixon, D., Islam, T., Ba Gari, I., Runz, H., Medland, S. E., Thompson, P. M., Jahanshad, N., & Whelan, C. D. (2022). Genetic map of regional sulcal morphology in the human brain from UK biobank data. Nature communications, 13(1), 6071. 10.1038/s41467-022-33829-1

Sun, H., Jiang, R., Dai, W., Dufford, A. J., Noble, S., Spann, M. N., Gu, S., & Scheinost, D. (2023). Network controllability of structural connectomes in the neonatal brain. Nature communications, 14(1), 5820. 10.1038/s41467-023-41499-w

Sun, H., Vernetti, A., Spann, M., Chawarska, K., Ment, L., & Scheinost, D. (2025). White-matter controllability at birth predicts social engagement and language outcomes in toddlerhood. bioRxiv. 10.1101/2025.02.19.638284

Sun, H., Toyonaga, T., Mehta, S., Liu, H., Camp, C., Rossano, S., Fowles, K., Martins, S., Spann, M., Groman, S., Carson, R. E., & Scheinost, D. (2026). Fetal network controllability co-develops with synaptic development and synchronizes with maternal network controllability during pregnancy. bioRxiv. 10.1101/2025.09.03.674103

Suzuki, R., & Shimodaira, H. (2006). Pvclust: an R package for assessing the uncertainty in hierarchical clustering. Bioinformatics (Oxford, England), 22(12), 1540–1542. 10.1093/bioinformatics/btl117

Sydnor, V. J., Larsen, B., Bassett, D. S., Alexander-Bloch, A., Fair, D. A., Liston, C., Mackey, A. P., Milham, M. P., Pines, A., Roalf, D. R., Seidlitz, J., Xu, T., Raznahan, A., & Satterthwaite, T. D. (2021). Neurodevelopment of the association cortices: Patterns, mechanisms, and implications for psychopathology. Neuron, 109(18), 2820–2846. 10.1016/j.neuron.2021.06.016

Sykes, R. W. (2020). Kindred: 300,000 Years of Neanderthal Life and Afterlife. Bloomsbury Publishing.

Takagi, K., & Kondo, O. (2026). Postnatal interaction of size and shape in the human endocranium and brain structures. Journal of anatomy, 248(3), 395–411. 10.1111/joa.70040

Tanabe, M., Saito, Y., Takasaki, A., Nakano, K., Yamamoto, S., Suzuki, C., Kawamura, N., Hattori, A., Oikawa, M., Nagashima, S., Yanagi, S., Yamaguchi, T., & Fukuda, T. (2025). Role of immature choroid plexus in the pathology of model mice and human iPSC-derived organoids with autism spectrum disorder. Cell reports, 44(1), 115133. 10.1016/j.celrep.2024.115133

Tung, S. S., Yan, X., Fascendini, B., Tyagi, C., Reyes, C. M., Ducre, K., Perez, K., Allen, A., Horenziak, J., Wu, H., Keil, B., Natu, V. S., & Grill-Spector, K. (2026). How do infant brains fold? Sulcal deepening is linked to development of sulcal span, thickness, curvature, and microstructure. Communications biology, 9(1), 258. 10.1038/s42003-026-09536-8

Tzourio-Mazoyer, N., Landeau, B., Papathanassiou, D., Crivello, F., Etard, O., Delcroix, N., Mazoyer, B., & Joliot, M. (2002). Automated anatomical labeling of activations in SPM using a macroscopic anatomical parcellation of the MNI MRI single-subject brain. NeuroImage, 15(1), 273–289. 10.1006/nimg.2001.0978

Van Essen, D. C., Glasser, M. F., Dierker, D. L., Harwell, J., & Coalson, T. (2012). Parcellations and hemispheric asymmetries of human cerebral cortex analyzed on surface-based atlases. Cerebral cortex (New York, N.Y. : 1991), 22(10), 2241–2262. 10.1093/cercor/bhr291

Váša, F., Seidlitz, J., Romero-Garcia, R., Whitaker, K. J., Rosenthal, G., Vértes, P. E., Shinn, M., Alexander-Bloch, A., Fonagy, P., Dolan, R. J., Jones, P. B., Goodyer, I. M., NSPN consortium, Sporns, O., & Bullmore, E. T. (2018). Adolescent Tuning of Association Cortex in Human Structural Brain Networks. Cerebral cortex (New York, N.Y. : 1991), 28(1), 281–294. 10.1093/cercor/bhx249

Vickery, S., Patil, K. R., Dahnke, R., Hopkins, W. D., Sherwood, C. C., Caspers, S., Eickhoff, S. B., & Hoffstaedter, F. (2024). The uniqueness of human vulnerability to brain aging in great ape evolution. Science advances, 10(35), eado2733. 10.1126/sciadv.ado2733

Vos de Wael, R., Benkarim, O., Paquola, C., Lariviere, S., Royer, J., Tavakol, S., Xu, T., Hong, S. J., Langs, G., Valk, S., Misic, B., Milham, M., Margulies, D., Smallwood, J., & Bernhardt, B. C. (2020). BrainSpace: a toolbox for the analysis of macroscale gradients in neuroimaging and connectomics datasets. Communications biology, 3(1), 103. 10.1038/s42003-020-0794-7

Wagstyl, K., Adler, S., Seidlitz, J., Vandekar, S., Mallard, T. T., Dear, R., DeCasien, A. R., Satterthwaite, T. D., Liu, S., Vértes, P. E., Shinohara, R. T., Alexander-Bloch, A., Geschwind, D. H., & Raznahan, A. (2024). Transcriptional cartography integrates multiscale biology of the human cortex. eLife, 12, RP86933. 10.7554/eLife.86933

Wang, C., Yang, C., Western, D., Ali, M., Wang, Y., Phuah, C. L., Budde, J., Wang, L., Gorijala, P., Timsina, J., Ruiz, A., Pastor, P., Fernandez, M. V., Dominantly Inherited Alzheimer Network (DIAN), Alzheimer’s Disease Neuroimaging Initiative (ADNI), Panyard, D. J., Engelman, C. D., Deming, Y., Boada, M., Cano, A., … Cruchaga, C. (2024). Genetic architecture of cerebrospinal fluid and brain metabolite levels and the genetic colocalization of metabolites with human traits. Nature genetics, 56(12), 2685–2695. 10.1038/s41588-024-01973-7

Watanabe, K., Stringer, S., Frei, O., Umićević Mirkov, M., de Leeuw, C., Polderman, T. J. C., van der Sluis, S., Andreassen, O. A., Neale, B. M., & Posthuma, D. (2019). A global overview of pleiotropy and genetic architecture in complex traits. Nature genetics, 51(9), 1339–1348. 10.1038/s41588-019-0481-0

Wei, Y., de Lange, S. C., Scholtens, L. H., Watanabe, K., Ardesch, D. J., Jansen, P. R., Savage, J. E., Li, L., Preuss, T. M., Rilling, J. K., Posthuma, D., & van den Heuvel, M. P. (2019). Genetic mapping and evolutionary analysis of human-expanded cognitive networks. Nature communications, 10(1), 4839. 10.1038/s41467-019-12764-8

Weiss, C. V., Harshman, L., Inoue, F., Fraser, H. B., Petrov, D. A., Ahituv, N., & Gokhman, D. (2021). The *cis*-regulatory effects of modern human-specific variants. eLife, 10, e63713. 10.7554/eLife.63713

Western, D., Timsina, J., Wang, L., Wang, C., Yang, C., Phillips, B., Wang, Y., Liu, M., Ali, M., Beric, A., Gorijala, P., Kohlfeld, P., Budde, J., Levey, A. I., Morris, J. C., Perrin, R. J., Ruiz, A., Marquié, M., Boada, M., de Rojas, I., … Cruchaga, C. (2024). Proteogenomic analysis of human cerebrospinal fluid identifies neurologically relevant regulation and implicates causal proteins for Alzheimer’s disease. Nature genetics, 56(12), 2672–2684. 10.1038/s41588-024-01972-8

Wightman, D. P., Jansen, I. E., Savage, J. E., Shadrin, A. A., Bahrami, S., Holland, D., Rongve, A., Børte, S., Winsvold, B. S., Drange, O. K., Martinsen, A. E., Skogholt, A. H., Willer, C., Bråthen, G., Bosnes, I., Nielsen, J. B., Fritsche, L. G., Thomas, L. F., Pedersen, L. M., Gabrielsen, M. E., … Posthuma, D. (2021). A genome-wide association study with 1,126,563 individuals identifies new risk loci for Alzheimer’s disease. Nature genetics, 53(9), 1276–1282. 10.1038/s41588-021-00921-z

Wolna, A., Wright, A., Casto, C., Hutchinson, S., Lipkin, B., & Fedorenko, E. (2026). The extended language network: Language-responsive brain areas whose contributions to language remain to be discovered. The Journal of neuroscience : the official journal of the Society for Neuroscience, e0638252026. Advance online publication. 10.1523/JNEUROSCI.0638-25.2026

Wu, J., Ngo, G. H., Greve, D., Li, J., He, T., Fischl, B., Eickhoff, S. B., & Yeo, B. T. T. (2018). Accurate nonlinear mapping between MNI volumetric and FreeSurfer surface coordinate systems. Human brain mapping, 39(9), 3793–3808. 10.1002/hbm.24213

Wu, Q., Chen, J., Yang, X., Zhang, X., He, W., & Xia, J. (2025). Associations of ventriculomegaly and white matter hyperintensities with glymphatic dysfunction in idiopathic normal pressure hydrocephalus. European radiology, 35(7), 4315–4324. 10.1007/s00330-024-11320-3

Xiong, Y., Yu, Q., Zhi, H., Peng, H., Xie, M., Li, R., Li, K., Ma, Y., & Sun, P. (2024). Advances in the study of the glymphatic system and aging. CNS neuroscience & therapeutics, 30(6), e14803. 10.1111/cns.14803

Yamaguchi, A., & Jitsuishi, T. (2024). Structural connectivity of the precuneus and its relation to resting-state networks. Neuroscience Research, 209, 9–17. 10.1016/j.neures.2023.12.004

Yao, J., Sun, X., Chen, Y., Xu, X., Feng, J., Zhang, M., Liu, X., & Shi, X. (2025). Histone deacetylase 6 inhibition attenuates pathological cardiac hypertrophy by promoting autophagy through MAP1LC3B ubiquitination. The Journal of pathology, 266(2), 217–229. 10.1002/path.6419

Yue, G. H., Bilodeau, M., Hardy, P. A., & Enoka, R. M. (1997). Task-dependent effect of limb immobilization on the fatigability of the elbow flexor muscles in humans. Experimental physiology, 82(3), 567–592. 10.1113/expphysiol.1997.sp004048

Zappaterra, M. D., Lisgo, S. N., Lindsay, S., Gygi, S. P., Walsh, C. A., & Ballif, B. A. (2007). A comparative proteomic analysis of human and rat embryonic cerebrospinal fluid. Journal of proteome research, 6(9), 3537–3548. 10.1021/pr070247w

Zeloni, R., Amaolo, A., Morez, A., Zapparoli, E., Akl, Y., Shafie, M., Huerta-Sanchez, E., Pizzagalli, F., Provero, P., Pagani, L., & Marnetto, D. (2026). Effects of introgressed Neanderthal alleles on present-day brain morphology. bioRxiv. 10.64898/2026.04.14.718380

Zhang, A., Bergmans, S., Van Dyck, A., Beckers, A., Moons, L., & Masin, L. (2025). Successful axonal regeneration is associated with intraneuronal metabolic reprogramming. iScience, 28(10), 113631. 10.1016/j.isci.2025.113631

Zhao, B., Li, T., Yang, Y., Wang, X., Luo, T., Shan, Y., Zhu, Z., Xiong, D., Hauberg, M. E., Bendl, J., Fullard, J. F., Roussos, P., Li, Y., Stein, J. L., & Zhu, H. (2021). Common genetic variation influencing human white matter microstructure. Science (New York, N.Y.), 372(6548), eabf3736. 10.1126/science.abf3736

Zhao, B., Li, T., Yang, X., Shu, J., Wang, X., Luo, T., Yang, Y., Wu, Z., Fan, Z., Jiang, Z., Chen, J., Shan, Y., Tang, J., Xiong, D., Zhu, Z., Gao, M., Wyliena, G., Tomlinson, C., Dong, Q., Li, Y., Stein, J., Wang, Y., & Zhu, H. (2022). Genetic influences on the shape of brain ventricular and subcortical structures. MedRXiv. 10.1101/2022.09.26.22279691

Zhao, C., OuYang, Y., Zhang, G., Zang, D., Xia, J., Liang, G., Ye, M., Wang, J., Gan, Y., Zhou, Y., Yang, J., & Li, X. (2025). Association of Glymphatic and White Matter Impairment With the Postoperative Outcome of Pediatric Hydrocephalus. Neurosurgery, 96(1), 193–204. 10.1227/neu.0000000000003050

Zhu, H., Simons, Y. B., Spence, J. P., Sella, G., & Pritchard, J. K. (2026). Genetic architectures of brain-related traits are shaped by strong selective constraints. bioRxiv. 10.64898/2026.03.22.713538

