## Supplementary Figures for "Contextualizing the polygenic basis of human brain shape"

Suppl. Figure 1. *Effect direction relations for each cerebellar component and the average across components with respect to globularity, ventricular morphology, and glymphatic function measures.*

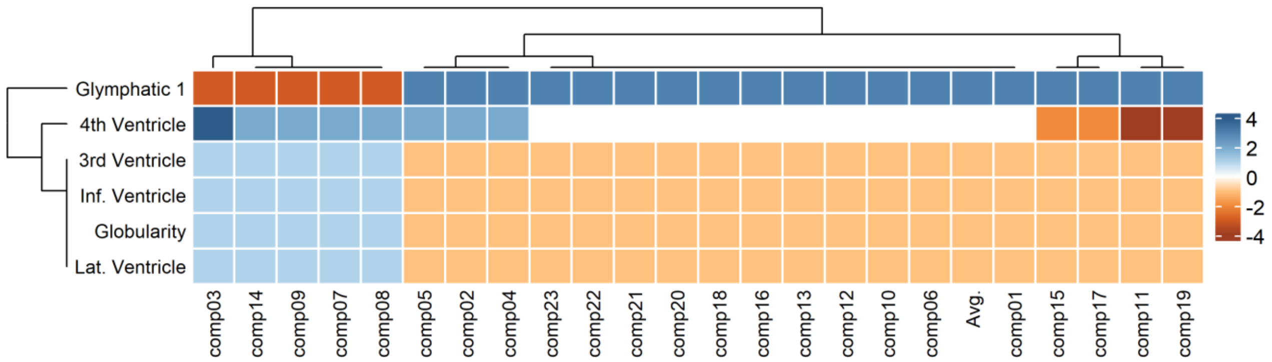


For each cerebellar component, the number of concordant (> 0) and discordant (< 0) effect directions with the brain ventricles, globularity and the glymphatic system are represented. Glymphatic function is again opposed to ventricular morphology and globularity. Besides, a stark contrast is manifested between components 03, 14, 07, 08, 09, fully/mostly concordant with ventricles and globularity, and the rest of components, among which 01, 15, 17, 11 and 19 feature as the most discordant for genome-wide significant overlaps.

Suppl. Figure 2. *Concordance scores for Mekki's (2022) white matter microstructural parameters, by tract.*

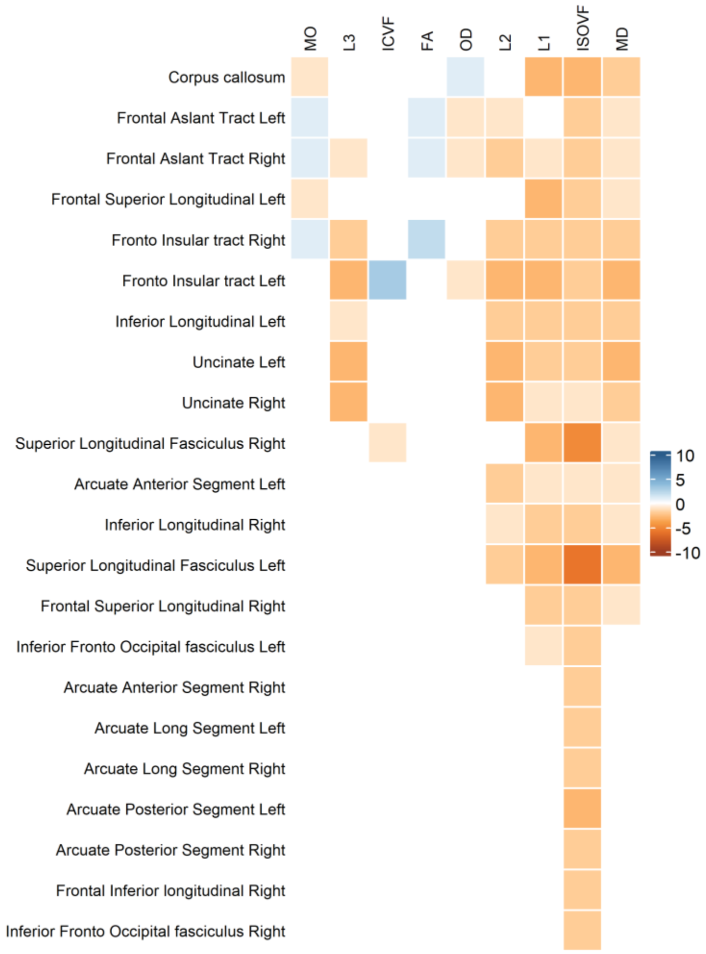


The frontal aslant and fronto-insular tract concentrate concordances for architectural parameters of white matter microstructure (MO, ICVF, FA), whereas the corpus callosum and the left frontal superior longitudinal tract bear exceptional discordances for MO. The corpus callosum is also exceptional in its concordance with OD. As for ISOVF and diffusivity (L1, L2, L3, MD) parameters, widespread discordances are observed across tracts.

Suppl. Figure 3. *Pairwise correlations between traits according to their genes' spatial patterns of expression.*

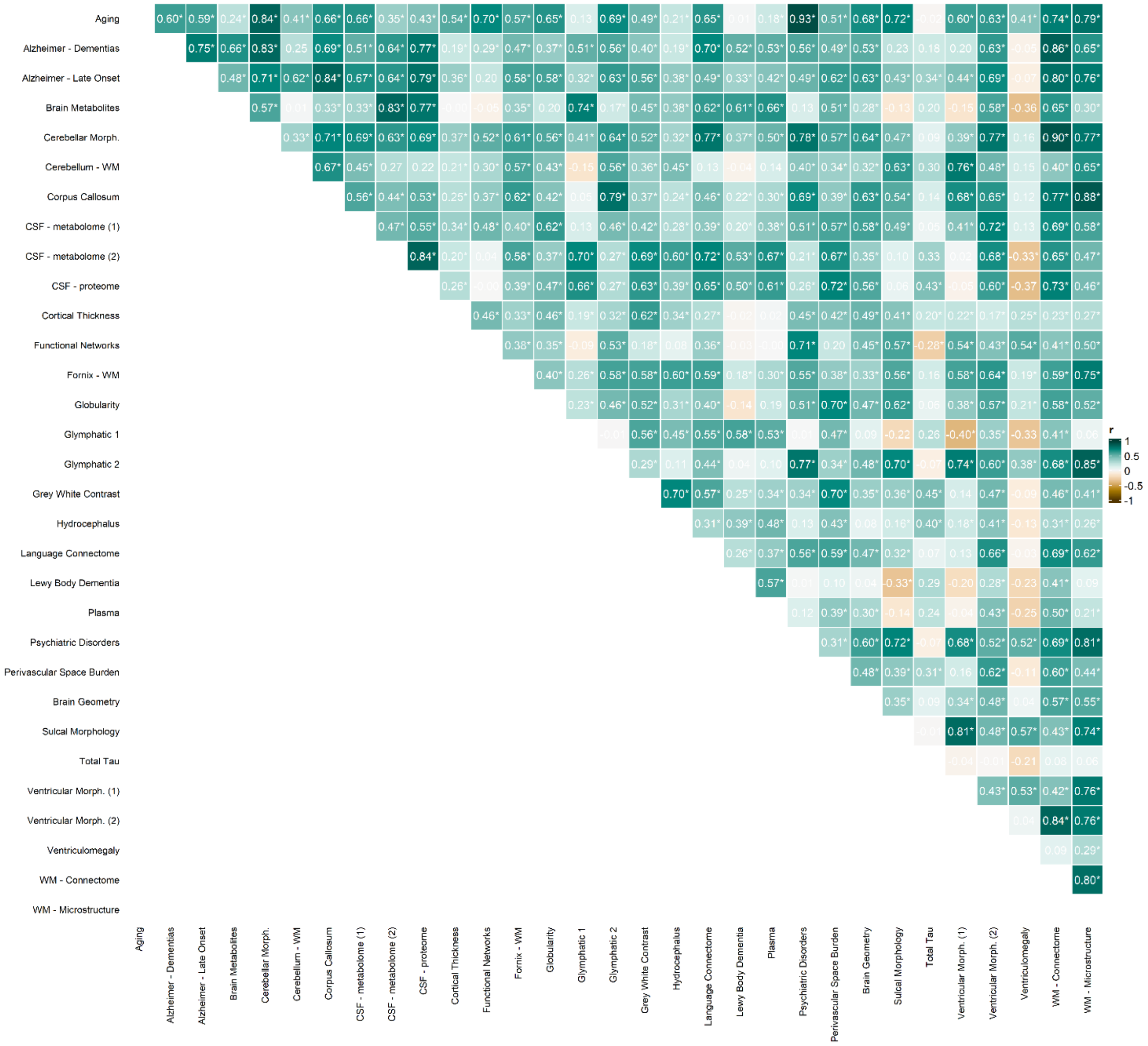


Few negative correlations (brown) are observed, and most of the observed ones concern the size of the ventricles (*ventriculomegaly, ventricular morphology 1)* —for instance, in inverse relations to the glymphatic system (1) or lewy body dementia (2). Despite widespread positive correlations, variation in their strength is also noted. In that sense, globularity's most positive associations with the perivascular space burden, sulcal morphology, the white matter connectome and ventricular morphology (2) may be accentuated.

**Supplementary Figure 4.** *Correlations between our datasets' spatial patterns of gene expression and the sensorimotor association axis components (Sydnor et al., 2021).*

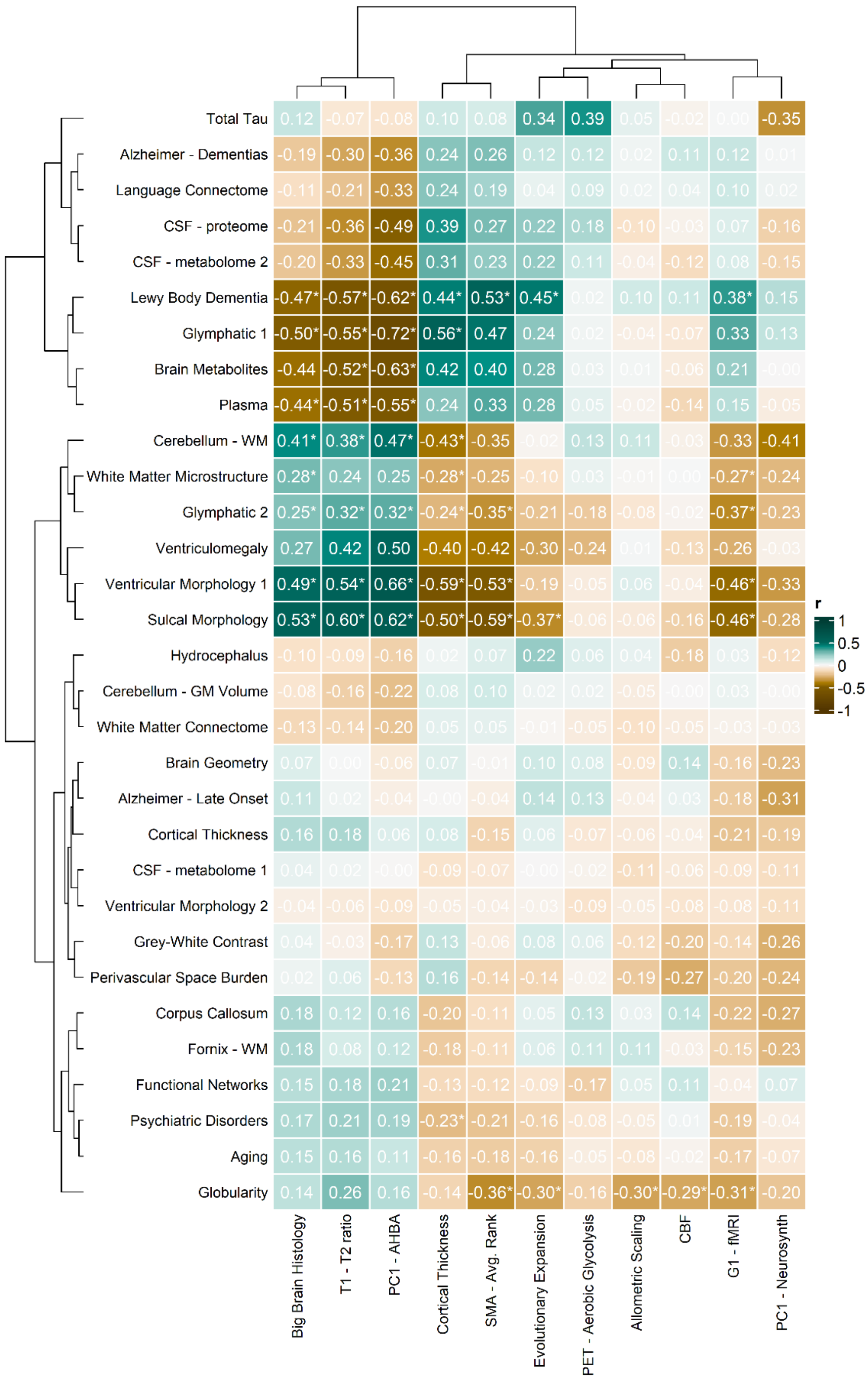


Datasets at the top of the heatmap (especially, from *Alzheimer – Dementias* to *Sulcal Morphology*) show marked patterns of association with the features in the SMA axis, with either an associative signature (*Alzheimer – Dementias* to *Plasma*) or a sensorimotor one (*Cerebellum – WM* to *Sulcal Morphology*). At the bottom of the heatmap, globularity seems to align with the latter group, albeit with milder patterns of correlation to SMA components. Other phenotypes (*Hydrocephalus* to *Aging*) seem to show weaker correlational profiles, with rather insignificant or not fully consistent results.

Suppl. Figure 5. *Datasets significantly correlated with Sydnor et al.'s (2021) sensorimotor association axis components are significantly split into a sensorimotor and an association cluster through multiscale bootstrap resampling with 5000 replicates via pvclust.*

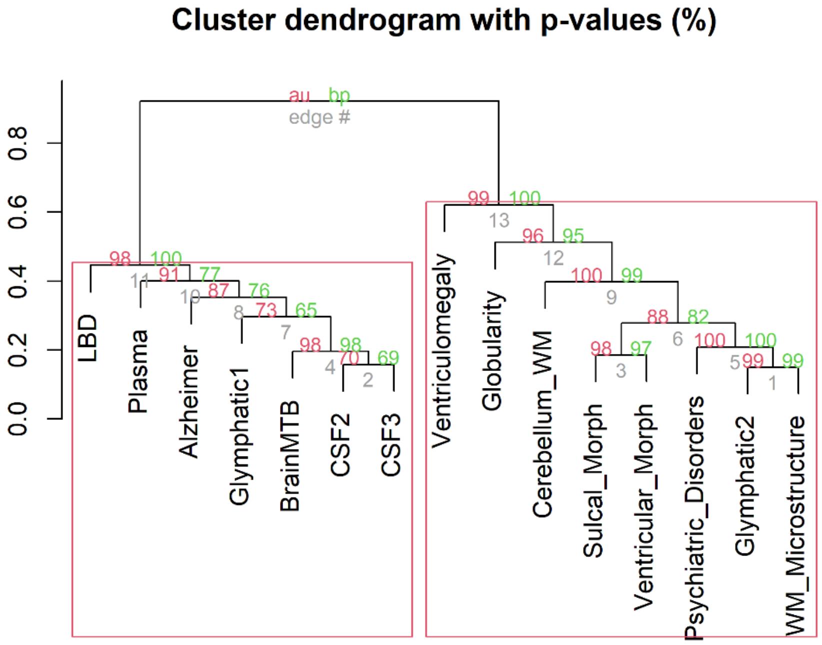


Multiscale bootstrap resampling with 5000 replicates discerns a dissociation between associatively-oriented datasets (LBD, Plasma, Alzheimer, Glymphatic 1, Brain Metabolites, CSF2 (metabolome), CSF3 (proteome); AU = 98%) and sensorimotor-oriented datasets (Ventriculomegaly, Globularity, Cerebellum White Matter, Sulcal Morphology, Ventricular Morphology, Psychiatric Disorders, Glymphatic 2, White Matter Microstructure), given their spatial patterns of expression in relation to Sydnor *et al.*'s (2021) features of cortical organisation.

Suppl. Figure 6. Temporal trajectories of gene expression associated to globularity and brain-related traits.

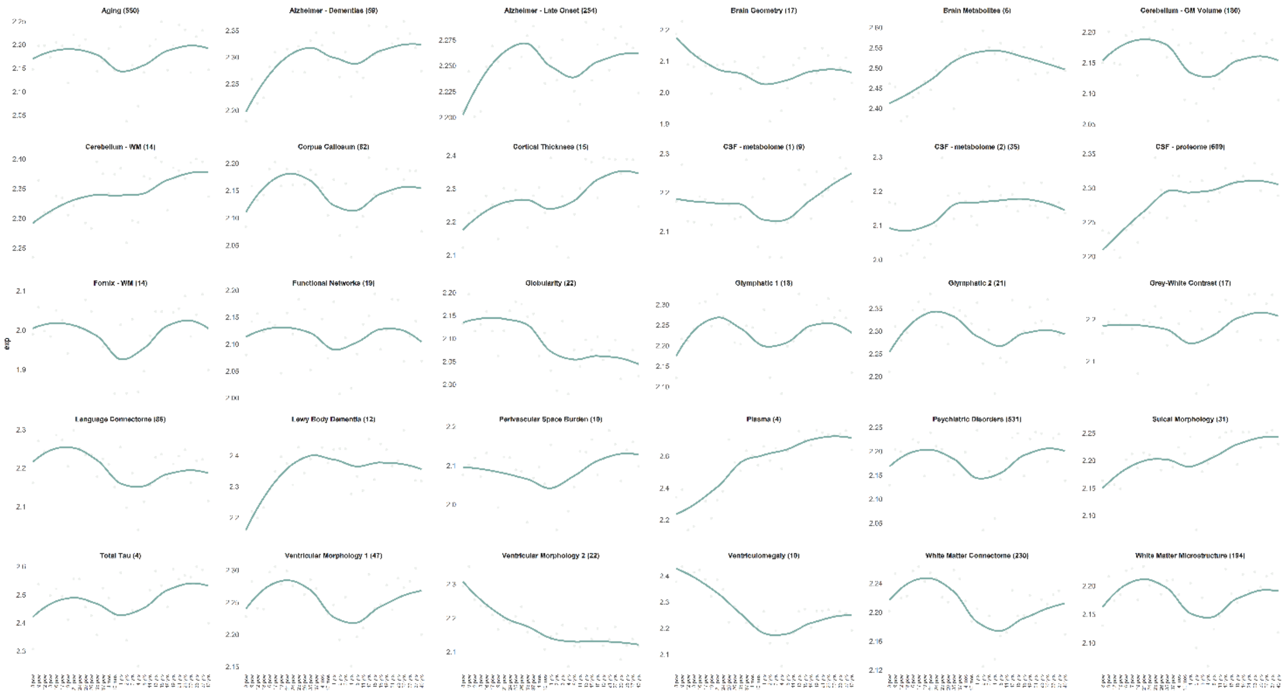


Trajectories of gene expression throughout the lifespan are plotted for each of 30 datasets, based on BrainSpan's original age categories. Whereas some phenotypes are related to increasing transcriptomic trajectories throughout the lifespan (e.g., *Alzheimer – Dementias*), globularity shows a decreasing trajectory from early prenatal peaks, similarly to ventriculomegaly and ventricular morphology (2). Next to each phenotype's name at the corresponding panel, the number of genes from which the trajectory was derived is noted in parentheses.

Suppl. Figure 7. *Complete temporal mapping of the times of emergence associated with variants from Kuhlwilm & Boeckx's (2019) catalog intersecting with our datasets.*

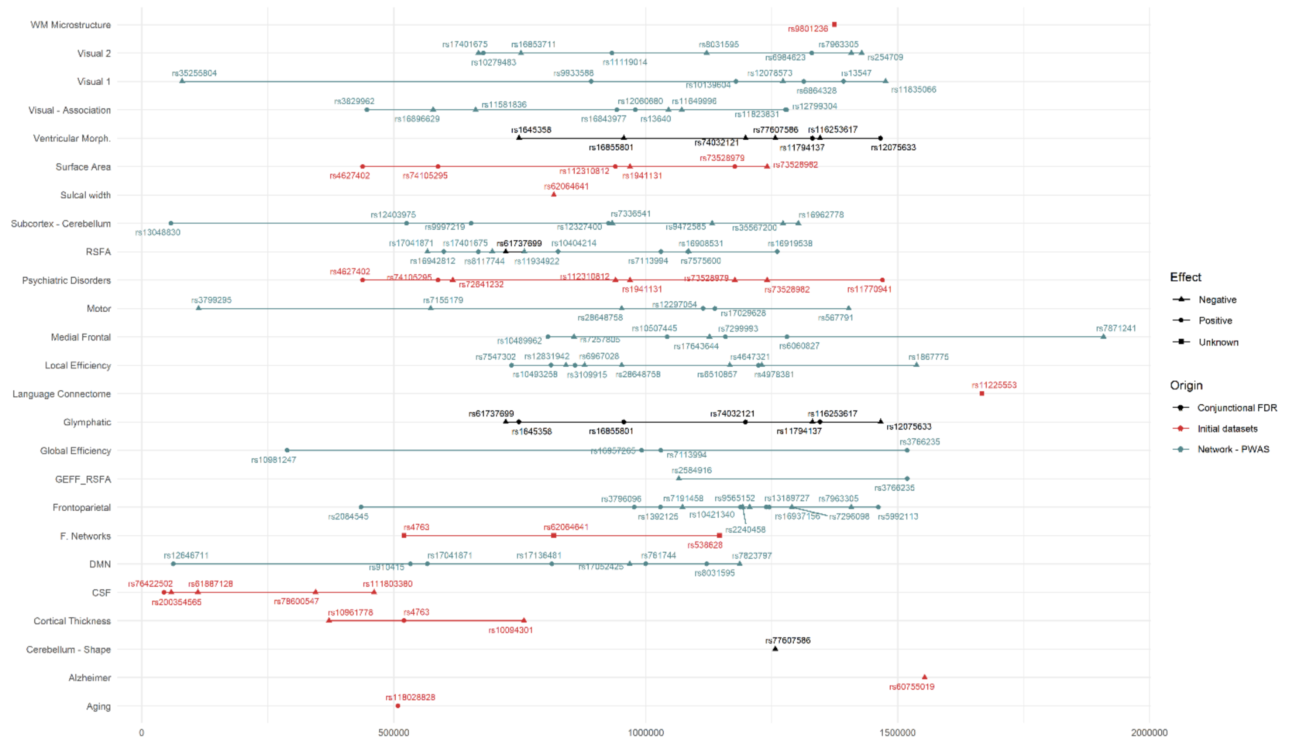


The (x) axis concerns times of emergence, from present (left-most) to 2 Mya (right-most). The (y) axis separates timelines by phenotype. Colours indicate the origin of the datasets represented: our conjunctional FDR analyses, studies from our initial pool (mainly, GWAS), or Bell *et al.*'s (2022) inspection of network measures of brain activity. Shapes indicate the effects of variants' major alleles: positive (circle), negative (triangle) or unknown (square). Recent evolutionary single-nucleotide changes are seen for the CSF (within the last 500,000 years), whereas glymphatic, ventricular, sulcal and functional network phenotypes are generally presented with longer evolutionary histories.

Supplementary Figure 8. *All (concordance/discordance) counts lend consistent results with other measures of convergence and divergence between traits, as reflected by the relations between the glymphatic system's patterns of concordance with sulcal width and Sydnor et al.'s (2021) SMA axis features.*

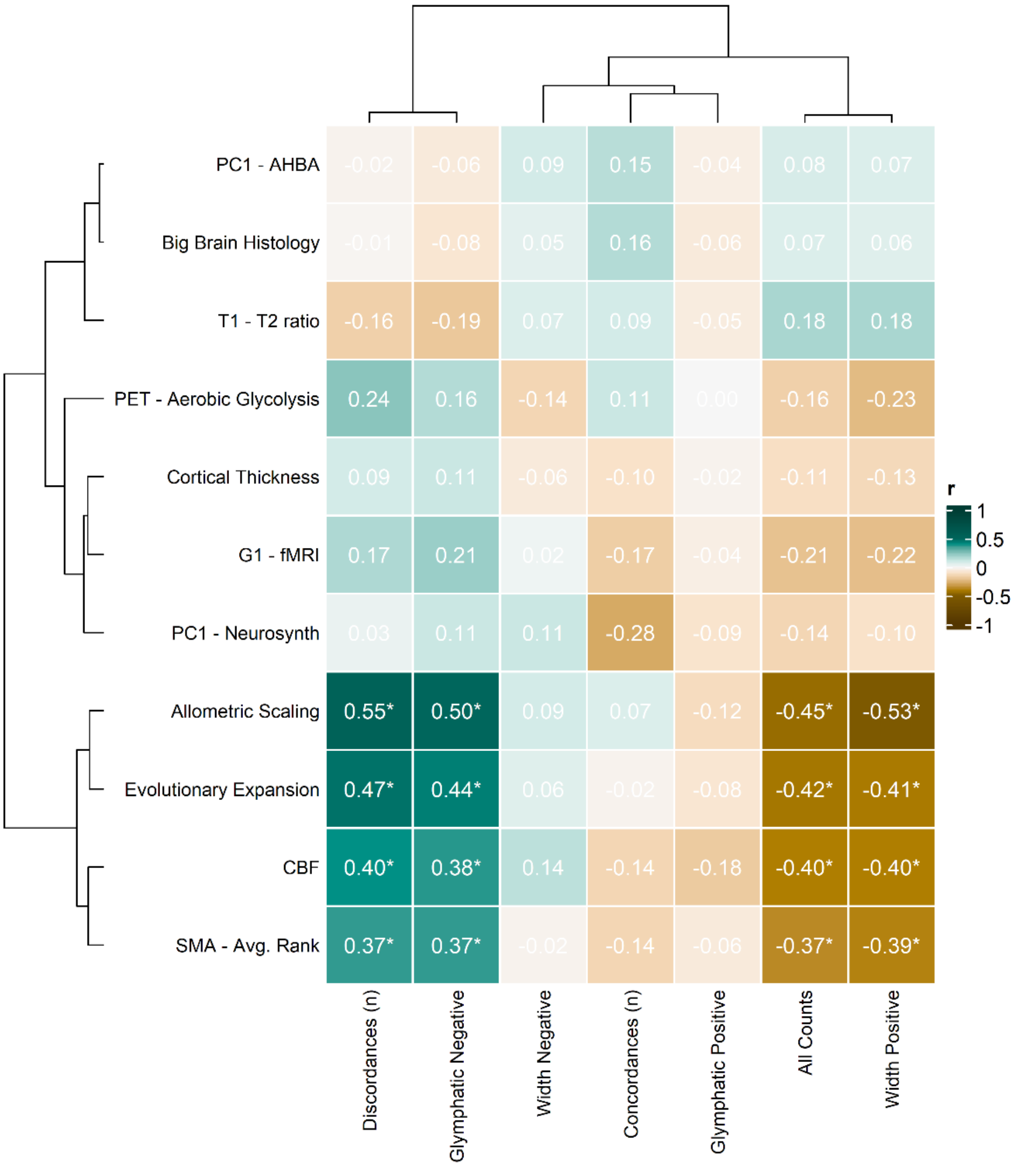


*All Counts* reflects our integrative measure of concordant and discordant variants (*n concordant – n discordant*). For the glymphatic system, given its overall discordance with sulcal width, the score is generally negative, and correlates negatively with associative features of the SMA axis (e.g., *allometric scaling*). Consistent with this, variants with positive effects on sulcal width (*Width Positive*) generally show negative effects on the glymphatic system, and these negative effects again inversely correlate with associative features of the SMA axis. There are few variants with positive effects on the glymphatic system (*Glymphatic Positive*) and negative effects on sulcal width (*Width Negative*), which therefore yields weak correlations. Similarly, counts of concordant variants do not achieve a strong signature. Instead, discordant variant counts (*Discordant (n)*, where each discordant SNP = +1) yield an inverse signature with respect to our *All Counts* and *Width Positive* parameters, with positive correlations with associative features of the SMA axis. This is expected, as here discordances are counted positively, whereas they are computed negatively in *All Counts*. Finally, variants with negative effects on the glymphatic system (*Glymphatic Negative*) tend to bear positive effects on sulcal width, which in turn are positively correlated with associative features of the SMA axis. Overall, the consistency of results achieved for our *All Counts* measure with respect to other approaches suggests it can be a functional parameter to account for both concordant and discordant dimensions of pleiotropy.
